# Imaging membrane damage in ferroptosis and necrosis by wash-free fluorogenic chemical probes

**DOI:** 10.1101/2023.06.02.543437

**Authors:** Philipp Mauker, Daniela Beckmann, Annabel Kitowski, Constanze Heise, Chantal Wientjens, Andrew J. Davidson, Will Wood, Christoph Wilhelm, Julia Thorn-Seshold, Thomas Misgeld, Martin Kerschensteiner, Oliver Thorn-Seshold

## Abstract

Selectively labelling cells with damaged membranes is needed in contexts as simple as identifying dead cells in culture, or as complex as imaging membrane barrier functionality *in vivo*. The commonly used dyes are permanently coloured/fluorescent dyes that are simply excluded by intact membranes, but to achieve good image contrast therefore requires removing their extracellular signal by washing or background subtraction, which are not possible *in vivo*. Here, we develop fluorogenic probes which sensitively and selectively reveal damaged cells, without needing washing steps since their fluorescence turns on from near-zero background. From a set of novel fluorogenic probes impermeabilised by sulfonations along different vectors, we identify a specific disulfonated fluorogenic scaffold that enters cells only upon membrane damage, where it is enzymatically activated to mark them. The esterase probe **iPS-FS**_**2**_ is a reliable tool to reveal live cells that have been permeabilised by biological, biochemical, or physical membrane damage; and it can be used in multicolour microscopy. We confirm the modularity of this approach by also adapting it for redox-unmasked cell-excluded probes with improved hydrolytic stability. This scaffold-based design thus provides tools for wash-free *in vivo* imaging of membrane damage, which is relevant across many pathologies. The insightss gained from these probes should also be translatable to damage-targeted prodrugs, for selective therapy of membrane-compromised cells.

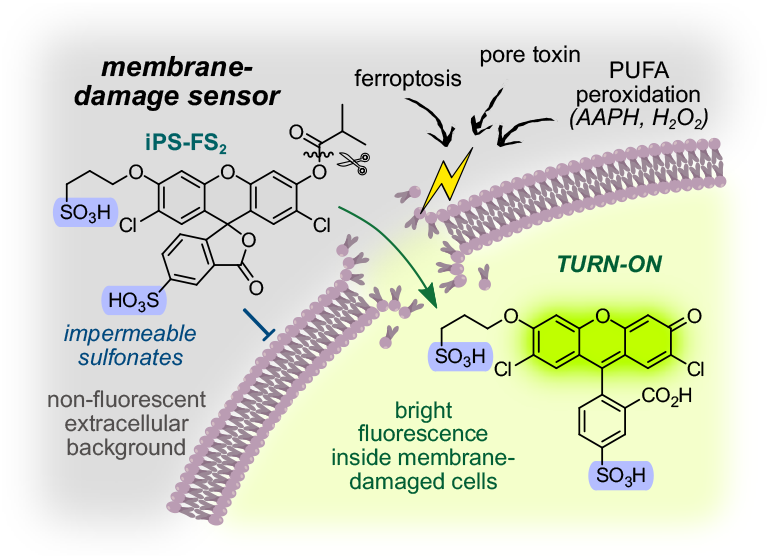

## INTRODUCTION

Cells are enclosed by the plasma membrane, which retains cellular components such as ions and proteins, and reduces exposure to extra-cellular molecules. Eukaryotic membranes are primarily bilayers of amphipathic membrane lipids that expose hydrophilic headgroups to water while aggregating their lipophilic tails.^1^ While small apolar molecules readily cross membranes, larger or more polar species such as ions and proteins are “membrane impermeable”, because the energy penalty for desolvation and traversal through the lipophilic inner region of the membrane is high: i.e. they cannot cross membranes by passive diffusion, which is the mode of cellular entry that we focus on in this work.^2^

Loss of cell membrane integrity impairs the separation of intracellular and extracellular spaces, allowing otherwise impermeable molecules to cross membranes.^3,4^ This results in celluar stress and can initiate cell death pathways, due to the entry or mis-localisation of toxic species, aberrant signalling, or osmotic and energetic overload. Cell membrane integrity can be impaired by a variety of physiological and pathological processes: from physical stress (e.g., mechanical injury) to chemical modification (e.g., lipid peroxidation) or protein pore formation (e.g., induced by pyroptosis or bacterial toxins). Repair of membranes can potentially reverse cellular demise, which is particularly important in postmitotic cells such as neurons e.g. after traumatic or inflammatory insults in conditions such as blunt spinal trauma or multiple sclerosis. This duality renders membrane-damaged cells an interesting study population (see **Supporting Notes 1-2**).^5,6^ As such, finding ways to selectively address cell membrane integrity with small molecules is crucial: either for detecting membrane-compromised cells on the verge of death, or to therapeutically reroute them towards survival. Here we develop chemistry to do this, by using cells’ compromised membranes as a selective, passive entry pathway (we will refer to these compromised cells as “damaged” or “leaky”).

Charge-based membrane impermeabilisation of small molecules is often achieved by attaching “permanently” charged groups such as sulfonates (pK_a_ ≈ −2).^7^ Charged imaging agents are routinely used *in vitro* for detecting leaky cell membranes, e.g. for discriminating live and dead cells. For example, Trypan Blue is a polysulfonated dye used for counting dead cells, as it selectively passes their leaky membranes but is excluded from healthy cells.^8^ The cationic fluorophore propidium iodide is used similarly for staining dead cells in fluorescence microscopy and flow cytometry (**Fig 1a**).^9^ Conceptually however, the current agents have major caveats: (1) “Always-on” chromophores/fluorophores (such as Trypan Blue) require washing steps, background subtraction, and/or cell isolation to remove the majority of the dye which is not taken up in cells. (2) Environment-sensitive stains (like propidium iodide) feature intracellular signal turn-on by DNA binding, so they can be highly toxic. While this is not problematic for live/dead stainings, such agents are not useful for long-term tracking of live cells with permeabilised membranes (see **Supporting Note 1**). Both limitations are particularly severe for *in vivo* imaging. Rationally exploiting membrane (im)permeabilisation can also offer far more powerful applications than live/dead assays. Currently, charged bioactive molecules are being used to selectively address extracellular targets: sulfonating inhibitors to restrict them to extracellularly-exposed receptors^10^, using impermeable photocages to deliver signaling lipids locally at the plasma membrane^11^, or selective bioorthogonal labelling of cell-surface proteins by cell-excluded tags^12,13^. Yet, while these methods are elegant, there is an unmet need for generalised approaches to simultaneously mask and impermeabilise molecules, such that only intracellular reactions in permeabilised cells activate them: ideally giving *“impermeable, off → ON”* fluorogenic probes (or prodrugs).

**Figure 1:**
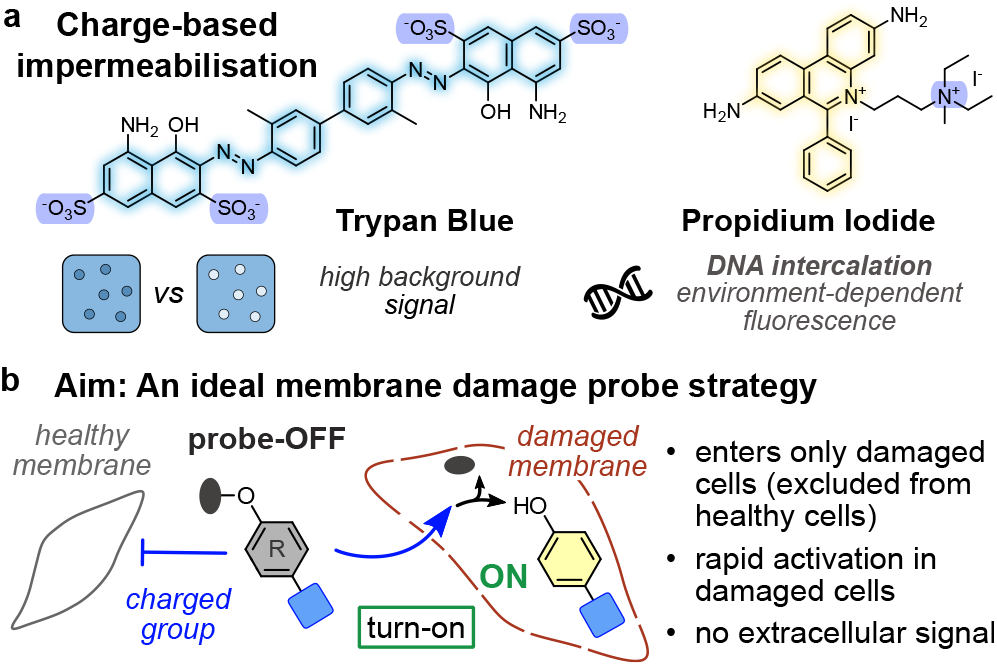
Cell-impermeable probes. **(a)** Current imaging agents for damaged membranes. **(b)** Goal for a damage-selective fluorogenic probe.

Fluorogenic probes are ideally nonfluorescent compounds that only develop fluorescence after activation by a target trigger. This off→ON mode could solve the problems that block current always-on membrane damage dyes from *in vivo* uses, since it eliminates nonspecific background and so maximises their sensitivity (signal to background) without requiring washout. Indeed, fluorogenic probes have become crucial tools to image and quantify biological processes noninvasively in living cells, particularly for investigating enzyme activities of peptidases, esterases, phosphatases, glycosidases, and oxidoreductases.^14–20^ Cell-permeable fluorogenic probes that become fluorescent and cell-trapped after entry are highly valued and widely used (e.g. fluorogenic acetoxymethyl ethers, such as calcein-AM^21^). However, until now, there have been no modular chemical systems that are silenced and cell *impermeable*, but likewise activate an imaging agent or release a drug upon entry.

Here, we aimed at developing fluorogenic membrane damage probes (**Fig 1b**), that use general chemical or biological features to meet these performance needs: (1) near-zero background fluorescence before activation, and no activation outside of cells, to enable wash-/subtraction-free imaging; (2) exclusion from cells with intact membranes; (3) entry into membrane-damaged cells; (4) rapid signal generation upon entry, by enzymatic unmasking of a modular capping group, and effective cellular retention; (5) bright, tunable fluorescence with narrow absorption and emission spectra matching typical biological imaging settings, to enable co-staining with other fluorophores.

We chose xanthenes as the fluorogenic scaffold. Xanthenes can adopt two forms: a completely non-fluorescent “closed” form where the π-conjugation of the xanthene is interrupted by spirocyclisation, or a conjugated, fluorescent, quinoid “open” form. The closed form can be trapped by chemical capping, resulting in a non-fluorescent probe. Ideally, this can be unmasked rapidly and selectively by target enzymes, allowing equilibration with the open form and so generating fluorescence. Fluorescein ester probes for example can be intracellularly activated in many cell types.^22^ Although fluorescein acetates are famously susceptible to spontaneous hydrolysis, other esters can resist spontaneous hydrolysis (discussion below);^23^ and a range of more stable masking groups with other enzymatic targets are known (need 1). Xanthene fluorescence is also bright, narrow, easily tuned by substitutions, and fluorescein is compatible with standard microscopes and multiplexed imaging (need 5).

We explored polysulfonatation for cell exclusion. Tanaka *et al*. had reported the first cell-excluded fluorescein probe, 5-sulfofluorescein diacetate, in 1995.^24^ Raines and colleagues’ work on improving the aqueous instability of fluorescein acetates^23^ instead suggested isobutyrates as esters, to balance speed of activation in damaged cells against acceptably low spontaneous hydrolysis rates, while harnessing n→π* donation in the 2’s,7’-dichlorofluorescein (**H**_**2**_**-FS**_**0**_) scaffold to maximise hydrolytic resistance.^23^ Taken together, our starting point was to test the selectivity of anionically decorated 5-sulfo-2’,7’-dichlorofluorescein esters as membrane-impermeable nonfluorescent probes, that should enter damaged cells and be activated therein; with a longer-term goal of applying similar probes or prodrugs for disease detection or modification *in vivo* (**Supporting Note 2**).^5,6^

## 2. RESULTS AND DISCUSSION

### 2.1 Doubly capped monosulfonate probes enter healthy cells

We synthesised the novel, sulfonated (charged) fluorophore 5-sulfo-2’
s,7’-dichlorofluorescein (**H**_**2**_**-FS**_**1**_) by adapting published procedures (for probe nomenclature see **Fig S1**).^25^ Condensation of 4-chlororesorcinol and 4-sulfophthalic acid gave a 68% yield of the mixed 5-sand 6-sulfofluoresceins. Conveniently, this could be purified to >95% purity of the 5-sulfo regioisomer by precipitation and wash/filter steps, without chromatography of the very polar mixture. The absorption, excitation and emission properties of **H**_**2**_**-FS**_**1**_ (ex/em maxima ca. 500/525 nm) match the strong fluorophore dichlorofluorescein **H**_**2**_**-FS**_**0**_ (**Fig S2a-c**).

We synthesised the fluorogenic probes **i**_**2**_**-FS**_**1**_ and **a**_**2**_**-FS**_**1**_ (**Fig 2a**) by double *O-*acylation of **H**_**2**_**-FS**_**1**_ using 5-10 eq. of acid anhydride in DMF (see **Supporting Information**). As expected, these were locked as the nonfluorescent spirolactones (**Fig S2d-e**).

**Figure 2:**
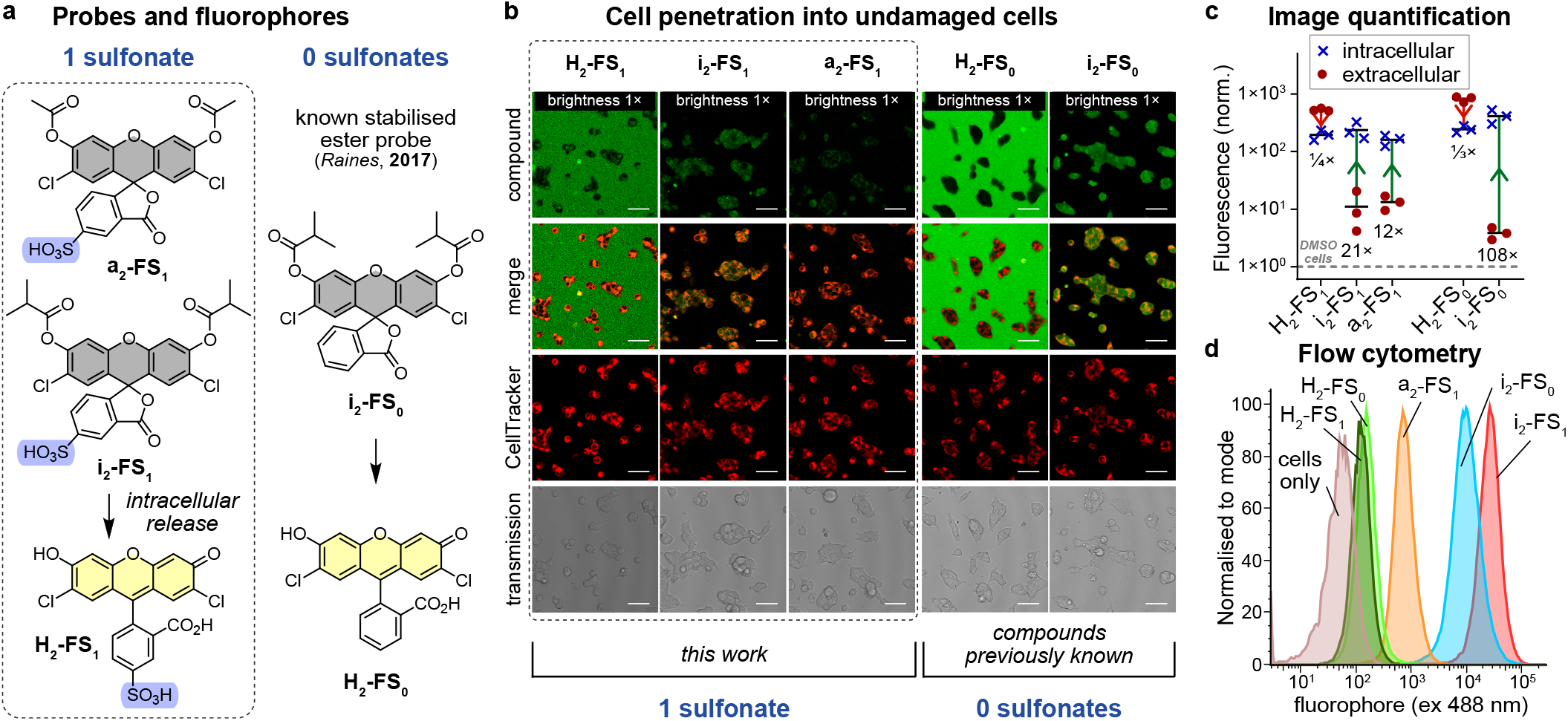
Cell penetration and fluorogenicity of double-capped probes or fluorophores, with healthy cells. **(a)** Sulfonated probes **i2-FS**_**1**_ and **a**_**2**_**-FS**_**1**_ from fluorophore **H**_**2**_**-FS**_**1**_; and known reference **i**_**2**_**-FS**_**0**_ from **H**_**2**_**-FS**_**0**_. **(b-c)** Confocal microscopy of healthy HEK cells incubated with test compounds (5 µM) for 10 min without washing (CellTracker is a cytosolic stain; scale bars: 50 µm). **(c)** Image quantification for intracellular vs extracellular fluorescence (note log_10_ vertical axis; values normalised to autofluorescence of DMSO control as 1). **(d)** Orthogonal quantification of cellular entry-and-activation from flow cytometry on healthy HeLa cells (10 µM test compounds, 20 min incubation; all data: *n* = 3). (“Brightness 1×” is the adjustment relative to settings used in **Fig. 4**; full legend in **Supporting Information**).

To perform wash-free damage imaging, the probes must remain non-fluorescent outside cells while avoiding spontaneous nonenzymatic hydrolysis, while also being rapidly enzymatically processed upon cellular entry to give fluorescence. We first tested probe stability. During a typical timeframe for cell experiments (15 min incubation at 37 °C), the iso-butyrates of **i**_**2**_**-FS**_**1**_ were relatively stable in PBS or HBSS buffers (<1% activation), while the less sterically hindered acetates of **a**_**2**_**-FS**_**1**_ were ca. 8-fold more labile as expected (see **Supporting Information at Fig S2g-h** for discussion). These observations emphasise the need for isobutyrate capping. We then assessed probe activation by the model enzyme porcine liver esterase (PLE; **Fig S2f**) showing that both probes were activated above their spontaneous hydrolysis rates (**Fig S2f**).

We then tested the exclusion of the probes from healthy cells, using confocal microscopy to localise their activated fluorescence and quantify the intracellular vs. extracellular intensities (**Fig 2b**,**c**). We anticipated the probes **i**_**2**_**-FS**_**1**_ and **a**_**2**_**-FS**_**1**_, and the fluorophore **H**_**2**_**-FS**_**1**_, would be cellexcluded due to their sulfonation. We also used neutral **H**_**2**_**-FS**_**0**_ as a slowly cell-entering reference fluorophore, and Raines’ neutral **i**_**2**_**-FS**_**0**_ ^23^ as a reference probe with good cell penetration and enzymatic deacylation. Predictably, fluorophores **H**_**2**_**-FS**_**0**_ and **H**_**2**_**-FS**_**1**_ gave the highest extracellular signals (red circles in **Fig 2c**), with lower intracellular signal (blue crosses in **Fig 2c**), so cells were seen as “shadow images” in microscopy (**Fig 2b**). We observed slow hydrolytic increase of extracellular fluorescence for **i**_**2**_**-FS**_**1**_ (faster for **a**_**2**_**-FS**_**1**_), towards a maximum value for full uncapping set by control **H**_**2**_**-FS**_**1**_ as expected. This supports that monosulfonate **H**_**2**_**-FS**_**1**_ is membrane-impermeable, as we had desired to see also for its ester-capped probes.

Against our hopes, sulfonated **i**_**2**_**-FS**_**1**_ gave almost equal intracellular fluorescence as **i**_**2**_**-FS**_**0**_: i.e. when doubly *O-*acylated to the fluorogenic spirolactone, a single sulfonate is no longer sufficient for cell exclusion (**Fig 2b**,**c**). **a**_**2**_**-FS**_**1**_ showed a lower intracellular signal than **i**_**2**_**-FS**_**1**_, supporting that isobutyrate lipophilicity promotes cellular entry despite the sulfonate charge penalty.^26^ Thus, we needed to develop even more hydrophilic probes to enforce cell exclusion. Still, before starting this, we noted two useful results:

Firstly, isobutyrate ester capping on the sulfo-chlorofluorescein is an excellent platform for low-background, high-sensitivity imaging as needed for damaged-cell probes. The extracellular background signal from *O-*deacylation of diester **i**_**2**_**-FS**_**1**_ reached only 1-2% of the signal of uncapped **H**_**2**_**-FS**_**1**_, without using any washing steps before imaging (red circles, **Fig 2c**). Secondly, the microscopy quantifications in HEK cells matched qualitatively to population averages from flow cytometry over thousands of HeLa cells (**Fig 2d, Fig S3**). Thus, the effect that two lipophilic *O-*capping groups can bring a monosulfonated spirocyclised probe across the membrane^27^ for intracellular activation is conserved across different cell types. The flow cytometry data are also monomodal, supporting that a single mechanism is responsible for entry – a result that is encouraging for further tuning.

### 2.2 A capped disulfonate probe is excluded by healthy cells

To tune membrane permeability via increasingly polar *O-*derivatised spiro probes, while keeping the stability of the isobutyrate cap strategy, we alkylated one of the xanthene phenols with a more polar moiety before installing the ester: resulting in mono-capped probes (**Fig 3a**). Mono-capping also simplifies data analysis, since a single ester cleavage gives full signal (with double capping, the first and second cleavages contribute differently to sample fluorescence.)

**Figure 3:**
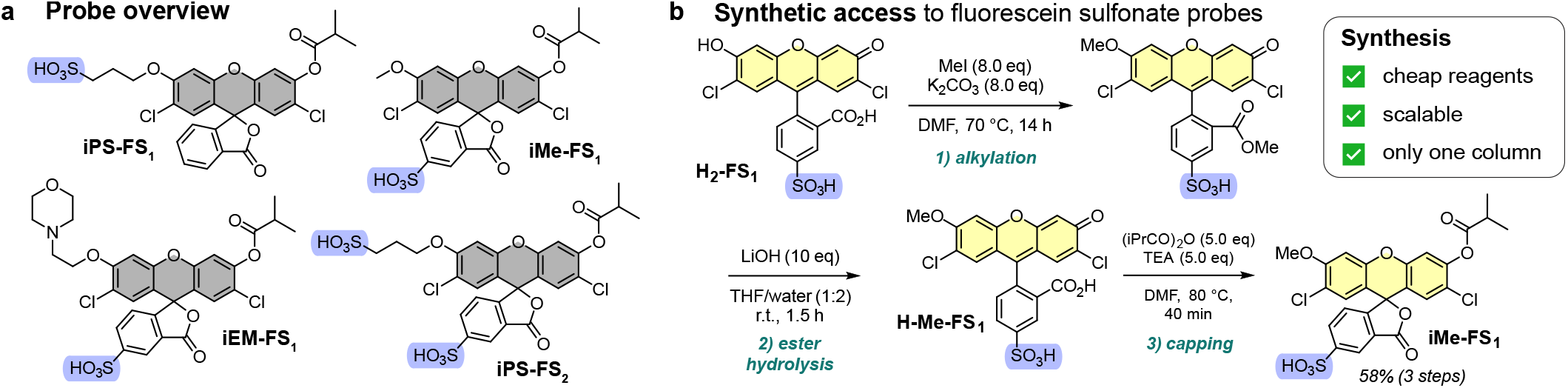
Mono-capped probes. **(a-b)** Overview and typical synthetic route for mono-capped mono/bis-anionic fluorescein probes.

As polar *O-*alkyl moieties for the **H**_**2**_**-FS**_**1**_ scaffold, we used methyl (**iMe-FS**_**1**_), amine (β-ethylmorpholino **iEM-FS**_**1**_), or sulfonate (γ-propylsulfonate **iPS-FS**_**2**_). To test the role of sulfonate count vs. orientation, we compared **iMe-FS**_**1**_ with **iPS-FS**_**1**_, whose single sulfonate is on the *O-*alkyl chain, not the pendant ring (**Fig 3a**). The probe syntheses largely avoided chromatography (**Fig 3b** and **Supporting Information**). E.g., for the synthesis of **iMe-FS**_**1**_, **H**_**2**_**-FS**_**1**_ was doubly alkylated at both the carboxylate and the phenol, then the ester was cleaved mildly with LiOH to return the mono-alkylated fluorophores (**H-Me-FS**_**1**_), which were then capped with isobutyric anhydride to afford the probes in good yield (58% over three steps, **Fig 3b**).

The optical properties of the mono-alkylated fluorescein cores were slightly different from e.g. non-alkylated **H**_**2**_**-FS**_**1**_ (**Fig S2a**,**b**): they have two absorption maxima (ca. 460 nm and 485 nm), with 4-fold weaker absorbance; and their fluorescence emission (λ_max_ ca. 525 nm) is ca. half as intense, which are expected results^28^ for the less symmetric chromophore (due to this intensity difference, their microscopy images will be displayed at different brightness levels, to keep focus on the key feature that is their degree of exclusion from healthy cells relative to uptake in damaged cells). All probes and fluorophores were photostable (**Fig S2c**), and all probes were stable enough in lyophilisation and handling to ensure <5% ester cleavage in stocks (**Fig S2d**,**e**). Matching our expectations, the signal from the mono-capped ester probes was enzymatically activated 5–10× faster to its maximum than the doubly capped probes (**Fig S2f**). The hydrolytic stability of the ester probes was good in PBS, limited in HBSS, and poor in DMEM medium (**Fig S2g**,**h**).

The probes’ entry into healthy cells aligns with their polarity, e.g. permeable **iPS-FS**_**1**_ (23-fold higher intracellular than extracellular signal), vs poorly uptaken **iMe-FS**_**1**_ / **iEM-FS**_**1**_ (<3-fold). Disulfonated **iPS-FS**_**2**_ seemed to be our best probe candidate, with nearly no intracellular fluorescence in healthy cells relative to background (**Fig 4a**,**b**). Matching expectations, the fluorophore cores of these sulfonated probes were also cell-excluded (**Fig S4**). These microscopy results were matched on a population level by flow cytometry assessments (**Fig S3c**,**d**).

**Figure 4:**
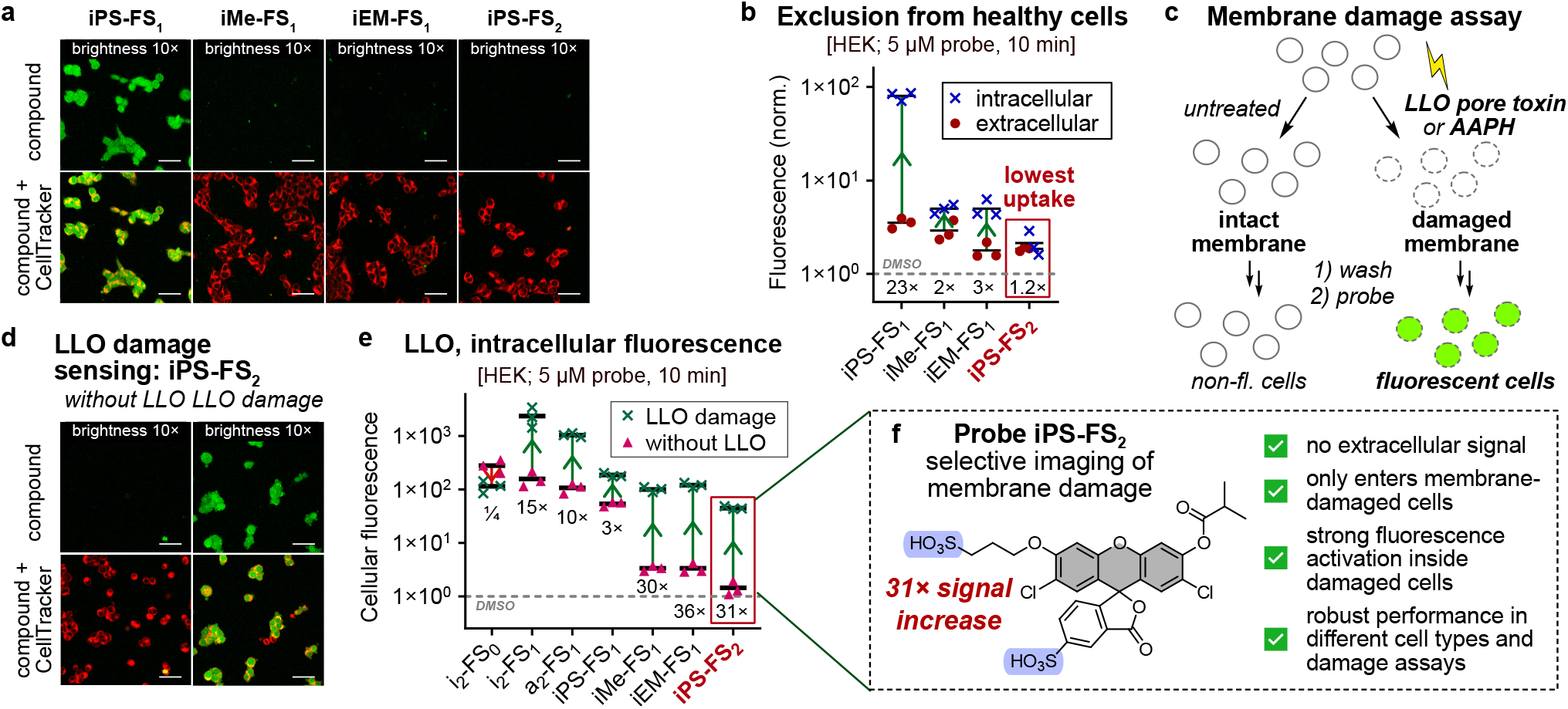
Fluorogenic probes that are excluded from healthy cells but enter and turn on inside damaged cells. **(a-b)** Healthy HEK cells treated with probes (5 µM, 10 min), quantified for intracellular and extracellular fluorescence relative to DMSO control (set to 1). **(c)** Scheme of membrane damage assays. **(d-e)** Listeriolysin (LLO) damage assay (HEK cells, 5 µM probe for 10 min, no washing). Quantification of intracellular fluorescence (relative to DMSO autofluorescence) for damaged vs. undamaged cells. **(f)** Damage probe **iPS-FS**_**2**_. (Scale bars 50 µm; “Brightness 10×” is the adjustment relative to settings used in **Fig. 2**; full legend in **Supporting Information**).

### 2.3 Selective entry of the mono-capped disulfonate probe across damaged cell membranes

To investigate probe uptake and activation in damaged cells, we first used the pore-forming bacterial protein toxin listeriolysin O (LLO) to induce membrane damage (**Fig 4c**).^29^ We chose LLO as a model as it induces small-diameter pores that are suitable for small molecule uptake,^30^ by a well-studied mechanism of action.^31^ Microscopy showed that all sulfonated probes had higher intracellular signal with LLO than without: but only **iPS-FS**_**2**_ proved suitable as a wash-free probe, since it is the only probe with both low extracellular background *and* the high ratio of signal in LLO-damaged vs healthy cells (>30×) needed for selective and wash-free imaging of damaged cells (**Fig 4d-f, Fig S5**; see **Supporting Information** at **Fig S4-6** for discussion of other probe types). Thus, **iPS-FS**_**2**_ became our best probe for charge-based discrimination of membrane damage.

We next explored whether **iPS-FS**_**2**_ would also report on radical damage to membrane integrity. Among the many biological instances of membrane damage, we were interested in targeting membrane-damaged axons as are found in inflammatory lesions in models of multiple sclerosis.^5,6^ Although the cause of axonal membrane permeabilisation in neuroinflammatory lesions is unknown, such lesions feature high loads of reactive oxygen and nitrogen species^32^ and it is speculated that permeabilisation may result from plasma membrane lipid peroxidation (see **Supporting Note 2**). We therefore treated the neuronal cell line PC12^33^ with radical-generating initiator 2,2’-azobis(2-amidinopropane) (**AAPH**) to peroxidise membrane lipids.^34^ Again, **iPS-FS**_**2**_ was mostly excluded from healthy cells, yet stained the AAPH-damaged cells with the highest selectivity (8-fold higher fluorescence, **Fig 5a**,**b**; see **Supporting Information** at **Fig S7** for further discussion), thus identifying this fluorogenic disulfonate as a reliable membrane damage probe.

**Figure 5:**
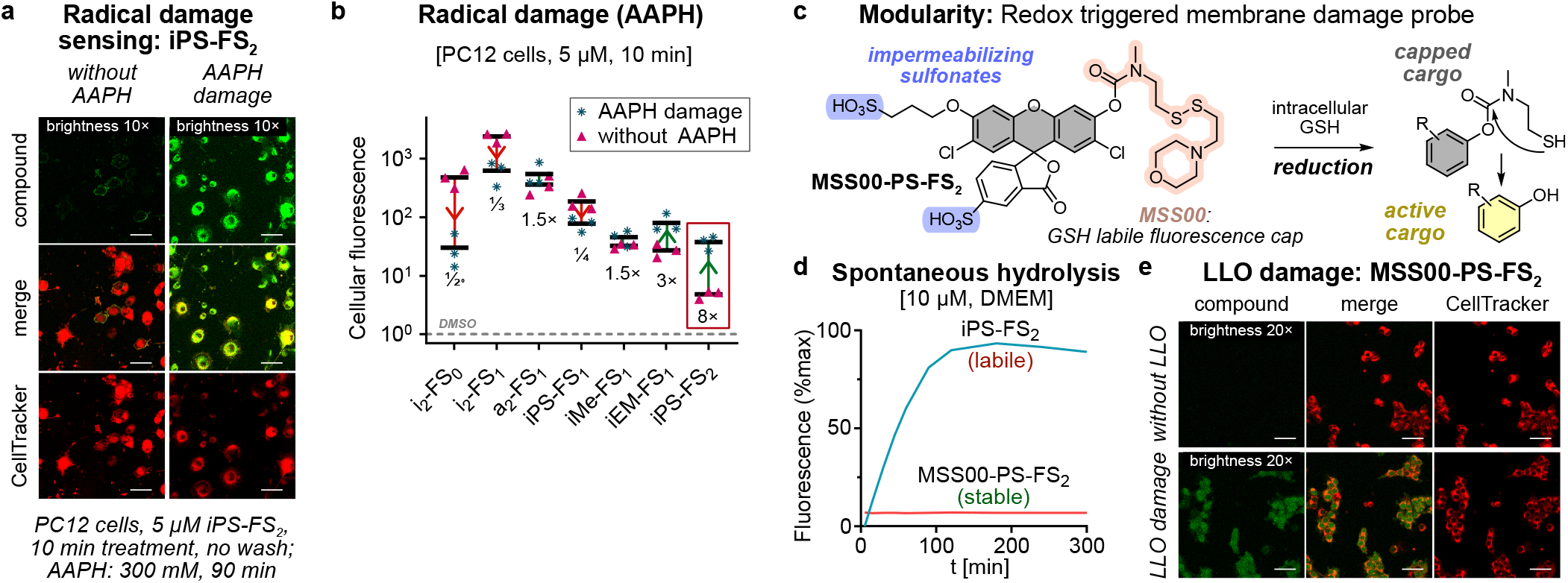
Biological and chemical scope of fluorogenic cell-excluded probes. **(a-b)** Nonspecific membrane damage (PUFA peroxidation) by radical initiator AAPH. Images and quantified intracellular fluorescence from probes (5 µM, 10 min, no wash). **(c-e)** Probe **MSS00-PS-FS**_**2**_, with a GSH-labile reduction trigger attached as a phenolic carbamate, has far less spontaneous probe hydrolysis in cell culture medium, but retains the damage-selective performance in the LLO assay (HEK cells, 5 µM probe for 10 min, no wash, scale bar: 50 µm; “Brightness 10/20×” is the adjustment relative to settings used in **Fig. 2,4**; full legend in **Supporting Information**).

### 2.4 Modularity of the cell-excluded fluorogenic probe design

The design of this probe system is intentionally modular, in that the impermeabilisation is ensured by the fluorophore, so leaving a flexible choice of trigger. We aim at alternative fluorogenic triggers for two reasons: (1) to identify more extracellularly-robust triggers that could withstand standard cell culture media over longer imaging timecourses, and might even resist extracellular processing *in vivo*, as more performant and applicable damage imaging probes; (2) to work towards molecular imaging either of specific cell-surface enzymes on healthy cells, or of specific enzymes at work in damaged cells.

In this study we explored one approach for extracellularly robust probes, using an intracellularly reducible disulfide trigger^35^ that adapts onto the fluorogenic scaffold as the tertiary carbamate probe **MSS00-PS-FS**_**2**_. This ought to be far more hydrolytically resistant than the ester of **iPS-FS**_**2**_, yet also effectively cleaved by esterases upon cellular entry (**Fig 5c**). Indeed, in cell-free tests of probe stability in DMEM media, no undesired **MSS00-PS-FS**_**2**_ activation was observed over five hours (**Fig 5d, Fig S10a**), but even 0.1 mM GSH rapidly activated it (**Fig S10b**; intracellular GSH is typically 1-5 mM^36^). **MSS00-PS-FS**_**2**_ was as well-excluded from healthy cells as **iPS-FS**_**2**_ (**Fig 5e, Fig S11**), and likewise showed good signal increase in damaged cells (8-fold in LLO assay, 3-fold in AAPH assay, **Fig S11**). Although its intracellular signal intensities after standard 10 min incubation were ca. 3 times lower than with ester probe **iPS-FS**_**2**_, this could be overcome by taking advantage of the stability of **MSS00-PS-FS**_**2**_ to apply higher concentrations for longer, if needed for sufficient signal intensity (although in our model systems we observe strong enough fluorescence). Taken together, we find the mono-capped disulfonate fluorogenic design to be a synthetically accessible, flexible, and reliable platform for small molecule probes that are excluded from healthy cells.

### 2.5 Comparison to size-based cell exclusion

An alternative approach for fluorogenic probes to selectively report on cell membrane damage might be to develop macromolecular probes that are excluded from healthy cells on the basis of their size (rather than their charge), yet passively enter more porous, damaged cells. While the focus of this work was on small molecule probes, we briefly tested the accessibility of this approach (full discussion in the **Supporting Information**, section **Dextrans**); though it should be noted that active macromolecular uptake by healthy cells occurs by varied mechanisms and to significant degrees, which can prevent complete size-based exclusion from healthy cells - for more information see e.g. ref^37^.

In brief, we first tested commercially available permanently fluorescent dextrans in the size range 6-2000 kDa in an AAPH assay and observed slightly increased uptake after damage (**Fig S9**). To test this approach for turn-on probes, we then required no-wash fluorogenic dextrans; however, these are barely reported (first monofunctional fluorogenic dextran only published in 2018^37^) and are not commercially available. We therefore prepared mono-isobutyrate-capped, chloro-stabilised fluorogenic fluorescein NHS ester **NHS-i-Flu**, guessing that its one-step off/on unmasking would make it more suitable for quantification than the previous double-capped^37^ fluorogen. Fina Biosolutions LLC (MD, USA) conjugated this fluorogen to 70 kDa dextran, however, carefully controlled AAPH assays did not indicate sufficient damage-dependent cellular entry (**Fig S9**). Thus, we did not continue these investigations, assuming that the background rates of active uptake in healthy cells are too high, and any change in passive membrane permeability after damage is too small, for macromolecular carriers to be a robust approach to probe membrane integrity.

### 2.6 Membrane biology 1: cellular imaging of ferroptosis

Ferroptosis is a non-apoptotic form of cell death first described by Stockwell in 2012,^39^ where radical chain reactions with (poly)unsaturated fatty acids in membrane lipids, mediated by molecular oxygen, form lipid hydroperoxides that lead to catastrophic loss of membrane integrity and to cell death. Several cellular mechanisms suppress ferroptosis. These include the reductase GPx4 that reduces reactive lipid hydroperoxides to unreactive alcohols to stop propagation;^3,40^ thus, GPx4 inhibitors such as RSL3 are useful as ferroptosis inducers. Chemical methods such as H2O_2_ overload also induce lipid peroxidation by initiating Fenton chemistry, albeit intervening at a different stage of the cascade^41^.

Typically, ferroptosis is imaged by proxy: using reactive membrane-integrating probes designed to intercept oxygen-centred radicals (e.g. **BODIPY-C11**^42^, **Fig 6f**) or to react with hydroperoxide products (e.g. phosphine “Liperfluo”). However, neither method directly reveals the biologically relevant result of ferroptosis, i.e. the breakdown of membrane integrity; and both probes are also open to criticism for their invasive nature, since they alter the peroxidation biochemistry involved.^42^ Furthermore, **BODIPY-C11** requires ratiometric measurement that blocks two imaging channels.

**Figure 6:**
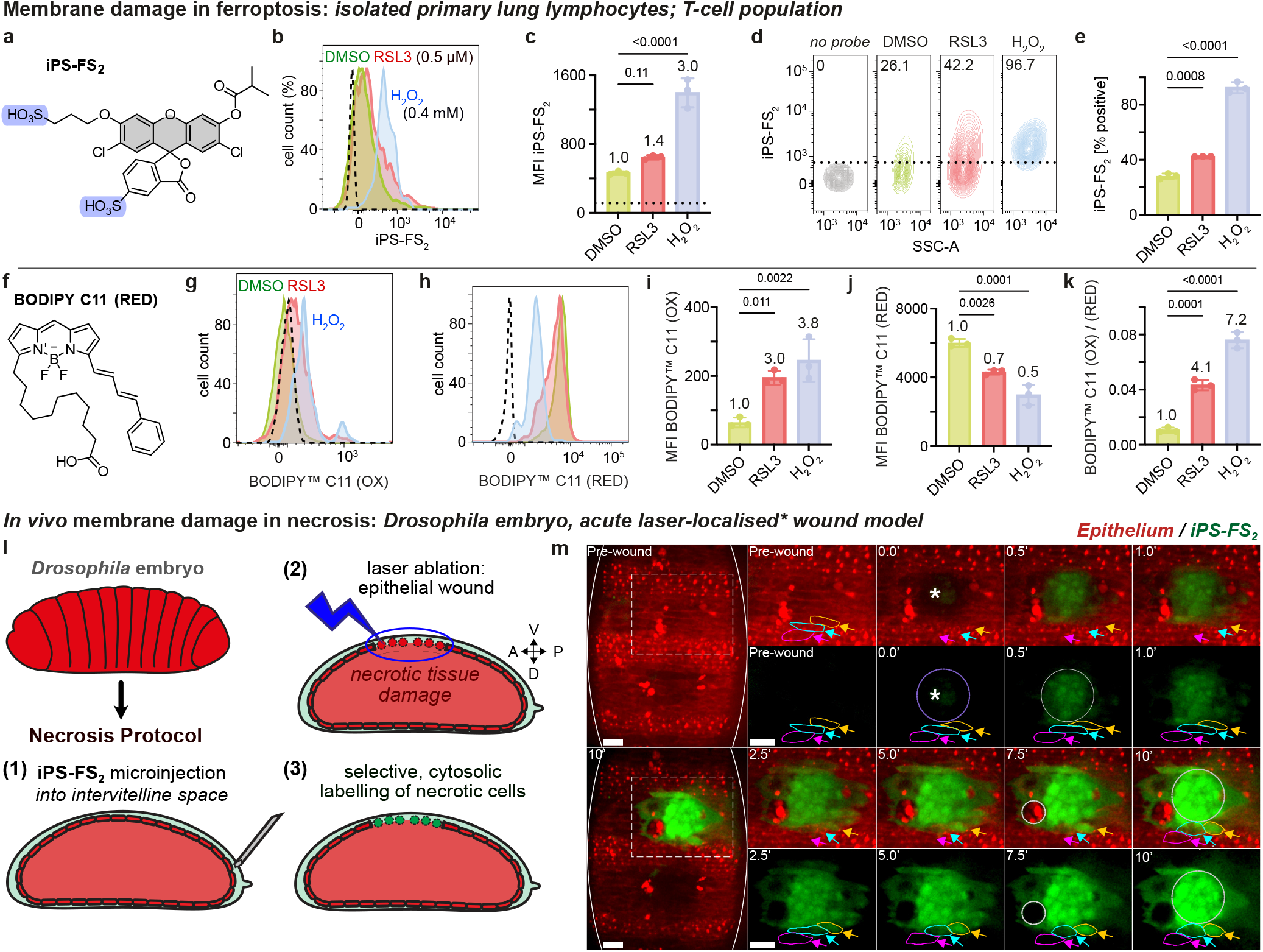
Sensing membrane damage in ferroptosis and necrosis. **(a-k)** T-cells were isolated from a culture of total lung lymphocytes, optionally pre-treated with RSL3 or H_2_O_2_, then treated for 1 h with probes **iPS-FS**_**2**_ (50 μM) or **BODIPY-C11** (RED) (250 μM), and analysed by flow cytometry. **(a-e)** Cells pre-treated with H_2_O_2_ are strongly marked by **iPS-FS**_**2**_, while control cells are not **(b**,**d). (f-k) BODIPY-C11** also reveals cells pre-treated with H_2_O_2_, although it needs ratiometric evaluation of the oxidised/reduced fluorescence intensities to do so **(k). (l-m)** After treating live *Drosophila* embryo with **iPS-FS**_**2**_, local necrotic damage was triggered by local laser wounding (at asterisk) of the ventral epithelium (mcherry-Moesin cell surface marker in red), and probe fluorescence was imaged (green). Cells in the ablation damage focus (violet ring at time 0.0’) rapidly take up and activate the probe (time 0.5’); most cells that contact the damage zone are also labelled over time (e.g. yellow and cyan-indicated epithelial cells), but neighbouring undamaged cells remain dark (e.g. magenta-indicated epithelial cell, or macrophage indicated by grey ring at time 7.5’). (Time in minutes, scale bar 5 µm; embryo cartoon adapted from ref^38^; full legend in **Supporting Information**; higher resolution imaging in **Fig S12** and **Movie S1**).

We hypothesised that the redox-independent probe **iPS-FS**_**2**_ could instead directly report the degree of membrane integrity loss during ferroptosis, providing an alternative strategy for ferroptosis imaging. To this end, we challenged T-cells isolated from mouse lungs with RSL3 or H_2_O_2_ to induce ferroptotic lipid peroxidation, then probed them with **iPS-FS**_**2**_. Flow cytometry showed strong challenge-dependent signal enhancement (**Fig 6a-e**), that corresponded well to a **BODIPY-C11** proxy readout of peroxidation (**Fig 6f-k**). Although such a match is not strictly needed, since these probes measure different biological/biochemical aspects, the similarity of their responses is highly satisfying.

### 2.7 Membrane biology 2: *in vivo* imaging of necrosis

The requirements for a probe to succeed *in vivo*, i.e. in a live intact animal (3D system), are much more stringent than in 2D cell culture. In culture, all cells contact a vast reservoir of solution where convective currents supply fresh probe while diluting away any fluorophore that may be activated and/or released from a cell, which can result in fluorescence appearing more cell-localised than is the underlying chemistry. In 3D however, both supply and dilution are limited; so in order for a probe to label damaged cells robustly in a *cell-resolved* manner, probe entry, activation, and retention must be efficient and exceptionally selective for damaged cells, while exclusion from healthy cells as well as avoiding activation in the extracellular space must both be ensured. We now used **iPS-FS**_**2**_ in a demanding *in vivo* cell damage assay to test its resolution.

Acute necrotic injury can be induced *in vivo* with high spatial precision by laser ablation; and *Drosophila* embryos are a well-studied and ethically acceptable model organism in which to probe it. Traditional laser damage assays with live *Drosophila* embryos microinject cell-impermeable chemical dyes into the intervitelline space (a fluid layer surrounding the epithelium)^43^, then image their fluorescence signal following laser ablation (**Fig 6l**), typically observing the accumulation of environment-dependent DNA-binding fluorophores such as Sytox or Draq7 in the nuclei of necrotic cells^44^. We performed similar assays using **iPS-FS**_**2**_, expecting that it would instead reveal the entire cytosolic volume of the necrotic cell areas. Compared to nuclear-only staining, this would be a strong advantage when determining the boundaries of necrotic zones across stacks of 2D image slices, which do not necessarily each map the nucleus together with accessible cell outline markers.

Time-lapse microscopy of embryos pre-treated with **iPS-FS**_**2**_ before local laser ablation revealed outstanding performance (**Fig 6m**). Cells in the focus of the laser damage (visible since their cell outline marker^45^ is photobleached by the laser, at time 0) were durably labelled by **iPS-FS**_**2**_ fluorescence within just 30 s (time 0.5, **Fig 6m**). Cells contacting the damage focus were also labelled, at a predictably slower rate, which is consistent with the hypothesis that localised loss of membrane integrity even at one side or tip of the cell can be detected by **iPS-FS**_**2**_ entry/activation (yellow/cyan-indicated cells, time 5.0, **Fig 6m**). Pleasingly, viable epithelial cells or macrophages neighbouring the damaged cells were left completely nonfluorescent, even appearing as “shadow images” (magenta-indicated epithelial cell at time 5.0, white-indicated macrophage at time 7.5, **Fig 6m**). Against this sharply spatially defined pool of fluorescent damaged cells, all other (non-damaged) cells and areas remained nonfluorescent throughout the imaging experiment, supporting that **iPS-FS**_**2**_ is a highly sensitive as well as selective probe for robustly, fluorogenically imaging necrosis *in vivo* (further detail in **Fig S12, Supporting Note 3**, and **Movie S1**).

Taken together, we believe that direct, noninvasive reagents such as **iPS-FS**_**2**_, which selectively reveal the membrane permeabilisation that is a driving force not only in ferroptosis, necrosis, bacterial toxicity, or axonal degeneration, but also in a host of other pathological situations, may find broad applications across biophysics and biology.

## 3. CONCLUSIONS

We have harnessed molecular charges and polarity in a modular fluorogenic probe design, to develop a reliable platform for probing cell membrane integrity. We aimed for “no-wash” molecular imaging probes that are excluded from healthy cells, yet enter cells with compromised membranes, whereupon their fluorescence is activated from near-zero background. The resulting disulfonated probes, such as the ester probe **iPS-FS**_**2**_ or the reducible probe **MSS00-PS-FS**_**2**_, promise applications in non-invasive tracking and quantification of the induction of membrane permeability or the recovery of membrane integrity, across a range of biological or chemical stressors or diseases.

Several avenues for development are open. In particular, the extracellularly-stable tertiary carbamate probe offers an intriguing lead for no-wash *in vivo* imaging of membrane damage, with obvious chemical avenues for improvement by e.g. (a) tuning the fluorophore toward red/NIR operation by adaptation e.g. to a Si-fluorescein^46^; and (b) improving the intracellular turn-on speed set by the trigger^47^. Wider biological exploration of the disease model scope where membrane permeability probes usefully report on biology is another productive avenue for development. The opportunity to use cell-excluded probes on healthy cells to perform molecular imaging of their cell-surface enzymology is a biochemical avenue that will be fascinating to explore, particularly in a redox context^48^. Finally, the opportunities to harness similar chemical design principles to selectively deliver pharmacologically active agents instead of fluorophores, that could ameliorate disease states characterised by increased membrane permeability, are alluring.

Conceptually, we suspect that sets of chemical probes such as the ones devised here can reveal rich lessons for the broader chemical biology community. The 3D orientation of the sulfonates of **iPS-FS**_**2**_, far from each other along orthogonal vectors, was an active design choice based on observations that chemically simpler disulfonates with a simple overall head/tail structure caused blebbing or membranolytic stress, which we attributed to surfactant-like effects (results being prepared for separate publication). A flip side to that observation, as pointed out by a colleague, is that our disulfonate probes may themselves *induce* mild membrane stress which only integrity-compromised cells are unable to resist. That would make their design far more actively involved in achieving their biological readout, than a picture of “N charges per M aromatic rings” would suggest: giving many opportunities for nuanced research.

With many avenues for chemical, biochemical, biological and biophysical explorations now opening up, we look forward to further investigation and applications of these deceptively simple chemical tools, as we move towards a deeper awareness of the nuances hidden behind that humblest of models: the simple lipid bilayer membrane.

## ASSOCIATED CONTENT

### Supporting Information

The Supporting Information is available free of charge on the ACS Publications website.

Synthesis, analysis, biochemical and cell biological evaluations (PDF) Movie S1: Timelapse imaging of fly embryo laser damage, corresponding to **Figure 6** (AVI)

## AUTHOR INFORMATION

### Author Contributions

P.M. performed synthesis, chemical analysis, enzymatic cell-free studies, cell biology, flow cytometry, coordinated data assembly. D.B. performed cell biology, confocal microscopy and image analysis / quantification. A.K. and C.H. performed cell biology and flow cytometry. C. Wientjens performed cell biology for ferroptosis sensing. A.D. performed necrosis imaging in *Drosophila*. A.K., J.T.-S., C. Wilhelm, T.M. and M.K. supervised cell biology. W.W. performed necrosis imaging in Drosophila. T.M. and M.K. supervised confocal microscopy. T.M., M.K. and O.T.-S. designed the concept and experiments. O.T.-S. supervised all other experiments and wrote the manuscript with input of the other authors. Correspondence and requests for materials should be addressed to O.T.-S.

## ACKNOWLEDGMENT

We thank Dr. Andrew Lees (Fina Biosolutions LLC, MD, USA) for many helpful discussions while planning and designing dextran conjugates. We thank Dr. Lees and Dr. Samson Gebretnsae (Fina Biosolutions LLC, MD, USA) for performing all custom syntheses of dextran conjugates. We thank Alexander Sailer and Adrian Müller-Deku (LMU) for initial work on fluorescein diacetate NHS ester conjugations to aminodextrans; Bekkah Bingham (LMU) for chromatographic analysis of the integrity of fluorescein diacetate in dextran conjugates; and Philip Woelfle (LMU) for initial testing of dextran conjugates. We are grateful to Henrietta Lacks, now deceased, and to her surviving family members for their contributions to biomedical research.

