## Supplementary Information for "Imaging membrane damage in ferroptosis and necrosis by wash-free fluorogenic chemical probes"

4.6 Necrosis imaging in the fly ... 19
5 Biological Materials and Methods ... 20
5.1 Cell lines ... 20
5.2 Cell-free characterisation ... 20
5.3 Flow cytometry ... 21
5.4 Confocal microscopy ... 21
5.4.1 LLO and AAPH damage assay procedures ... 22
5.4.2 Image analysis ... 22
5.5 Ferroptosis sensing, isolated mouse lung lymphocytes ... 22
5.6 Necrosis sensing, live fly embryo ... 23
6 Chemistry materials and methods ... 24
6.1 Analytical methods and techniques ... 24
6.1.1 Analytical methods ... 24
6.1.2 Synthetic techniques ... 24
6.2 Synthetic procedures ... 25
6.2.1 Literature procedures ... 25
6.2.2 Symmetric Fluorophores and Probes [non- or bis-acylated] ... 25
6.2.3 General Procedure for mono-capped probes and fluorophores ... 26
6.2.4 7'-*O*-alkylated-2'-*O*-acylated [mono-capped] fluorogenic probes ... 27
6.2.5 2'-*O*-[mono]alkylated fluorophores ... 28
6.2.6 Reducible disulfide probe ... 30
6.2.7 Esterase-labile fluorogenic probe for attachment onto macromolecules ... 31
7 Raw Biological Assay Data ... 32
8 Flow cytometry (undamaged HeLa cells): Gating and controls ... 33
9 NMR Spectra ... 35

### 1 Overview of all fluorogenic probes and fluorophores

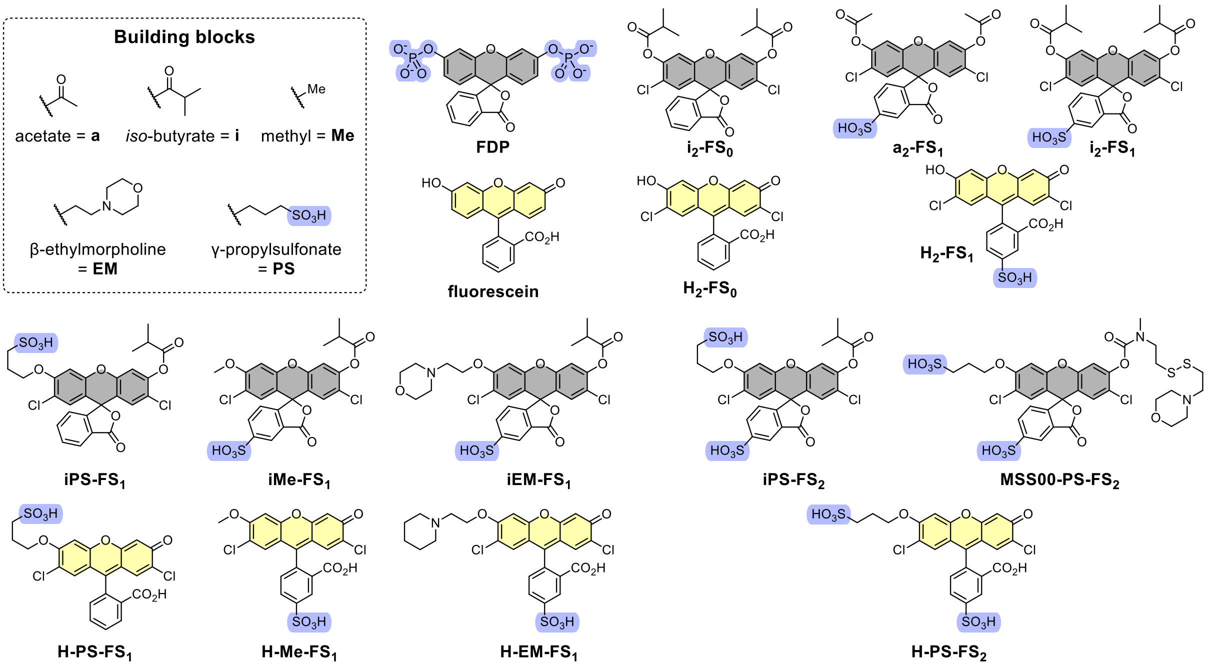

**Fig S1**: Structure overview of all probes and fluorophores and their naming rationale.

### 2 Photocharacterisation and cell free stability / esterase characterisation

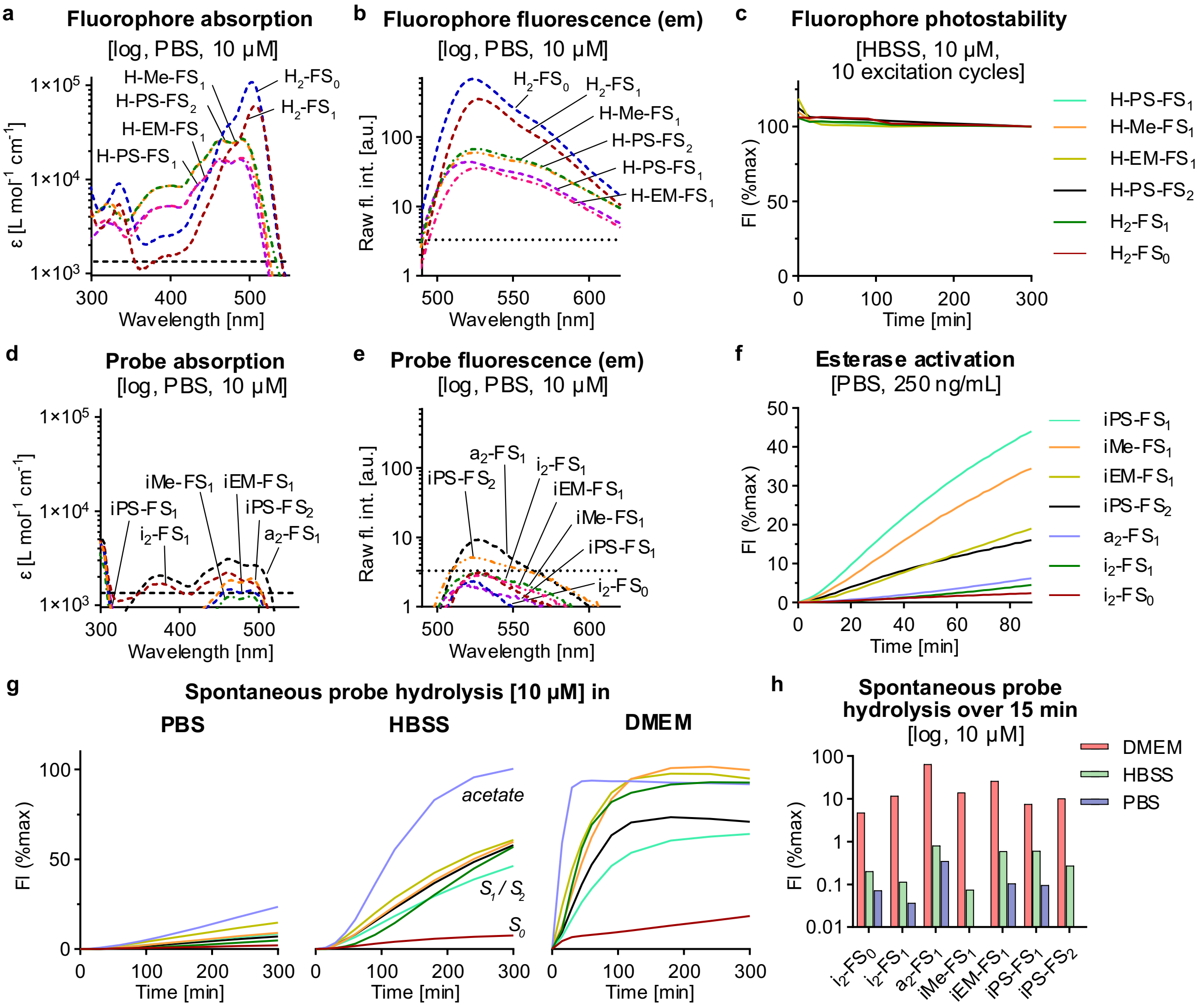

**Fig S2**: **Photocharacterisation, probe stability towards hydrolysis and *in vitro* activation by esterase (a)** UV-vis absorption spectra of fluorophores (10 µM in PBS); **(b)** fluorescence emission spectra (excitation: 485 nm) of the fluorophores (10 µM in PBS); **(c)** photostability of the fluorophores during plate-reader imaging (10 acquisitions over 6 h). **(d)** UV-vis absorption spectra of probes (10 µM in PBS); **(e)** fluorescence emission spectra (excitation: 485 nm) of the probes (10 µM in PBS); **(f)** time-course of probe activation by porcine liver esterase (250 ng/mL, probe conc.: 10 µM in PBS). **(g)** spontaneous probe hydrolysis time-courses in DMEM, PBS and HBSS (probe conc.: 10 µM); **(h)** summary of spontaneous probe hydrolysis after 15 min in DMEM, HBSS and PBS (probe conc.: 10 µM). Note logarithmic vertical scales in panels **a-e and h**, vertical dashed line marks 5% level of H-Me-$FS_1$ fluorescence / extinction coefficient.

**Regarding fluorescence properties: $H_2$-$FS_0$** shows one absorption peak with a maximum at 503 nm in PBS (pH = 7.4) whereas **$H_2$-$FS_1$** is red-shifted slightly to 507 nm as expected for electron withdrawing substituents such as the sulfonate.[1]

**Regarding enzymatic turn-on:** To be detectable, the probes must be rapidly enzymatically processed upon cellular entry to generate a fluorescent signal. We assessed probe activation by the model enzyme porcine liver esterase (PLE, 250 ng/mL) in PBS as the least hydrolyzing buffer (**Fig S2f**). All probes are activated above spontaneous hydrolysis in PBS proving the activation by esterases as desired, but clear differences between the two probe classes are observed: all mono-capped probes are activated much faster than the doubly capped probes (in terms of percentage maximum fluorescence) which stems from their different activation profiles discussed above representing a key advantage of mono-capped probes.

**Regarding spontaneous probe hydrolysis:** We expected our sulfonated probes to be less hydrolytically stable than their non-sulfonated analogues, due to a lower tendency to aggregate and potentially to local pH depression, which both should enhance hydrolytic reactivity. We examined the probe stabilities in PBS, the standard cell culture medium DMEM (without FCS) and HBSS. PBS is a very simple cell buffer containing sodium and potassium chloride as well as sodium hydrogen- and dihydrogen-phosphates. The HBSS we used contains the same ingredients (in different amounts) plus additional salts (calcium and magnesium chloride and sulfate, sodium hydrogencarbonate), as well as glucose, to ensure longer cell viability in experiments as compared to PBS, although we expected that the stronger Lewis acids could better promote ester hydrolysis. The cell culture medium DMEM additionally contains amino acids, which we expected to give even higher ester cleavage by trans-acylation, as well as vitamins (we did not supplement it with FCS). We thus expected PBS to show the lowest spontaneous probe hydrolysis and DMEM to show the highest, while cellular viability during longer-term experiments should increase in the same order.

Indeed, the hydrolysis time-courses show that sulfonated probes are hydrolysed faster than non-sulfonated; and that the acetate probe (**$a_2$-$FS_1$**) is much more labile than any isobutyrate probe (**Fig S2g**). After 15 min incubation at 37 °C which is a typical timeframe for cell experiments, all sulfonated probes show high hydrolysis in DMEM (70% for **$a_2$-$FS_1$**, 5-25% for isobutyrate probes) which is strongly reduced in HBSS (<1% activation) and in PBS there is almost no activation (<0.1% activation except for **$a_2$-$FS_1$**) (**Fig S2h**).

**For all cellular experiments, we now used HBSS by default** as it offers low hydrolysis of chloro-stabilised isobutyrates with better cell viability maintenance than PBS.

### 3 Supporting Notes

#### 3.1 Supporting Note 1 - Cell-Impermeable Stains and Fluorogenic Probes

***Always-On Imaging Agents vs Fluorogenic Probes:*** Permanently fluorescent, cell-excluded compounds can be useful *in vitro* stains for cells with leaky membranes, since low-background images can be taken by exchanging the extracellular medium before imaging and/or by background-subtracting the homogenous signal of the extracellular medium. However, for *in vivo* imaging, "washing out" free probe is usually impossible so a high background signal remains, and this background signal usually cannot be compensated away due to its inhomogeneous 3D distribution. In this situation, fluorogenic probes offer a key advantage. Due to their fluorescence switch-on, true off→on fluorogenic probes can give high signal-to-background ratios and therefore be useful for wash-free *in vivo* imaging, as long as they are suitably stable in the extracellular medium, and as long as they are retained in those cells which originally activated them.

***Other Factors, Propidium iodide (PI):*** Conceptually, PI achieves fluorescence turn-on by DNA-intercalation, not by enzymatic bond cleavage: so, its "impermeable-off→entered-activated" switch concept is not transferable to other types of bioactives or imaging agents or even to other situations such as cell surface imaging. Even as a stain for permeabilised cells, it is optically limited in even its cellular applications, due to poor fluorescence brightness, and broad spectral peaks which makes it incompatible with many other dyes. Biologically, the DNA intercalation of PI becomes problematic when not used for live/dead staining but for live cell imaging of cells which are damaged but not necessarily dead yet, particularly in an *in vivo* context where washout of PI that has not been spontaneously uptaken cannot be ensured before endocytosis causes nonspecific uptake, and where PI's bioactivity as a DNA intercalator is then unacceptable over the longer experimental timescales of useful studies, due to its slow/very slow excretion.

***Phosphates as a non-general impermeabilising capping group:*** The dianion fluorescein diphosphate (**FDP**) is an example of an impermeabilised probe, that natively addresses cell-surface enzymes. Its two charged phosphates that also serve as fluorescence blocking groups are activated by enzymatic triggers (phosphatases); however, suitable phosphatases act at the cell surface of all healthy cells, whereupon the more lipophilic fluorescein product can then enter the cell or disengage to diffuse away into the medium; and this activation is therefore not selective for damaged cells. Due to the 2D activation surface, the activation of **FDP** is also comparably slow; and even after cellular uptake, the released fluorophore can diffuse back out of the cell which reduces the observed intracellular over extracellular fluorescence intensities.

***Evaluation of a membrane damage probe by confocal microscopy:*** Our membrane damage probe evaluation goal for *healthy cell* treatments was to assess (a) intracellular signal, that quantifies how much probe undesirably entered healthy cells and was activated by deacylation, i.e. a "false positive" signal for a membrane damage probe; to (b) extracellular signal, that quantifies spontaneous probe hydrolysis in the medium, which is also undesirable since this sets the background level against which a probe must provide higher signal for damaged cells.

#### 3.2 Supporting Note 2 - The Biology of Membrane-Damaged Cells

Cell membrane integrity can be damaged by a variety of physiological and pathological processes: from physical stress (e.g. mechanical injury) to chemical modification (e.g. peroxidation), or protein pore formation (e.g. programmed cell death, or insertion of bacterial toxins). All of these impair the separation of intracellular and extracellular spaces and allow otherwise membrane-impermeable species to cross membranes.[2,3]

Membrane-damaged but still-living cells are an interesting study population as they find themselves at the crossroads between cell death and survival: they can either heal and recover, or else they will die. This crossroads is especially relevant in the context of neurodegeneration since neurons are post-mitotic (non-dividing) cells and can therefore not be replaced by new cells once lost. For example, two recent studies have shown that the degeneration of axons (long-distance neuronal projections) can be initiated by loss of plasma membrane integrity.[4,5] In the mouse models of both studies (traumatic spinal cord contusion injury, and neuroinflammatory degeneration in the multiple sclerosis mouse model of *experimental autoimmune encephalomyelitis* (EAE)), membrane-damaged axons entered a meta-stable state from which they could completely recover (by re-establishing membrane-integrity and calcium homeostasis) or proceed to axonal fragmentation (irreversible disintegration). Diagnostic and therapeutic interventions in such situations could therefore rely on probes to selectively label these meta-stable membrane-damaged axons, or on platform chemistries to similarly design selectively membrane-damaged-cell-targeted pro-regenerative drugs.

More broadly than the cases of SCCI and EAE, studying membrane-compromised cells on the verge of cell death, or therapeutically promoting their survival, would be helpful in a variety of physiological and pathological situations: and for this, strategies to target them selectively with small molecules are needed.

The study of membrane-compromised cells on the verge of cell death, or therapeutically promoting their survival, would be helpful in a variety of physiological and pathological situations, beyond axon degeneration. Therefore, strategies to target them selectively with small molecules are needed.Initially, to investigate probe uptake and activation in damaged cells, we used the pore-forming bacterial toxin listeriolysin O (LLO) to induce membrane damage (**Fig 4c**). LLO is a protein that is secreted by bacteria to penetrate the host cell by ring-oligomerisation on the target membrane where a subsequent conformational change leads to pore formation.[6]

Next however, we followed our main longterm motivation which had been to develop turn-on membrane-damage probes for selective targeting of (membrane-)damaged axons as are commonly found in inflammatory lesions in mouse models of multiple sclerosis as well as in traumatic spinal cord contusion injury.[4,5] We therefore examined a neuronal cell line, PC12 cells, which are derived from rat pheochromocytoma and can be differentiated into neurite-extending neurons using neuronal growth factor.[7] Although the origin of altered axonal membrane permeability in neuroinflammatory lesions is unknown, we speculated that it may either be mediated by an unspecific protein pore as employed by cytotoxic immune cells or cell-autonomous cell death pathways or, since inflammatory lesions feature high levels of reactive oxygen and nitrogen species[8], may alternatively be the consequence of plasma membrane lipid peroxidation. Since LLO had already modelled membrane damage by insertion of protein pores, we now decided to test the robustness of our approach by modelling lipid peroxidation. To this end, PC12 cells were treated with 2,2'-azobis(2-amidinopropane) dihydrochloride (**AAPH**), a free-radical generating azo compound which can be used to peroxidise membrane lipids experimentally.[9] Axonal membrane damage in SCCI is caused mechanically and was not addressed in this study

Fluorescence microscopy reveals different degrees of cell-exclusion from healthy PC12 cells also deviating from the ones found in HEK and HeLa cells (see **Fig S7** for further discussion). **iPS-FS$_2$** is efficiently excluded from healthy PC12 cells, but strongly stains AAPH-damaged cells with eight-fold higher cell-fluorescence as shown by confocal imaging (**Fig 5a,b**). We speculate that the "signal enhancement" in the AAPH assay may be lower than in the LLO assay because peroxidative-type damage does not create such enormous structured voids for the probe to permeate through.

#### 3.3 Supporting Note 3 - Full Figure Legends

**Figure 2: Cell penetration and fluorogenicity of double-capped probes or fluorophores, with healthy cells. (a)** Sulfonated probes **$i_2$-$FS_1$** and **$a_2$-$FS_1$** from fluorophore **$H_2$-$FS_1$**; plus, known references **$H_2$-$FS_0$** and **$i_2$-$FS_0$**. **(b-c)** Confocal microscopy of healthy HEK cells incubated with test compounds (5 µM) for 10 min without washing (CellTracker is a cytosolic stain; scale bars: 50 µm). **(c)** Image quantification for intracellular vs extracellular fluorescence (note $log_{10}$ vertical axis; values normalised to autofluorescence of DMSO control as 1; all data: $n$ = 3). *Note: the intracellular fluorescence quantified for the active fluorophores $H_2$-$FS_1$ and $H_2$-$FS_0$ is somewhat higher than expected in this procedure, because automatic recognition of intracellular vs extracellular space using the CellTracker channel is imperfect*. **(d)** Orthogonal quantification of cellular entry-and-activation from flow cytometry on healthy HeLa cells (10 µM test compounds, 20 min incubation).

**Figure 4: Fluorogenic probes that are excluded from healthy cells but enter and are then activated inside damaged cells. (a)** Confocal microscopy images of HEK cells treated with **iMe-$FS_1$**, **iEM-$FS_1$**, **iPS-$FS_1$** and **iPS-$FS_2$** (5 µM, 10 min) ($n$ = 3); **(b)** quantified intracellular and extracellular fluorescence for **iMe-$FS_1$**, **iEM-$FS_1$**, **iPS-$FS_1$** and **iPS-$FS_2$** (5 µM, 10 min) in HEK cells, dashed line marks auto-fluorescence (DMSO control) ($n$ = 3); **(c)** Membrane damage assay overview. **(d)** Confocal microscopy images of HEK cells either untreated (without LLO) or pre-treated with LLO (0.2 µg/mL) for 5 min followed by **iPS-$FS_2$** treatment (5 µM). Images taken 10 min after probe application without prior washing ($n$ = 3). **(e)** Quantified intracellular fluorescence for each probe (5 µM, 10 min, non-wash) for healthy and LLO damaged HEK cells ($n$ = 3); **(f)** Chemical structure of **iPS-$FS_2$**. *Scale bar: 50 µm*. *Note*: "Brightness 10×" indicates images are adjusted as compared to brightness in Fig. 2.

**Figure 5: Biological and chemical scope of the fluorogenic cell-excluded probe concept. (a-b)** Works for nonspecific oxidative AAPH-induced membrane damage (PUFA peroxidation). Confocal microscopy images of PC12 cells either untreated (without AAPH) or treated with AAPH (300 mM) for 90 min followed by probe treatment (5 µM). Images taken 10 min after probe application without prior washing ($n$ = 3). **(b)** Quantified intracellular fluorescence for each probe (5 µM, 10 min, without washing) with and without pre-treatment with AAPH (300 mM, 90 min) ($n$ = 3); **(c-e) Other fluorogenic triggers can be modularly introduced to tune probe performance. (c)** Applying a GSH-labile redox trigger to our modular impermeable fluorophore system for intracellular fluorescence turn-on; **(d)** Spontaneous probe hydrolysis of ester probe **iPS-$FS_2$** and redox probe **MSS00-PS-$FS_2$** in standard cell culture medium DMEM; **(e)** Confocal microscopy images of HEK cells either untreated (without LLO) or treated with LLO (0.2 µg/mL) for 5 min followed by probe treatment (5 µM). Images taken 10 min after probe application without prior washing ($n$ = 3). *Scale bar: 50 µm*.

**Figure 6: (a-k) Sensing membrane damage in ferroptosis (T-cells in a culture of total lung lymphocytes; optional pre-treatment with RSL3 (0.5 µM) or hydrogen peroxide (0.4 mM)). (a)** Chemical structure of **iPS-$FS_2$**. **(b)** Histograms of the mean fluorescence intensity (MFI) of **iPS-$FS_2$** (administered at 50 µM for 30 min), in optionally pre-treated T cells, as assessed by flow cytometry. **(c)** Quantified MFI of **iPS-$FS_2$** of the histogram plots shown in panel **a**. **(d)** Contour plots of iPS-FS2 fluorescence vs. SSC-A pre-gated on T cells ($TCRb^+$) showing the gating on damage positive cells (**iPS-$FS_2^+$**). **(e)** Quantification of **iPS-$FS_2^+$** cells. **(f)** Chemical structure of the administered **BODIPY-C11** (RED) (reduced form). **(g-h)** Histograms of the MFI of **(g) BODIPY-C11** (OX) fluorescence and **(h) BODIPY-C11** (RED) fluorescence, following administration of **BODIPY-C11** (RED) (250 µM for 1 h). **(i-j)** Quantified MFI of **BODIPY-C11** (OX and RED) of the histogram plots shown in panels **g-h. (k)** Ratio of the MFI of **BODIPY-C11** (OX) divided by (MFI of **BODIPY-C11** (RED) plus MFI of **BODIPY-C11** (OX)), used as a proxy for the amount of lipid peroxidation, and thus as an indication of ferroptosis. **(l-n) Sensing membrane damage in necrosis in live *Drosophila* embryo.** The epithelial cell-surface-marker mcherry-Moesin (red channel) is used to outline epithelial cells; three cells of interest are annotated with the magenta, cyan and yellow arrows/outlines. Note also the presence of several macrophages which clear apoptotic epithelial cells by ingestion (so at the start of imaging they have collected multiple red-fluorescent conglomerates within themselves). **iPS-$FS_2$** was microinjected into the intervitelline space, then local necrotic tissue damage was triggered by local laser wounding using laser ablation at time zero (focal area indicated with asterisk), and probe fluorescence was imaged over the following minutes. **(l)** Experimental setup. **(m)** The ventral epithelium of the embryo was laser-wounded, resulting in expansion then rebound of the targeted tissue area (note slight movement of cells between frames), in local photobleaching of the mCherry-Moesin marker (dark zone already at the "time zero" frame collected immediately after wounding), and in necrotic cellular damage at the target site. Cells that had been entirely within the original damage zone are most rapidly and strongly labelled by **iPS-$FS_2$** activation (green channel); cells that had been partially covered by or contacting the damage zone are also labelled (e.g. yellow- and cyan-indicated epithelial cells); cells that were not contacting the damage zone are not labelled (e.g. magenta-indicated epithelial cell). Note also the two macrophages at centre-left (containing ingested mCherry conglomerates) which are not laser-damaged, and therefore are not labelled (they show up as shadows in the green channel). Images shown are Z-projections to account for curvature of the organism. Time shown in minutes; scale bars 5 µm; "A/P/V/D" indicates anterior / posterior / ventral / dorsal.

### 4 Biological characterisation

#### 4.1 Exclusion of probes and fluorophores from healthy HeLa and HEK cells

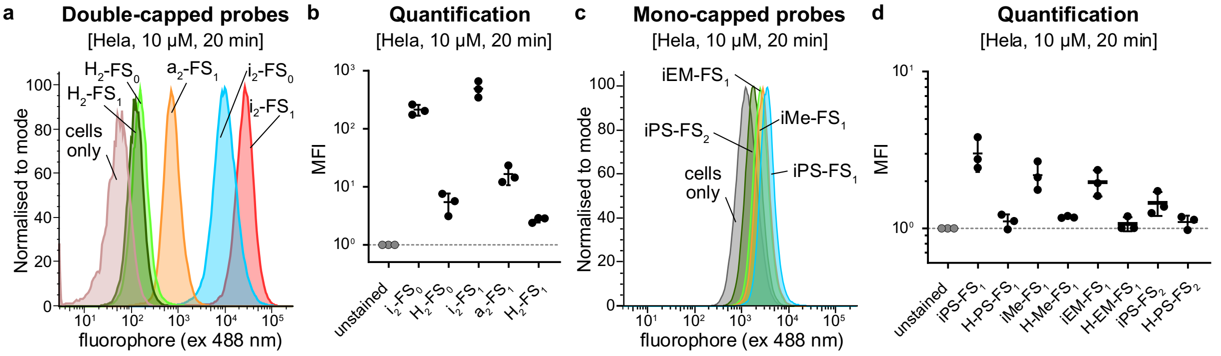

**Figure S3: Flow cytometry analysis of cell penetration of the probes and fluorophores into healthy HeLa cells. (a)** Histogram plots of cellular fluorescence (of fluorescein) for the doubly capped probes after treatment of HeLa cells for 20 min (probe conc.: 10 µM); **(b)** Mean fluorescence intensities (MFI) of doubly capped probes and fluorophores (norm. to DMSO control, *n* = 3); **(c)** Histogram plots of cellular fluorescence (of fluorescein) for the mono-capped probes after treatment of HeLa cells for 20 min (probe conc.: 10 µM); **(d)** Mean fluorescence intensities of mono-capped probes and fluorophores (norm. to DMSO control, *n* = 3).

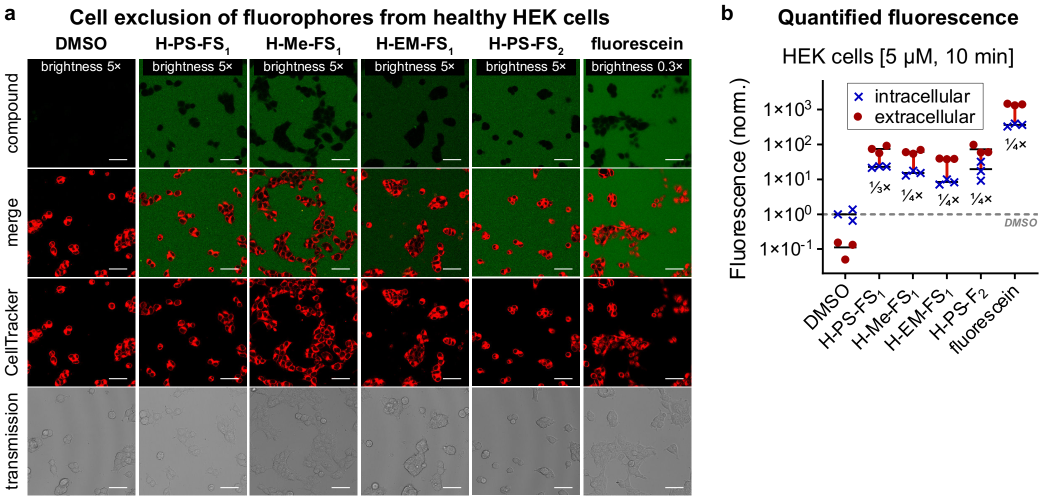

**Fig S4**: **Microscopy and quantification of cell exclusion of sulfonated fluorophores into healthy HEK cells; (a)** Confocal microscopy images of HEK cells after treatment with fluorophores (5 µM) for 10 min (non-wash, CellTracker™ CMTPX Red for cell staining) (*n* = 3); **(b)** quantified intracellular and extracellular fluorescence intensities (compound treatment: 5 µM, 10 min, non-wash) in HEK cells, horizontal line marks auto-fluorescence (DMSO control) (*n* = 3). *Scale bar: 50 µm.*

Note that the small, but real, differences in cell-uptake of the fluorophores (compare **Fig S3d**) cannot be seen by imaging because of the extremely bright extracellular signal, which is the major reason we focus on turn-on probes.

**Table S1**: Normalised cellular fluorescence values quantified from microscopy images for the cell penetration into healthy HEK cells (normalised to DMSO control (intracellular), n = 3).

| Compound | Intracellular fluorescence | | | Extracellular fluorescence | | | Mean (intra) | Mean (extra) | Ratio intra/extra |
|---|---|---|---|---|---|---|---|---|---|
| $i_2$-$FS_0$ | 417 | 526 | 301 | 4.75 | 2.93 | 3.81 | 415 | 3.83 | 108 |
| $i_2$-$FS_1$ | 325 | 213 | 171 | 4.16 | 20.49 | 8.57 | 236 | 11.1 | 21.3 |
| $a_2$-$FS_1$ | 167 | 124 | 190 | 9.43 | 16.8 | 13.2 | 160 | 13.1 | 12.2 |
| iPS-$FS_1$ | 84.6 | 85.7 | 70.9 | 3.57 | 3.07 | 3.94 | 80.4 | 3.53 | 22.8 |
| iMe-$FS_1$ | 4.42 | 5.51 | 5.01 | 2.35 | 2.63 | 3.75 | 4.98 | 2.91 | 1.71 |
| iEM-$FS_1$ | 4.43 | 4.32 | 6.27 | 1.57 | 1.58 | 2.20 | 5.01 | 1.78 | 2.81 |
| iPS-$FS_2$ | 1.92 | 2.89 | 1.61 | 1.94 | 1.76 | 1.87 | 2.14 | 1.86 | 1.15 |
| FDP | 9.60 | 21.5 | 14.7 | 10.5 | 8.85 | 11.4 | 15.3 | 10.3 | 1.49 |
| fluorescein | 362 | 398 | 323 | 1308 | 1467 | 1401 | 361 | 1392 | 0.26 |
| $H_2$-$FS_0$ | 212 | 278 | 240 | 727 | 867 | 889 | 243 | 828 | 0.29 |
| $H_2$-$FS_1$ | 235 | 158 | 195 | 517 | 571 | 513 | 196 | 534 | 0.37 |
| H-PS-$FS_1$ | 23.0 | 23.5 | 21.5 | 90.7 | 56.7 | 74.1 | 22.7 | 73.9 | 0.31 |
| H-Me-$FS_1$ | 12.9 | 17.7 | 15.1 | 60.4 | 69.8 | 53.6 | 15.2 | 61.3 | 0.25 |
| H-EM-$FS_1$ | 7.02 | 8.26 | 10.0 | 39.7 | 38.7 | 37.9 | 8.4 | 38.8 | 0.22 |
| H-PS-$FS_2$ | 9.10 | 17.2 | 32.0 | 60.0 | 59.0 | 97.9 | 19.4 | 72.3 | 0.27 |
| DMSO | 1.36 | 0.99 | 0.65 | 0.13 | 0.16 | 0.05 | **1** | 0.11 | 8.86 |

#### 4.2 LLO Damage Assay

##### 4.2.1 Entry of probes into LLO-damaged HEK cells

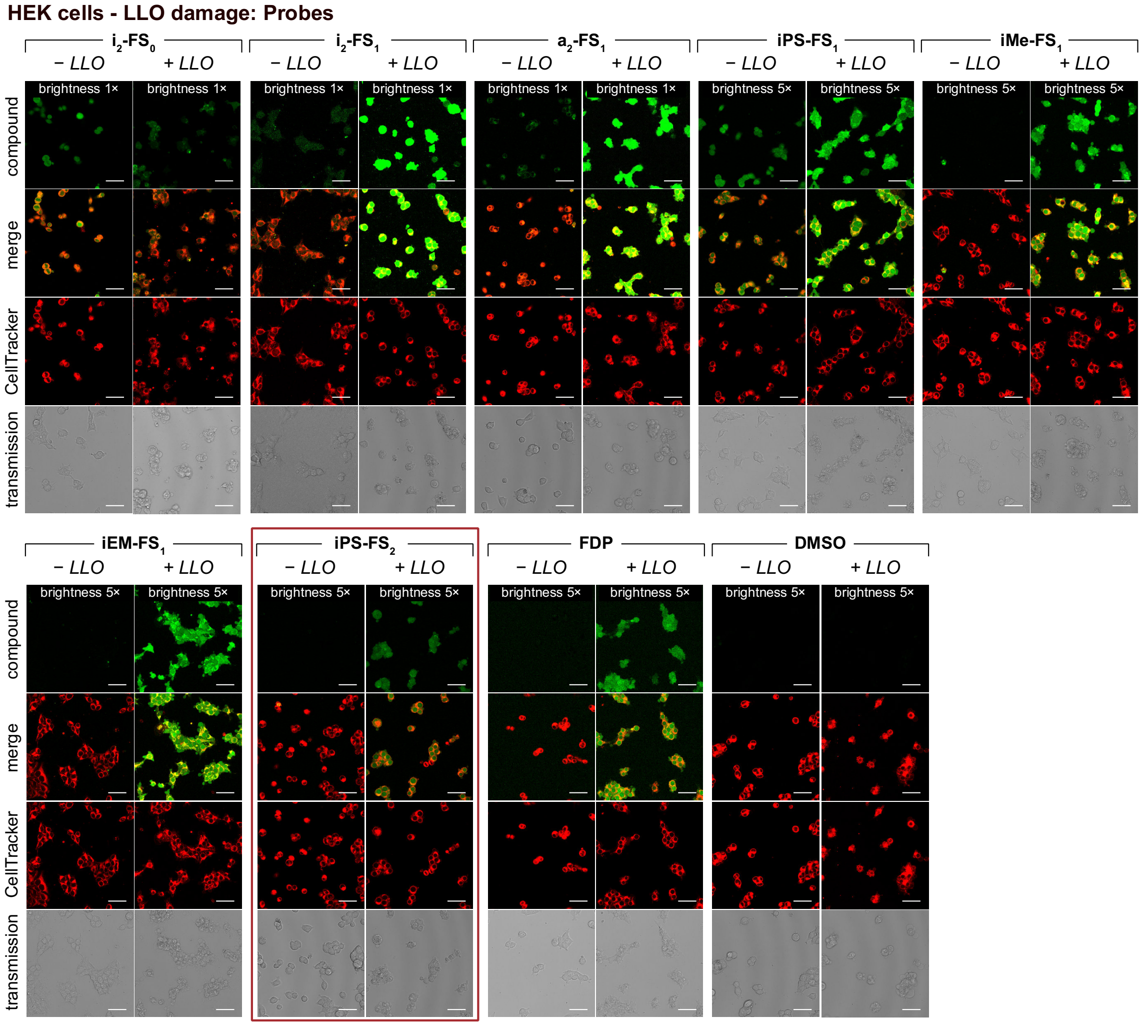

**Figure S5: Probe uptake in LLO-damaged HEK cells.** Confocal microscopy images for turn-on probes of HEK cells pre-treated with LLO (0.2 mg/mL) for 5 min followed by probe treatment (5 µM). Images taken 10 min after probe application without prior washing ($n$ = 3). *Scale bar: 50 µm.*

##### Detailed discussion of probe uptake

Quantifying the intracellular fluorescence reveals significant differences between the probes (also see **Fig 4e**). The membrane permeable probe **H$_2$-FS$_0$** shows lower fluorescence with LLO than without which could be explained by faster signal loss due to excretion of the released fluorophore **H$_2$-FS$_0$** through the created pores. **i$_2$-FS$_1$**, which is also permeable to healthy cells, on the other hand shows even higher signal generation in damaged cells. We reason that this mono-sulfonated probe already features very low permeability because of the polar sulfonate but very fast intracellular signal generation (see above: faster esterase activation and lower hydrolytic stability). Thus, we suspect that the better cell penetration into LLO-damaged cells outcompetes the signal loss of the released fluorophore. Also, the membrane-permeable probes **a$_2$-FS$_1$** and **iPS-FS$_1$** show moderate signal increase with LLO. However, all these membrane permeable probes are not suitable for membrane-damage imaging from low background as they already give high fluorescence with healthy cells (besides their moderate uptake ratios). For this goal the cell-excluded probes **iMe-FS$_1$**, **iEM-FS$_1$** and **iPS-FS$_2$** are more suited. Quantification shows around 30-fold signal increase in damaged cells for all three probes making them valuable tools to distinguish healthy from membrane-damaged cells with high contrast and sensitivity by far outperforming commercially available **FDP** (only 7-fold increase). **iPS-FS$_2$** reveals the best performance as it features the lowest fluorescence in healthy cells enabling the highest sensitivity. The permanently fluorescent fluorophores show high extracellular fluorescence and can only penetrate into permeabilised cells (**Fig S6**).

##### 4.2.2 Entry of fluorophores into LLO-damaged HEK cells

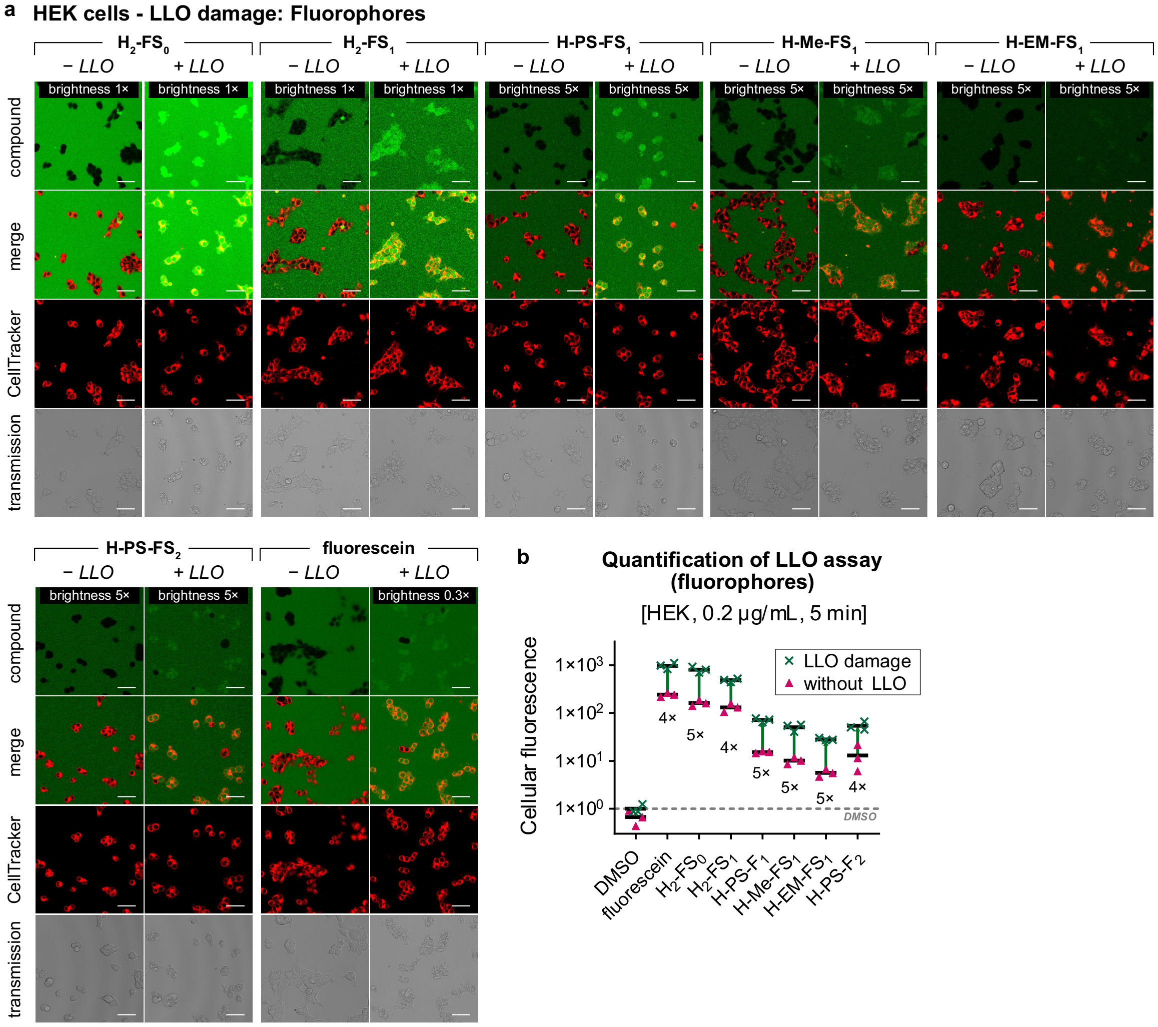

**Figure S6: Fluorophore uptake in LLO-damaged HEK cells. (a)** Confocal microscopy images for fluorophores of HEK cells pre-treated with LLO (0.2 mg/mL) for 5 min followed by probe treatment (5 µM). Images taken 10 min after probe application without prior washing (*n* = 3). **(b)** Quantified intracellular fluorescence for each fluorophore (5 µM, 10 min, non-wash) for healthy and LLO damaged HEK cells (*n* = 3). *Scale bar: 50 µm.*

##### 4.2.3 Data (normalised): Exclusion or entry into LLO-damaged HEK cells

**Table S2**: Normalised intracellular fluorescence values quantified from microscopy images for the LLO membrane damage assay in HEK cells (normalised to DMSO control with LLO (such that its intracellular fluorescence value is 1), n = 3).

| Compound | +LLO | | | −LLO | | | Mean (+LLO) | Mean (−LLO) | Ratio +/−LLO |
|---|---|---|---|---|---|---|---|---|---|
| $i_2$-$FS_0$ | 85.3 | 117 | 142 | 280 | 352 | 202 | 115 | 278 | 0.41 |
| $i_2$-$FS_1$ | 3391 | 1416 | 2251 | 218 | 143 | 115 | 2353 | 158 | 14.9 |
| $a_2$-$FS_1$ | 957 | 1122 | 1038 | 112 | 83 | 127 | 1039 | 107 | 9.67 |
| iPS-$FS_1$ | 204 | 174 | 178 | 56.7 | 57.5 | 47.5 | 185 | 53.9 | 3.44 |
| iMe-$FS_1$ | 89.4 | 102 | 111 | 2.97 | 3.69 | 3.36 | 101 | 3.34 | 30.2 |
| iEM-$FS_1$ | 109 | 120 | 132 | 2.97 | 2.90 | 4.20 | 121 | 3.36 | 36.0 |
| iPS-$FS_2$ | 48.3 | 43.6 | 43.0 | 1.29 | 1.94 | 1.08 | 45.0 | 1.44 | 31.3 |
| FDP | 77.8 | 58.5 | 66.1 | 6.43 | 14.4 | 9.85 | 67.5 | 10.2 | 6.59 |
| fluorescein | 986 | 1106 | 834 | 242 | 267 | 217 | 976 | 242 | 4.03 |
| $H_2$-$FS_0$ | 803 | 920 | 704 | 142 | 187 | 161 | 809 | 163 | 4.96 |
| $H_2$-$FS_1$ | 493 | 526 | 442 | 157 | 106 | 131 | 487 | 131 | 3.71 |
| H-PS-$FS_1$ | 77.3 | 64.3 | 73.2 | 15.4 | 15.7 | 14.4 | 71.6 | 15.2 | 4.72 |
| H-Me-$FS_1$ | 54.3 | 56.6 | 40.6 | 8.62 | 11.9 | 10.1 | 50.5 | 10.2 | 4.96 |
| H-EM-$FS_1$ | 30.1 | 27.5 | 27.0 | 4.70 | 5.54 | 6.71 | 28.2 | 5.65 | 4.99 |
| H-PS-$FS_2$ | 45.3 | 50.3 | 66.3 | 6.10 | 11.5 | 21.4 | 53.9 | 13.0 | 4.15 |
| DMSO | 1.25 | 0.88 | 0.88 | 0.91 | 0.66 | 0.43 | **1** | 0.67 | 1.49 |

#### 4.3 AAPH Damage Assay

##### 4.3.1 Entry of probes into AAPH-damaged PC12 cells

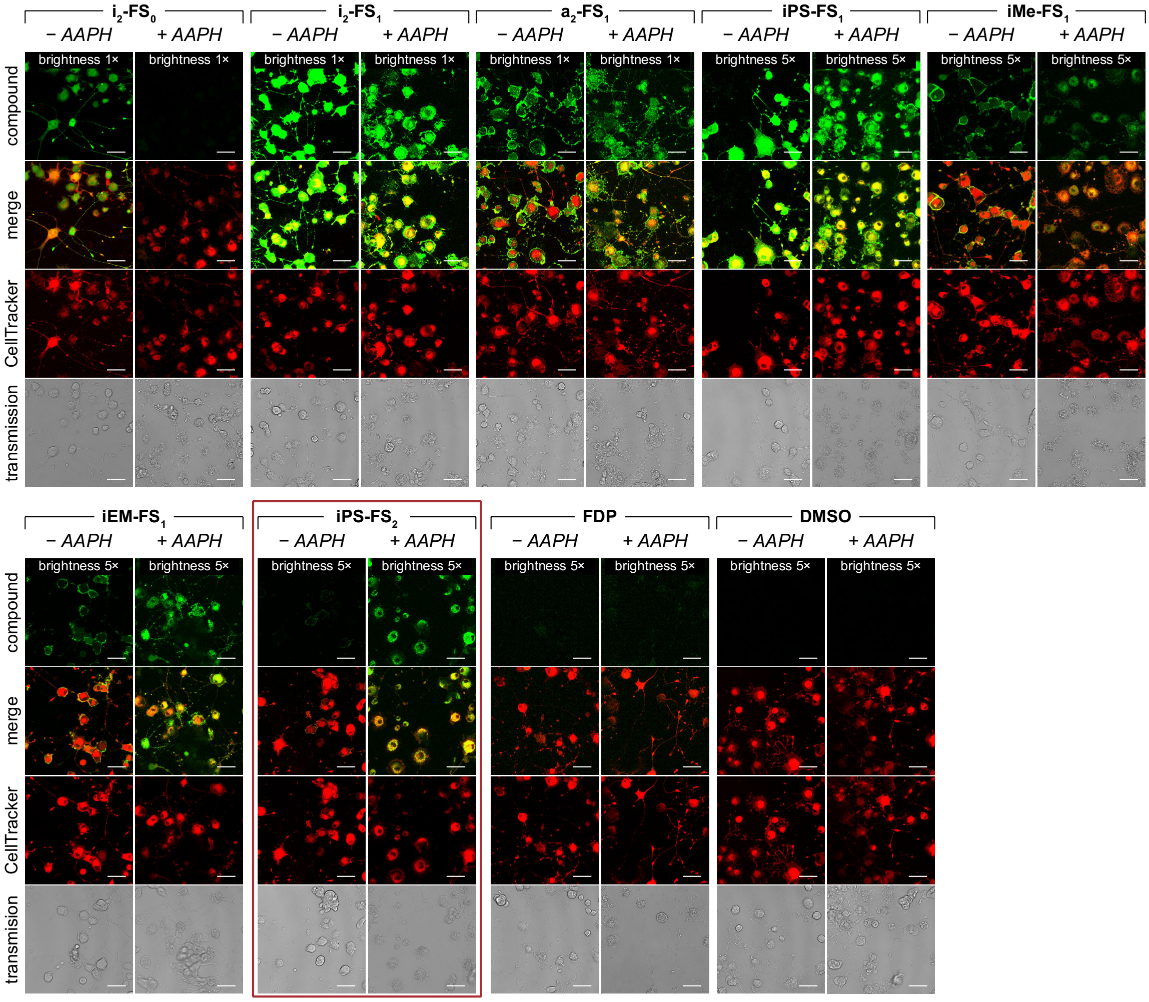

**Figure S7: Probe uptake in AAPH-damaged PC12 cells.** Confocal microscopy images for turn-on probes of PC12 cells pre-treated with AAPH (300 mM) for 90 min followed by probe treatment (5 µM). Images taken 10 min after probe application without prior washing ($n$ = 3). *Scale bar: 50 µm*.

Note: we interpret the **i$_2$-FS$_0$** result of lower fluorescence in treated than in untreated cells as indicating that its leak-out after activation is more severe from damaged than healthy cells, matching the trend from the LLO assay.

Fluorescence microscopy reveals different degrees of cell-exclusion from healthy PC12 cells which deviate from the ones found in HEK and HeLa cells. Lipophilic mono-sulfonates (**i$_2$-FS$_1$**, **a$_2$-FS$_1$**, **iPS-FS$_1$**) are well permeable as observed before, but interestingly **i$_2$-FS$_1$** is even brighter than its non-sulfonated parent **i$_2$-FS$_0$** in healthy PC12 cells (**Fig S7, Fig 5b**). The more polar mono-capped probes (**iMe-FS$_1$**, **iEM-FS$_1$**) turned out to be significantly more permeable than in HEK and HeLa cells, suggesting cell-type specific differences in membrane permeability. The more polar disulfonated **iPS-FS$_2$** however was efficiently excluded by healthy PC12 cells. The higher fluorescence in healthy cells brings along low turn-on ratios (low selectivity) for all probes except for **iPS-FS$_2$** (which also performed best in the LLO assay). The permanently fluorescent fluorophores are again excluded from healthy cells but penetrate into leaky cells as expected for the more polar open-form fluoresceins (**Fig S8**).

##### 4.3.2 Entry of fluorophores into AAPH-damaged PC12 cells

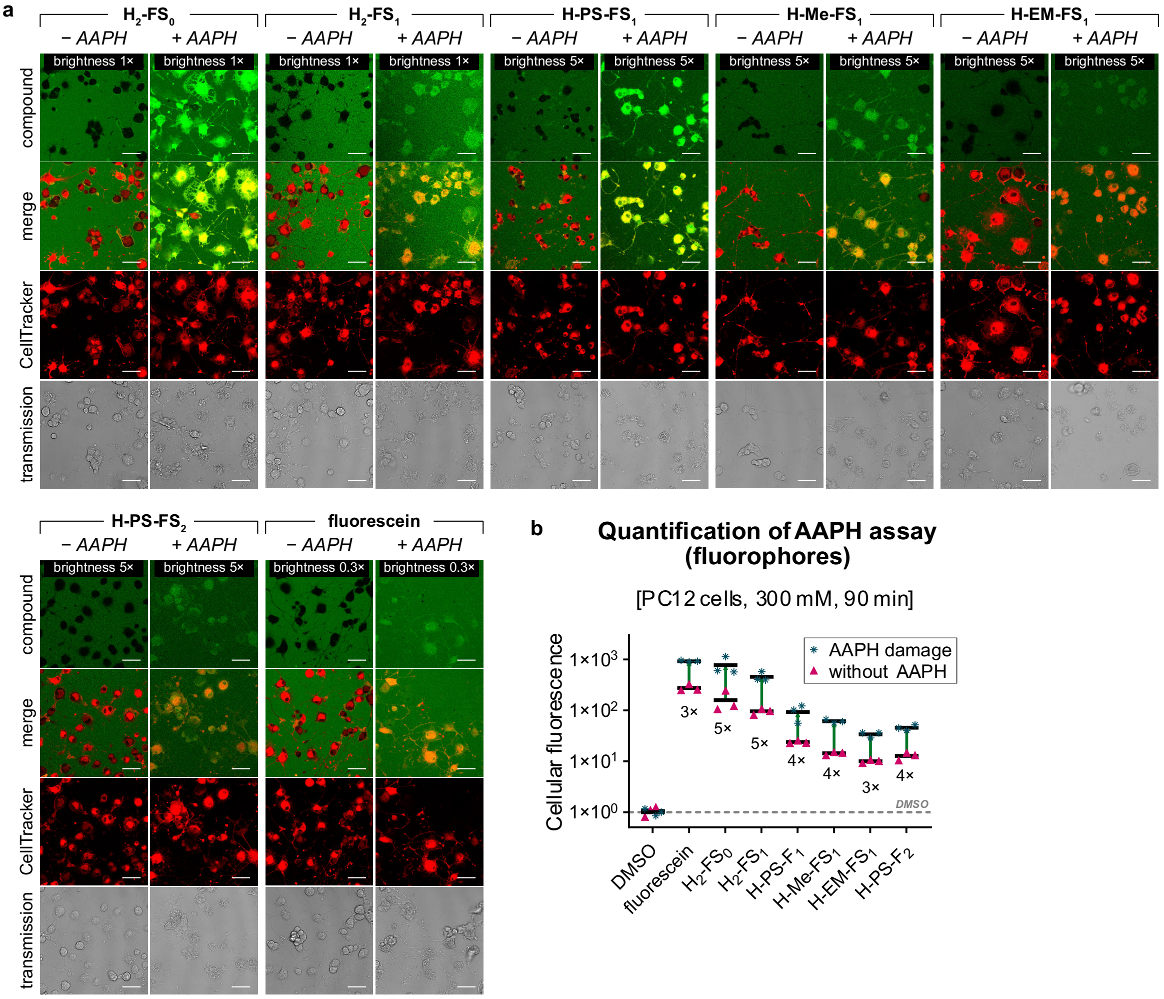

**Figure S8: Fluorophore uptake in AAPH-damaged PC12 cells. (a)** Confocal microscopy images for fluorophores of PC12 cells pre-treated with AAPH (300 mM) for 90 min followed by probe treatment (5 µM). Images taken 10 min after probe application without prior washing ($n$ = 3). **(b)** Quantified intracellular fluorescence for each fluorophore (5 µM, 10 min, non-wash) for healthy and AAPH damaged PC12 cells ($n$ = 3). *Scale bar: 50 µm*.

##### 4.3.3 Data (normalised): Exclusion or entry into AAPH-damaged PC12 cells

**Table S3**: Normalised intracellular fluorescence values quantified from microscopy images for the AAPH membrane damage assay (normalised to DMSO control (+AAPH), n = 3).

| Compound | +AAPH | | | −AAPH | | | Mean (+AAPH) | Mean (−AAPH) | Ratio +/−AAPH |
|---|---|---|---|---|---|---|---|---|---|
| $i_2$-$FS_0$ | 52.4 | 23.7 | 14.1 | 643 | 476 | 310 | 30.1 | 476 | 0.06 |
| $i_2$-$FS_1$ | 717 | 823 | 332 | 2633 | 2668 | 1865 | 624 | 2389 | 0.26 |
| $a_2$-$FS_1$ | 390 | 871 | 390 | 243 | 512 | 335 | 550 | 363 | 1.51 |
| iPS-$FS_1$ | 82.2 | 95.5 | 55.6 | 141 | 259 | 155 | 77.8 | 185.2 | 0.42 |
| iMe-$FS_1$ | 57.7 | 31.2 | 48.8 | 34.1 | 34.8 | 28.8 | 45.9 | 32.5 | 1.41 |
| iEM-$FS_1$ | 62.7 | 116.0 | 63.0 | 27.1 | 34.5 | 20.6 | 80.6 | 27.4 | 2.94 |
| iPS-$FS_2$ | 40.7 | 45.8 | 26.6 | 3.97 | 5.35 | 5.18 | 37.7 | 4.83 | 7.80 |
| FDP | 20.4 | 13.3 | 24.0 | 5.57 | 4.12 | 7.52 | 19.2 | 5.74 | 3.35 |
| fluorescein | 920 | 901 | 955 | 256 | 250 | 328 | 925 | 278 | 3.33 |
| $H_2$-$FS_0$ | 1151 | 604 | 569 | 246 | 106 | 123 | 775 | 158 | 4.90 |
| $H_2$-$FS_1$ | 411 | 576 | 395 | 82.1 | 97.2 | 107 | 460 | 95.4 | 4.83 |
| H-PS-$FS_1$ | 122 | 101 | 56.0 | 22.8 | 25.7 | 23.1 | 93.1 | 23.9 | 3.90 |
| H-Me-$FS_1$ | 66.8 | 57.2 | 59.8 | 15.3 | 13.2 | 14.7 | 61.3 | 14.4 | 4.26 |
| H-EM-$FS_1$ | 36.2 | 29.0 | 36.2 | 10.1 | 10.8 | 9.2 | 33.8 | 10.0 | 3.36 |
| H-PS-$FS_2$ | 45.0 | 52.4 | 40.0 | 10.5 | 14.8 | 13.2 | 45.8 | 12.8 | 3.57 |
| DMSO | 0.99 | 1.17 | 0.84 | 0.82 | 1.26 | 1.10 | **1** | 1.06 | 0.95 |

#### 4.4 Dextran conjugates: rationale and performance

An alternative approach for fluorogenic probes to selectively report on cell membrane damage might be to develop macro-molecular probes without specific uptake mechanisms, that are excluded from healthy cells on the basis of their size (rather than their polarity), yet passively enter more porous damaged cells. Although the focus of this work was on small molecule probes, we briefly tested the accessibility of this approach.

Intensely fluorescent, cell-excluded macromolecular reporters (typically bearing several fluorophores per macromolecule) are useful for e.g. anatomical tracking of blood vessels or cell surface labelling. Dextrans are biologically "innocent" platform macromolecules that can be obtained commercially in good size purity from low kDa up to mid MDa and bearing a large range of fluorophores or reactive functional groups with various degrees of labelling.

Initial testing of permanently fluorescent dextrans indicated increased PC12 cellular uptake following membrane damage by AAPH treatments (radical PUFA peroxidation), for commercial, permanently fluorescent dextran sizes 6-120 kDa, with peak effect around 40-70 kDa (assessed after washing to remove the cell-excluded dextran fraction).

Since washing brings complications in 2D cell culture and cannot be performed *in vivo*, we then wished to test fluorogenic dextrans in a no-wash protocol; however, these are barely reported (first monofunctional fluorogenic dextran only published in 2018[10]) and there are no commercial suppliers. We therefore prepared the isobutyrate-capped, chloro-stabilised fluorogenic fluorescein NHS ester **NHS-i-Flu** because we expected commercially available fluorescein diacetate to be labile to spontaneous hydrolysis at pH~7.4 based on the observations of Raines and others.[11] We chose a mono- capped probe since its simple off/on unmasking may make it more suitable for quantification than the known doubly-capped probe.[10] Fina Biosolutions LLC (MD, USA) performed custom conjugation of **NHS-i-Flu** to 70 kDa dextran (**i-Flu-AmDex**), however, followup testing of this custom-made fluorogenic **NHS-i-Flu**-derived dextran did not indicate any increase of cellular entry (and intracellular activation and retention) following AAPH damage (**Fig S9**); and results in the same assay without washing contraindicated any significant increase of uptaken signal from the permanently fluorescent carboxyfluorescein-derived dextrans (**Fig S9**: 70 kDa; also tested: 10, 40, 2000 kDa).

Therefore, these investigations into damage-selectivity of macromolecular uptake were not continued.

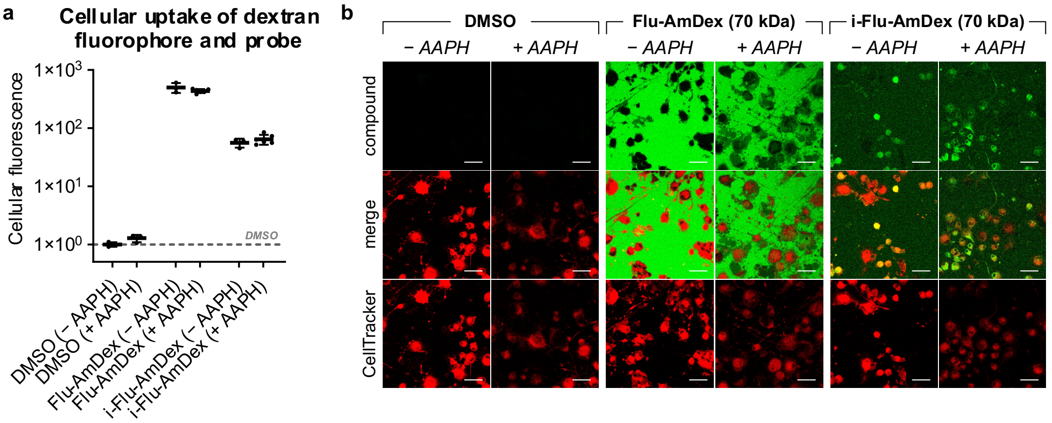

**Fig S9: Initial results for dextran conjugates. (a)** Quantified intracellular signal from permanently fluorescent, fluorescein dextran conjugate after washing (shown: 70 kDa) and from **NHS-i-Flu-**derived fluorogenic dextran conjugates (20 µM, 20 min, without washing) with and without pre-treatment with AAPH (300 mM, 90 min). **(b)** Confocal microscopy images of PC12 cells either untreated (without AAPH) or treated with AAPH (300 mM) for 90 min followed by treatment with dextran-conjugates (20 µM). Images taken 20 min after probe application without prior washing. *Scale bar: 50 µm*

#### 4.5 Concept adaptation into the disulfide-reduction probe MSS00-PS-FS$_2$

##### 4.5.1 Probe stability and GSH activation

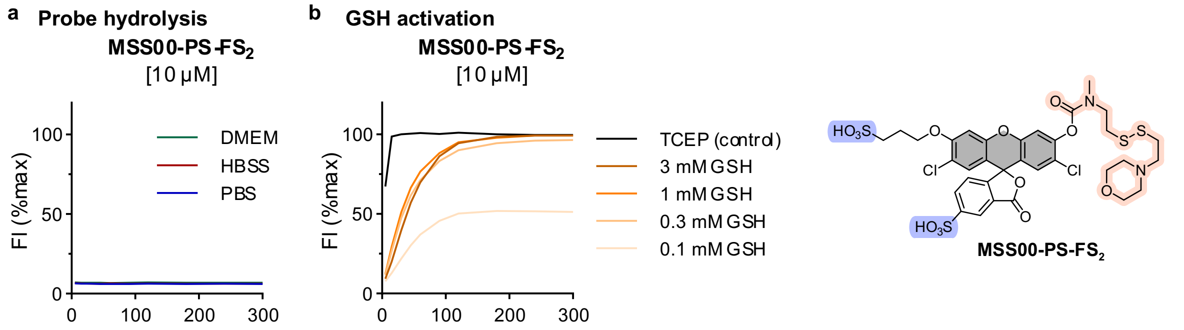

**Figure S10: Hydrolytic stability and GSH activation of MSS00-PS-FS$_2$.** (a) Probe hydrolysis of **MSS00-PS-FS$_2$** in PBS, HBSS and DMEM over 5 h (probe concentration 10 µM); **(b)** GSH activation time-course over 5 h for **MSS00-PS-FS$_2$** with different GSH concentrations (probe concentration: 10 µM).

##### 4.5.2 Exclusion or entry into membrane damaged cells

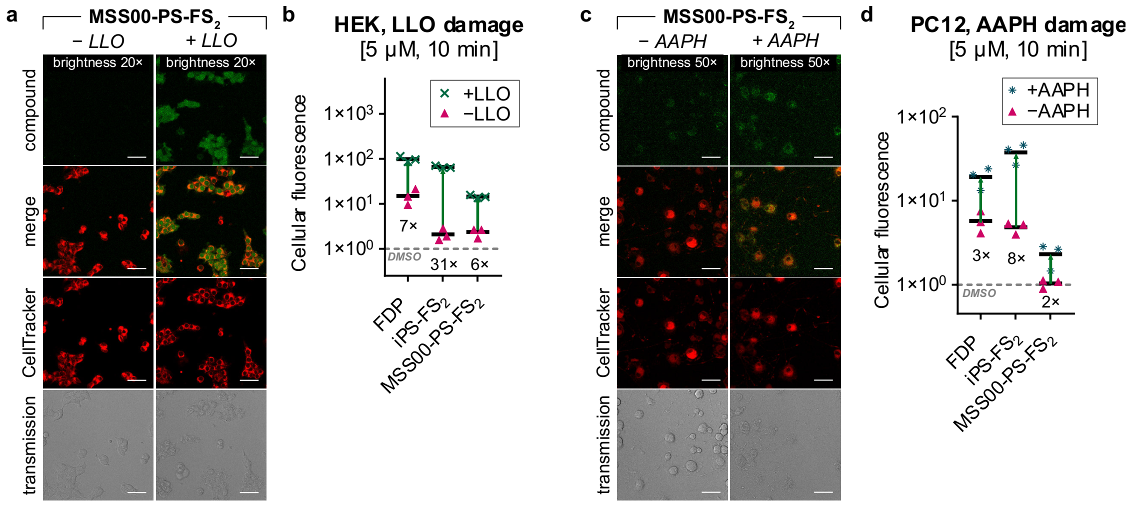

**Figure S11: Membrane damage selectivity of MSS00-PS-FS$_2$: (a)** Confocal microscopy images for MSS00-PS-FS$_2$ of HEK cells pre-treated with LLO (0.2 mg/mL) for 5 min followed by probe treatment (5 µM). Images taken 10 min after probe application without prior washing ($n$ = 3). **(b)** Quantified intracellular fluorescence for each fluorophore (5 µM, 10 min, non-wash) for healthy and LLO damaged HEK cells ($n$ = 3). **(c)** Confocal microscopy images of PC12 cells either untreated (without AAPH) or treated with AAPH (300 mM) for 90 min followed by probe treatment (5 µM). Images taken 10 min after probe application without prior washing ($n$ = 3). **(d)** Quantified intracellular fluorescence for each probe (5 µM, 10 min, without washing) with and without pre-treatment with AAPH (300 mM, 90 min) ($n$ = 3). *Scale bar: 50 µm.*

##### 4.5.3 Data (normalised): Exclusion or entry into LLO-damaged HEK cells

**Table S2**: Normalised intracellular fluorescence values quantified from microscopy images for the LLO membrane damage assay in HEK cells (normalised to DMSO control with LLO (such that its intracellular fluorescence value is 1), n = 3).

| Compound | +LLO | | | −LLO | | | Mean (+LLO) | Mean (−LLO) | Ratio +/−LLO |
|---|---|---|---|---|---|---|---|---|---|
| MSS00-PS-FS$_2$ | 14.4 | 13.5 | 15.8 | 1.73 | 2.66 | 2.66 | 14.6 | 2.35 | 6.21 |
| DMSO | 1.25 | 0.88 | 0.88 | 0.91 | 0.66 | 0.43 | 1 | 0.67 | 1.49 |

##### 4.5.4 Data (normalised): Exclusion or entry into AAPH-damaged PC12 cells

**Table S3**: Normalised intracellular fluorescence values quantified from microscopy images for the AAPH membrane damage assay (normalised to DMSO control (+AAPH), n = 3).

| Compound | +AAPH | | | −AAPH | | | Mean (+AAPH) | Mean (−AAPH) | Ratio +/−AAPH |
|---|---|---|---|---|---|---|---|---|---|
| MSS00-PS-FS$_2$ | 2.65 | 2.83 | 1.46 | 1.09 | 1.12 | 0.898 | 2.31 | 1.04 | 2.23 |
| DMSO | 0.99 | 1.17 | 0.84 | 0.82 | 1.26 | 1.10 | 1 | 1.06 | 0.95 |

#### 4.6 Necrosis imaging in the fly

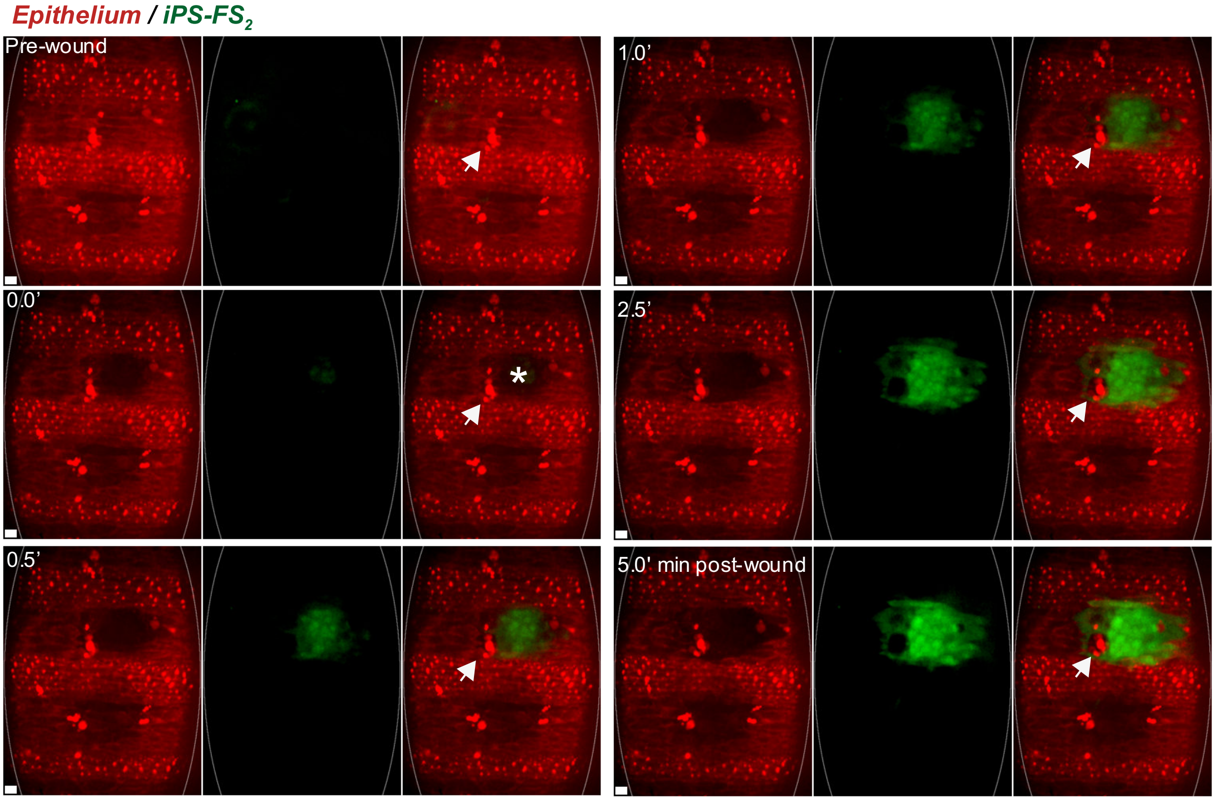

**Figure S12**: **iPS-FS$_2$ labels *in vivo* necrotic tissue damage within wounded *Drosophila* embryo.** Ventral epithelium (mcherry-Moesin, red channel, oval outline) of *Drosophila* embryo was locally wounded using laser ablation (asterisk). Laser-wounding results in necrotic tissue damage, which was rapidly labelled with previously microinjected **iPS-FS$_2$** (green). Time = minutes, scale bars = 10 μm. For further details and discussion, see **Fig 6** with its full legend which is given in **Supporting Note 3**.

### 5 Biological Materials and Methods

#### 5.1 Cell lines

**HeLa** cells were obtained from German Collection of Microorganisms and Cell Cultures (DSMZ Cat No. ACC 57) and grown in Dulbecco's modified Eagle's medium (DMEM, Sigma-Aldrich, D1145) supplemented with 10% heat-inactivated fetal bovine serum (Sigma-Aldrich, F0804), penicillin (100 U/mL) and streptomycin (100 µg/mL) (Sigma-Aldrich, P4333) at 37°C and 5% $CO_2$. Cell growth was monitored using an inverted microscope (Nikon Eclipse Ti).

**HEK 293T** cells were obtained from ATCC (Cat. No. crl-3216) and grown in DMEM (ThermoFisher 21885108) supplemented with 10% FBS (Biochrom S0615) and 1% Penicillin/Streptomycin (ThermoFisher 15140122) at 37°C and 5% $CO_2$. Cell growth was monitored using an inverted microscope (Leica DMi1).

**PC12** cells were obtained from Sigma-Aldrich (Cat. No. 88022401) and grown in RPMI (ThermoFisher 61870010) supplemented with 10% Horse Serum (Sigma-Aldrich H1138), 5% FBS and 1% Penicillin/Streptomycin at 37°C and 5% $CO_2$. Cell growth was monitored using an inverted microscope (Leica DMi1).

#### 5.2 Cell-free characterisation

**Plate reader instrument**

Raw fluorescence readout of cell-free activity was performed with a FluoStar Omega plate reader from BMG Labtech, Ortenburg (Germany), set to ex: 485bp10 and em: 520lp. All plate reader experiments were performed in black 96 well plates.

**Cell media for probe stability assay**

- Phosphate buffered saline (PBS) from Sigma Aldrich (Cat. No. D8537)
- Hanks´ balanced salt solution (HBSS) from Sigma Aldrich (Cat. No. H6648)
- Dulbecco´s modified eagle´s medium from Sigma Aldrich (Cat. No. D1145) without FBS supplementation

**Probe stability assay**

Water (10 µL) was placed in a black 96 well plate and the probes/fluorophores were added from a 1 mM stock solution in DMSO (1 µL) (water stock for H-PS-$FS_2$ and iPS-$FS_2$). Then the different media PBS, HBSS or DMEM (89 µL) were added to the probes with a multipipette (for smallest possible differences in the starting time) to reach a final concentration of 10 µM. All samples in technical duplicates. The fluorescence intensity in each well was measured with the plate reader at different time-points (0 min, 15 min, 30 min, 1 h, 2 h, 3 h, 4 h, 5 h, 6 h). Between measurements the samples were incubated at 37 °C under air atmosphere. At the end of the experiment, a piperidine solution (50 mM, 100 µL) was added to each well to determine the maximum intensity in each well. The recorded data were background subtracted (fluorescence at 0 min) and normalised to their maximum fluorescence (determined after 2 h incubation with piperidine).

**Esterase assay**

PBS (49 µL) was placed in a black 96 well plate and the probes/fluorophores were added from a 1 mM stock solution in DMSO (1 µL) (water stock for H-PS-$FS_2$ and iPS-$FS_2$). Porcine liver esterase (lyophilised powder, ≥15 units/mg, purchased from Sigma Aldrich, Cat.-No.: E3019) was diluted in PBS to give a stock solution of 500 ng/mL. The esterase stock solution was added to the probes with a multipipette (for smallest possible differences in the starting time) to reach a final probe/fluorophore concentration of 10 µM and a esterase concentration of 250 ng/mL (all samples in technical duplicates). The probes/fluorophores were placed in the pre-heated plate reader and incubated at 37 °C for the whole experiment. The fluorescence intensity in each well was measured every two minutes for the total duration of 90 min. At the end of the experiment, a piperidine solution (50 mM, 100 µL) was added to each well to determine the maximum intensity in each well. The recorded data were background subtracted (fluorescence at 0 min) and normalised to their maximum fluorescence (determined after 2 h incubation with piperidine).

#### 5.3 Flow cytometry

##### Instrument and data analysis

Flow cytometry experiments were performed on a LSR Fortessa flow cytometer (Becton Dickinson) run by BD FACSDiva software (version 8.0.1). 20,000 events per technical replicate were analysed. Channel 530/30 was used for probe/fluorophore quantification and channel 780/60 was used for Zombie NIR Dye quantification. FlowJo software from BD (version 10.8.1) was used for data analysis. Live/dead gating was performed from the heat-treated cells (for gating settings see **Fig S13**) and only live cells were used for fluorescence evaluation. All quantified fluorescence values were normalised to the auto-fluorescence of untreated cells (DMSO control).

##### Sample preparation

HeLa cells were seeded in 24-well plates (250.000 cells / well) and cultured in DMEM for 16 h. The medium was removed and the cells were carefully rinsed with HBSS (3 × 250 µL). The probes and fluorophores were prepared in HBSS from DMSO stocks (final compound concentration: 10 µM, final DMSO concentration: 1 %) and added to the cells (250 µL each). The plate was incubated at 37 °C under air atmosphere for 20 min. Then the medium was removed, the cells were trypsinised with 1 × Trypsin in PBS and centrifuged for 5 min at 500g. The supernatant was removed and the cells were resuspended in PBS (200 µL; + 1% FBS; + 1:200 Zombie NIR Fixable Dye from BioLegend®, cat.-no.: 423105) and placed on ice until measurement at the flow cytometer.

Live/dead control: these cells were not incubated with a probe/fluorophore but heated to 70 °C for 3 min in HBSS before applying the Zombie staining. Otherwise, the cells were treated like all other conditions.

#### 5.4 Confocal microscopy

##### Instrument

Confocal live cell imaging was performed at the Core Facility Bioimaging of the LMU Biomedical Center with an inverted Leica SP8X microscope, equipped with Argon laser, WLL2 laser (470 - 670 nm) and acusto-optical beam splitter. Live cells were recorded at 37 °C. For the duration of the assay, cells were kept in Hank's Balanced Salt Solution (HBSS, ThermoFisher 14025092) for minimal spontaneous hydrolysis of the probes and to allow incubation without $CO_2$.

##### Acquisition Settings

Images were acquired with a 20× 0.75NA objective and additional optical zoom of 2×. Image pixel size was 284×284 nm. The following fluorescence settings were used: probes/fluorophores (from here on comprehensively referred to as "compounds", excitation 488 nm (Argon), emission 500 – 540 nm) and CellTracker (excitation 594 nm (WLL), emission 605 – 645 nm). Recording was performed sequentially to avoid bleed-through. Compounds and CellTracker were recorded with hybrid photo detectors (HyDs), a transmitted light image was generated with a conventional photomultiplier tube. The microscope was programmed to take three images of different fields of view per condition tested, which were focused using reflection-based adaptive focus control. The scan was started after 10 min of compound incubation. Bias deriving from time delays between compounds imaged in different wells was minimised by permutation of the positioning of compounds between replicates. The time-delay between LLO- or AAPH-treated cells and their respective sham-treated controls was constant (2.5 min) due to the setup of the plate.

To account for vast brightness differences between the compounds, which were larger than the dynamic range of the detectors, brighter compounds were detected using narrower and red-shifted emission windows. $H_2$-$FS_1$, $i_2$-$FS_1$ and $a_2$-$FS_1$ were detected at 580 - 585 nm or 560 - 565 nm for HEK-LLO and PC12-AAPH assays respectively. $H_2$-$FS_0$, $i_2$-$FS_0$, fluorescein and FDP were detected at 545 - 570 nm or 550 - 565 nm for HEK-LLO and PC12-AAPH assays respectively. The normalisation factor was determined by taking a sequential scan of both detection windows (original 500 – 540 nm and red-shifted window) at low 488 nm Argon laser power.

##### Sample preparation

Assays were performed in 8-well glass bottom chambered coverslips (ibidi 80827). For HEK cell assays, chambered coverslips were coated one day before experiment by applying a 0.1 mg/mL poly-D-lysine (Sigma-Aldrich P7280) solution for 2 h at 37°C. HEK cells were seeded into the coated wells at a density of 30.000 cells per cm². For PC12 cell assays, chambered coverslips were coated six days before the experiment by applying a 0.1 mg/ml poly-D-lysine solution for 2 h at 37°C, followed by collagen IV (Sigma-Aldrich H4417) coating at 0.1 µg / $cm^2$ and 37°C overnight. PC12 cells were separated by drawing them ten times through a 30 G needle and then seeded into the coated wells at a density of 10.000 cells per cm². Two hours after seeding, medium was changed to differentiation medium (RPMI supplemented with 1% Horse Serum, 1% Penicillin/Streptomycin and 100 ng/ml ß-NGF (ThermoFisher 13257-019)). Medium was again changed to fresh differentiation medium 48 – 72 h after seeding.

##### 5.4.1 LLO and AAPH damage assay procedures

**HEK-LLO assays** were performed with the following timeline: To stain all cells irrespectively, HEK cells were incubated with 0.5 µM CellTracker$^{TM}$ Red CMTPX Dye (ThermoFisher C34552) at 37 °C for 15 min. Medium was then changed to ice cold HBSS containing 0.2 µg/ml lysteriolysin O (LLO, abcam ab83345) or ice cold HBSS only for control wells. Cells were incubated on ice for 5 min. Medium was then changed to 5 µM compound solutions prepared with 37 °C warm HBSS and chambered coverslips were subsequently put onto the microscope stage (37 °C).

**PC12-AAPH assays** were performed with the following timeline: To stain all cells irrespectively, PC12 cells were incubated with 0.5 µM CellTracker$^{TM}$ Red CMTPX Dye at 37 °C for 15 min. Medium was then changed to HBSS containing 300 mM AAPH (Cayman Chemical Cay82235-10) and 100 ng/ml ß-NGF or HBSS with ß-NGF only for control wells. Cells were incubated at 37°C on the microscope stage for 90 min. Medium was then changed to 5 µM compound solutions.

##### 5.4.2 Image analysis

Images were analysed using Fiji ImageJ. Based on the CellTracker channel, in which all cells are stained irrespective of membrane damage, images were segmented using the "Trainable Weka Segmentation" plugin. The classifier was trained individually for every single image. The plugin generates a mask that defines two regions: Intracellular and extracellular. The intracellular region of interest (ROI) was then eroded by 10 pixels to exclude the periphery of cells (where they are very thin and show low signal for both CellTracker and probes) from analysis. The resulting mask was then applied to the test probe channel, to obtain an intracellular and extracellular mean fluorescence intensity value.

For bright probes detected with red-shifted emission windows, these intensity values had to be multiplied with a normalisation factor which was obtained by measuring and averaging over mean fluorescence intensity of ten cells in original vs red-shifted detection window, thus obtaining a mean normalisation factor.

Images were excluded from analysis if they were out of focus, showed less than three cells, overgrown cells (> 66% of area covered by cells) or an intolerable amount of dead cells or debris. If more than one image had to be excluded, the complete replicate (LLO- or AAPH- and sham-treated condition) was repeated for the affected probe.

Data was plotted using GraphPad Prism. The same primary data was used for to quantify intra-/extracellular distribution of probes in healthy HEK cells (sham-treated) and to compare compound uptake into LLO- vs sham-treated cells (intracellular values used only). Likewise, intracellular quantification values were used to compare probe uptake into AAPH- vs sham-treated cells. For all resulting plots, one data point represents one replicate (mean over three fields of view) unless noted otherwise.

#### 5.5 Ferroptosis sensing, isolated mouse lung lymphocytes

**Mice**

C57BL/6 (WT) were bred in-house and maintained at in-house facilities. All mice were kept under specified pathogen free (SPF) conditions. All procedures were performed according to ethical protocols approved by the local and regional ethics committees.

**Whole lymphocyte culture**

Lungs were isolated, diced and digested with Liberase TL (0.25 mg/mL, Roche) and DNase I (1 mg/mL, Sigma) for 1 h at 37 °C. Lung cells were purified using a 37.5% Percoll gradient and red blood cell lysis was performed with Ammonium-Chloride-Potassium (ACK) lysing buffer. The single cell suspension was plated in 96-well U bottom plates with 3 x $10^5$ cells per well in RPMI medium supplemented with 3% FCS. Cells were then treated with RSL3, $H_2O_2$, or left untreated (PBS control) for 24 h at 37 °C.

**Flow cytometry**

To probe for signal from iPS-FS$_2$ or BODIPY, total lung lymphocytes were incubated with iPS-FS$_2$ (50 µM in FCS-free RPMI) for 30 min at 37 °C or with BODIPY- C11 (RED) (250 µM in FCS-free RPMI) (Image-iT® Lipid Peroxidation Sensor Kit, Fisher Scientific) for 1 h at 37 °C. To stain for surface markers fluorochrome-conjugated antibodies against the following surface antigens were used: CD45, CD11b, CD11c, Ly6G, Ly6C, SiglecF, F4/80, CD64, CD19, TCRb, CD4, CD8, CD44, NK1.1. DAPI (1:10000 from 5 mg/mL stock; at 4 °C for 20 min) was used to exclude dead cells. After washing the cells were resuspended in 50 µL FACS buffer and subsequently analysed using a BD LSR Fortessa.

#### 5.6 Necrosis sensing, live fly embryo

*; e22c-gal4, uas-mcherry-moesin Drosophila* embryos[12,13] were collected in cell strainers (Falcon), dechorionated with bleach (Jangro), and washed with water. Embryos were mounted ventral side up on scotch tape stuck down on a glass slide, then dechorionated in box with silica beads for 25-30 min. A droplet of VOLTALEF oil (VWR) was added to each embryo, following which **iPS-FS$_2$** (10 mM, until pressure reached 300 units) was microinjected into the intervitelline space surrounding the head of the embryo using an InjectMan4® microinjector (Eppendorf) and a FemtoJet injectman rig (Eppendorf) fitted with Femto tips (Eppendorf). A bridging glass cover slip was sealed over the top of the embryos, supported by two coverslips either side of the embryos. These were imaged live on an inverted spinning disc confocal microscope (Perkin Elmer Ultraview) with a plan-apochromat 63× objective (NA 1.4) and a Hamamatsu C9100-14 camera, and Volocity acquisition software (Perkin Elmer), with probe activation acquired in the green channel and mCherry-moesin in the red channel. Epithelial wounds were generated using a nitrogen-pumped Micropoint ablation laser tuned to 435 nm (single photon mode, focused within the epithelial layer at the area indicated by the asterisk; Andor Technologies). All images shown are z-projections.

The full imaging time-course is shown as the channel merge in **Movie S1**, with annotated still image zooms in **Fig 6**, and with larger resolution images of the first five minutes in **Fig S12**. For further details see the full legend to **Fig 6**, given in **Supporting Note 3**.

### 6 Chemistry materials and methods

#### 6.1 Analytical methods and techniques

##### 6.1.1 Analytical methods

High resolution mass spectrometry (**HRMS**) was conducted using two different instruments: (1) a *Thermo Finnigan LTQ FT Ultra FourierTransform* ion cyclotron resonance spectrometer from *ThermoFisher Scientific GmbH* applying electron spray ionisation (ESI) with a spray capillary voltage of 4 kV at temperature 250 °C with a method dependent range from 50 to 2000 u and (2) a *Finnigan MAT 95 from Thermo Fisher Scientific* applying electron ionisation (EI) at a source temperature of 250 °C and an electron energy of 70 eV with a method dependent range from 40 to 1040 u.

Nuclear magnetic resonance (**NMR**) spectroscopy was performed using different instruments: (1) a *Bruker Avance* (600/150 MHz, with TCI cryoprobe) or (2) a *Bruker Avance III HD Biospin* (400/100 MHz, with BBFO cryoprobe™) from Bruker Corp. either at 400 MHz or 500 MHz or (3) a Bruker Avance III HD (800 MHz, with cryoprobe). NMR-spectra were measured at 298 K, unless stated otherwise, and were analysed with the program *MestreNova 12* developed by *MestreLab Ltd.* $^{1}$H-NMR spectra chemical shifts (δ) in parts per million (ppm) relative to tetramethylsilane (δ = 0 ppm) are reported using the residual protic solvent ($CHCl_3$ in $CDCl_3$ : δ = 7.26 ppm, DMSO-$d_5$ in DMSO-$d_6$: δ = 2.50 ppm, $CHD_2OD$ in $CD_3OD$: δ = 3.31 ppm) as an internal reference. For $^{13}$C-NMR spectra, chemical shifts in ppm relative to tetramethylsilane (δ = 0 ppm) are reported using the central resonance of the solvent signal ($CDCl_3$: δ = 77.16 ppm, DMSO-$d_6$: δ = 39.52 ppm, $CD_3OD$: δ = 49.00 ppm) as an internal reference. For $^{1}$H-NMR spectra in addition to the chemical shift the following data is reported in parenthesis: multiplicity, coupling constant(s) and number of hydrogen atoms. The abbreviations for multiplicities and related descriptors are s = singlet, d = doublet, t = triplet, q = quartet, or combinations thereof, m = multiplet and br = broad. When rotamers were observed in the NMR spectra, the corresponding signals are separated by a slash ("/"). Where known products matched literature analysis data, only selected data acquired are reported.

Analytical high performance liquid chromatography (**HPLC**) analysis was conducted either using an *Agilent 1100* system from *Agilent Technologies Corp.,* Santa Clara (USA) equipped with a DAD detector and a *Hypersil Gold* HPLC column from *ThermoFisher Scientific GmbH*, Dreieich (Germany) or a *Agilent 1200 SL* system *Agilent Technologies Corp.,* Santa Clara (USA) equipped with a DAD detector, a *Hypersil Gold* HPLC column from *ThermoFisher Scientific GmbH*, Dreieich (Germany) and consecutive low-resolution mass detection using a LC/MSD IQ mass spectrometer applying ESI from *Agilent Technologies Corp.,* Santa Clara (USA). For both systems mixtures of water (analytical grade, 0.1 % formic acid) and MeCN (analytical grade, 0.1 % formic acid) were used as eluent systems.

**UV-Vis** spectra were recorded on an Cary 60 UV-Vis spectrophotometer from *Agilent Technologies Inc.*, Santa Clara (USA) using 1 cm quartz or PMMA cuvettes. The scan rate was set to 600 nm/min and 2.5 nm slit width was used. Unless stated otherwise, the probes and fluorophores were dissolved in PBS (pH = 7.4, 1 % DMSO) at 10 µM concentration.

**Fluorescence spectroscopy** was performed on a Cary Eclipse Fluorescence Spectrometer from *Agilent Technologies Inc.*, Santa Clara (USA) using quartz cuvettes. 480 nm light was used for excitation and spectra were recorded from 490−620 nm. The scan rate was set to 120 nm/min and 5 nm slit width was used. Unless stated otherwise, the probes and fluorophores were dissolved in PBS (pH = 7.4, 1 % DMSO) at 10 µM concentration.

##### 6.1.2 Synthetic techniques

Unless stated otherwise, all reactions were performed without precautions in regard to potential air- and moisture-sensitivity and were stirred with Teflon-coated magnetic stir bars. For work under inert gas (nitrogen) atmosphere, a Schlenk apparatus equipped with a liquid nitrogen trap and a high vacuum pump from Vacuubrand GmbH, Wertheim (Germany) were used. For solvent evaporation a *Laborota 400* from Heidolph GmbH, Schwabach (Germany) equipped with a vacuum pump was used. Flash column chromatography was conducted with a Biotage® Isolera One Chromatograph with Biotage® Sfär Silica D columns (10 g or 25 g silica) for normal-phase (np) chromatography or with Biotage® Sfär C18 D columns (12 g or 30 g silica) or Phenomenex AQ C18 spherical 20-35 µm columns (12 g) for reversed-phase (rp) chromatography. Reactions were monitored by thin layer chromatography (TLC) on TLC plates (*Si 60 F254 on aluminium sheets*) provided by Merck GmbH and visualised by UV irradiation and by analytical HPLC.

**Chemicals**

All chemicals, which were obtained from Sigma-Aldrich, TCI, Alfa Aesar, Acros, abcr or carbolution were used as received and without purification. Tetrahydrofuran (THF), dichloromethane (DCM) and dimethylformamide (DMF) were provided by Acros and were stored under argon atmosphere and dried over molecular sieves. TLC control, extractions and column chromatography were conducted using distilled, technical grade solvents. Whenever the term *hexanes* is used, the applied solvent actually comprised isomeric mixtures of hexane (2-methylpentane, 3-methylpentane, 2,2-dimethylbutane, 2,3-dimethylbutane).

#### 6.2 Synthetic procedures

##### 6.2.1 Literature procedures

The following molecules were synthesised according to literature procedures:

- **$i_2$-$FS_0$**[14]
- 4-(2-iodoethyl)morpholine[15]
- MSS00·HCl[16]

*Synthetic Note:* The mono-capped probes were synthesised by a route designed for simplicity and to avoid chromatography (**Fig 3b**). For example, **$H_2$-$FS_1$** was doubly alkylated at both the carboxylate and the phenol, then the ester was cleaved mildly with LiOH in THF/water at room temperature, to return the mono-alkylated fluorophores (e.g. **H-Me-$FS_1$, Fig 3b**); using typical conditions such as sodium hydroxide in methanol required heating and resulted in significant decomposition of the fluorescein core to the corresponding benzophenones. These mono-alkylated fluorophores were capped with isobutyric anhydride, affording the probes in good yield after the first and only column chromatography (e.g. **iMe-$FS_1$**, 58% over three steps, **Fig 3b**). Pure reference samples of uncapped, mono-alkylated fluorophores (e.g. **H-Me-$FS_1$**) could also be obtained by re-cleaving these probes with sodium hydroxide at room temperature within five minutes.

##### 6.2.2 Symmetric Fluorophores and Probes [non- or bis-acylated]

**5-Sulfo-2′,7′-dichlorofluorescein ($H_2$-$FS_1$)**

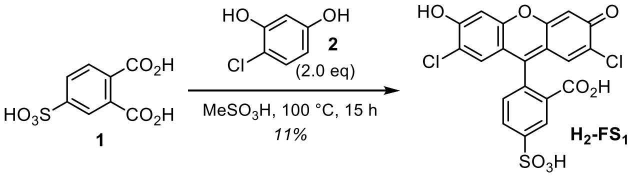

**$H_2$-$FS_1$** was prepared adapting a previously described procedure.[17] Sodium hydroxide (2M, aq., 50 mL, 100 mmol, 3.3 eq) was added to 4-sulfophthalic acid (**1**, 50% in water, 11.4 mL, 30.0 mmol, 1.0 eq) until the solution was basic (checked by pH paper). The mixture was evaporated affording a purple solid which was dissolved in methanesulfonic acid (60 mL). 4-Chlororesorcinol (**2**, 8.67 g, 60.0 mmol, 2.0 eq) was added. The reaction mixture was heated to 100 °C for 15 h, poured into ice water (200 mL) and stirred at r.t. for 15 min. The suspension was filtered, and the wet precipitate was suspended in acetone (130 mL) and heated to reflux for 2 h. The suspension was filtered affording regioisomerically pure **$H_2$-$FS_1$** (1.51 g, 3.14 mmol, 11%) as orange solid (isomeric ratio >20:1). *Note*: *the acetone filtrate can be evaporated to afford a regioisomeric mixture of 5- and 6-sulfofluoresceins. Filtering the precipitated reaction mixture directly without acetone suspension increases the yield to 68% (isomeric mixture 1:1).*

**TLC** $R_f$ = 0.70 (*rp,* 40% MeCN); **$^1$H-NMR** (400 MHz, DMSO-$d_6$): $\delta$ (ppm) = 11.10 (s, br, 1H), 8.08 (s, 1H), 8.01 (d, J = 7.7 Hz, 1H), 7.28 (d, J = 7.8 Hz, 1H), 6.90 (s, 2H), 6.71 (s, 2H); **$^{13}$C-NMR** (101 MHz, DMSO-$d_6$): $\delta$ (ppm) = 167.95, 155.19, 151.59, 150.61, 150.03, 133.23, 128.37, 125.74, 123.79, 121.56, 116.35, 110.25, 103.68, 81.40; **HRMS** (ESI−): m/z calc. for $C_{20}H_9Cl_2O_8S^-$ $[M-H]^-$: 478.9401, found: 478.9398.

**5-Sulfo-2′,7′-dichlorofluorescein diacetate ($a_2$-$FS_1$)**

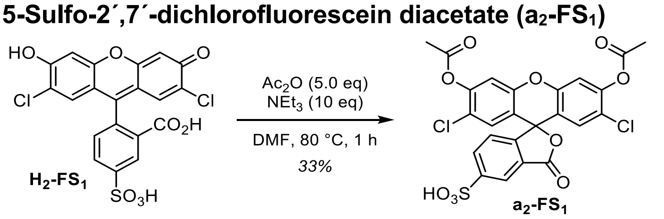

**$a_2$-$FS_1$** was prepared adapting a previously described procedure.[17] **$H_2$-$FS_1$** (100 mg, 208 µmol, 1.0 eq) was dissolved in dry DMF (5 mL) and triethylamine (0.37 mL, 2.08 mmol, 10 eq) and acetic anhydride (0.10 mL, 1.04 mmol, 5.0 eq) were added. The reaction mixture was heated to 80 °C for 30 min, then the volatiles were removed *in vacuo*. The crude product was purified by column chromatography (reversed phase, 5→40% MeCN (0.1% FA)). From the product column fractions the acetonitrile was removed *in vacuo* (*Note*: 40 °C water bath, not higher to avoid probe hydrolysis) and the remaining water was lyophilised over night to afford **$a_2$-$FS_1$** (39 mg, 69.0 µmol, 33%) as light yellow solid.

**TLC** $R_f$ = 0.48 (*rp,* 40% MeCN); **$^1$H-NMR** (400 MHz, DMSO-$d_6$): $\delta$ (ppm) = 8.13 (s, 1H), 8.02 (d, J = 7.8 Hz, 1H), 7.57 (s, 2H), 7.44 (d, J = 7.8 Hz, 1H), 7.14 (s, 2H), 2.35 (s, 6H); **$^{13}$C-NMR** (101 MHz, DMSO-$d_6$): $\delta$ (ppm) = 168.05, 167.67, 151.28, 150.87, 149.28, 148.29, 133.38, 128.83, 125.39, 123.92, 122.03, 121.94, 117.59, 113.26, 79.71, 20.42; **HRMS** (ESI−): m/z calc. for $C_{24}H_{13}Cl_2O_{10}S^-$ $[M-H]^-$: 562.9612, found: 562.9624.

###### 5-Sulfo-2´,7´-dichlorofluorescein diisobutyrate ($i_2$-FS$_1$)

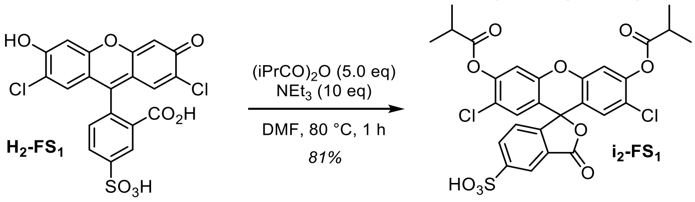

**$i_2$-FS$_1$** was prepared adapting a previously described procedure.[17] **$H_2$-FS$_1$** (2.19 g, 4.55 mmol, 1.0 eq) was dissolved in dry DMF (25 mL) and TEA (7.74 mL, 45.5 mmol, 10 eq) and isobutyric anhydride (15.6 mL, 91.0 mmol, 5.0 eq) were added. The reaction mixture was heated to 80 °C for 1 h, then the volatiles were removed *in vacuo*. The crude product was purified by column chromatography (reversed phase, 5→50% MeCN (0.1% FA)). To the product fractions, sodium hydrogencarbonate was added until pH = 7 was reached, then DCM was added, the layers were separated and the aq. layer was extracted with DCM. The combined organic layers were dried over sodium sulfate, filtered and concentrated affording **$i_2$-FS$_1$** (1.38 g, 3.69 mmol, 81%) as colourless solid.

**TLC** $R_f$ = 0.22 (*rp,* 40% MeCN); **$^1$H-NMR** (400 MHz, DMSO-d$_6$): $\delta$ (ppm) = 8.13 (d, J = 1.7 Hz, 1H), 8.03 (dd, J = 8.0, 1.5 Hz, 1H), 7.57 (s, 2H), 7.44 (d, J = 8.0 Hz, 1H), 7.15 (s, 2H), 2.91 (hept, J = 7.0 Hz, 2H), 1.28 (d, J = 7.0 Hz, 12H); **$^{13}$C-NMR** (101 MHz, DMSO-d$_6$): $\delta$ (ppm) = 174.17, 168.13, 151.71, 151.34, 149.76, 148.72, 133.83, 129.31, 125.83, 124.31, 122.48, 122.40, 118.00, 113.67, 80.15, 33.77, 19.07; **HRMS** (ESI−): m/z calc. for $C_{28}H_{21}Cl_2O_{10}S^-$ [M−H]$^-$: 619.0238, found: 619.0248.

##### 6.2.3 General Procedure for mono-capped probes and fluorophores

The mono-capped probes were synthesised in three steps from **$H_2$-FS$_1$** by alkylation with the respective halide, ester cleavage with lithium hydroxide and capping with isobutyric anhydride (adapted from Woodroofe *et al.*[17]). The pure fluorophores were obtained by cleaving purified probe with sodium hydroxide in methanol/water.

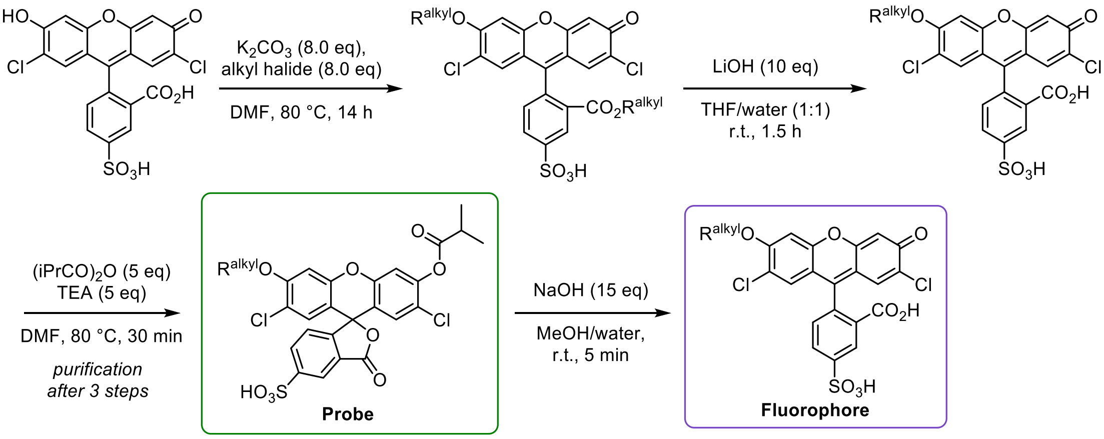

###### General Procedure A: mono-capped probes

**$H_2$-FS$_1$** (1.0 eq) was dissolved in anhydrous DMF (0.01−0.04 M). Potassium carbonate (8.0 eq) and the respective **alkyl halide** (8.0 eq) were added and the reaction was heated 80 °C for 2.5−21 h until full conversion to the dialkyl-product (monitored by HPLC/MS). The solvent was removed *in vacuo* and the crude product was dissolved in THF/water (~0.02 M). Lithium hydroxide (10 eq) was added and the reaction mixture was stirred at r.t. for 1.5 h. Then aqueous hydrochloric acid (2 M, 12 eq) was added to acidify the solution (checked by pH paper) and the volatiles were removed *in vacuo* affording the crude alkylated fluorophore which was dissolved in DMF (~0.02 M). Triethylamine (5.0 eq) and isobutyric anhydride (5.0 eq) were added and the reaction mixture was heated to 80 °C for 30 min. The volatiles were removed *in vacuo* and the crude product was purified by reversed-phase flash column chromatography (acetonitrile/water, 0.1% FA). From the column fractions containing the product, the acetonitrile was removed at the rotary evaporator (bath temperature: 40 °C to avoid probe hydrolysis), then the aqueous solution was lyophilised overnight.

<u>Notes</u>: (1) If the probe can not be afforded in the desired purity by this procedure, we recommend capping the purified fluorophores (see General Procedure B). (2) For **iPS-FS$_2$** this procedure was adapted due to lower solubility in organic solvents.

**General Procedure B: mono-alkylated fluorophores**

The corresponding probe (1.0 eq) was dissolved in methanol (~0.02 M). Sodium hydroxide (aqueous, 2 M, 15 eq) was added and the reaction mixture was stirred at r.t. for 5 min. Hydrochloric acid (aqueous, 2 M, 20 eq) was added to acidify the solution and the volatiles were removed *in vacuo*. The crude product was purified by reversed-phase flash column chromatography (acetonitrile/water, 0.1% FA). The solvent of the product fractions was removed *in vacuo*.

Notes: We found that the purification of the fluorophores was simplest to perform by preparing the probes in three steps only purifying the product and then cleaving the probe to the fluorophore again.

##### 6.2.4 7'-*O*-alkylated-2'-*O*-acylated [mono-capped] fluorogenic probes

**iMe-FS$_1$**

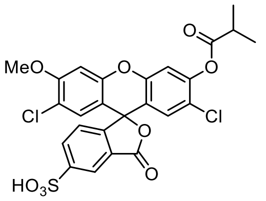

Prepared according to General Procedure A from **H$_2$-FS$_1$** (60 mg, 125 µmol, 1.0 eq) and methyl iodide (62 µL, 997 µmol, 8.0 eq). Reaction time for alkylation: 14 h. Ester cleavage in THF/water = 1:1. Purification of the probe by rp-column chromatography: 10→50% MeCN.

The product **iMe-FS$_1$** (41.0 mg, 72.5 µmol, 58% (3 steps)) was obtained as light yellow solid.

**TLC** $R_f$ = 0.33 (*rp,* 40% MeCN); **$^1$H-NMR** (400 MHz, DMSO-d$_6$): $\delta$ (ppm) = 8.12 (d, J = 11.6 Hz, 1H), 8.01 (dd, J = 8.0, 1.2 Hz, 1H), 7.52 (s, 1H), 7.35 (d, J = 8.0 Hz, 1H), 7.19 (s, 1H), 7.12 (s, 1H), 6.89 (s, 1H), 3.96 (s, 3H), 2.99 – 2.84 (m, 1H), 1.27 (d, J = 7.0 Hz, 6H); **$^{13}$C-NMR** (101 MHz, DMSO-d$_6$): $\delta$ (ppm) = 174.19, 168.24, 156.92, 151.90, 151.27, 150.48, 150.00, 148.61, 133.78, 129.33, 128.67, 125.93, 124.22, 122.29, 122.16, 118.19, 117.99, 113.52, 111.21, 101.78, 80.73, 57.37, 33.77, 19.10; **HRMS** (ESI−): m/z calc. for $C_{25}H_{17}Cl_2O_9S^-$ [M−H]$^-$: 562.9975, found: 562.9979.

**iEM-FS$_1$**

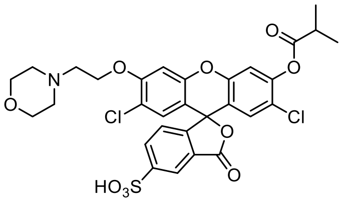

Prepared according to General Procedure A from **H$_2$-FS$_1$** (80 mg, 166 µmol, 1.0 eq) and 4-(2-iodoethyl)morpholine (321 mg, 1.33 mmol, 8.0 eq). Reaction time for alkylation: 16 h. Ester cleavage in THF/water = 1:2. Purification of the probe by rp-column chromatography: 5→60% MeCN.

The product **iMe-FS$_1$** (56.0 mg, 84.3 µmol, 51% (3 steps)) was obtained as off-white solid.

**TLC** $R_f$ = 0.33 (*rp,* 40% MeCN); **$^1$H-NMR** (400 MHz, DMSO-d$_6$): $\delta$ (ppm) = 9.86 (s, br, 1H), 8.12 – 8.10 (m, 1H), 8.02 (dd, *J* = 8.0, 1.5 Hz, 1H), 7.52 (s, 1H), 7.35 – 7.30 (m, 2H), 7.17 (s, 1H), 6.95 (s, 1H), 4.56 (s, 2H), 3.99 (s, 2H), 3.66 (s, 8H), 2.92 (hept, J = 7.0 Hz, 1H), 1.27 (d, J = 7.0 Hz, 6H); **$^{13}$C-NMR** (101 MHz, DMSO-d$_6$): $\delta$ (ppm) = 174.19, 168.20, 155.25, 151.88, 151.28, 150.30, 149.90, 148.63, 133.80, 129.43, 128.85, 125.87, 124.08, 122.36, 122.29, 118.25, 118.11, 113.47, 112.21, 102.94, 80.53, 64.85, 63.92, 63.82, 52.66, 33.77, 19.07; **HRMS** (ESI+): m/z calc. for $C_{30}H_{28}Cl_2NO_{10}S^+$ [M+H]$^+$: 664.0805, found: 664.0809.

**iPS-FS$_1$**

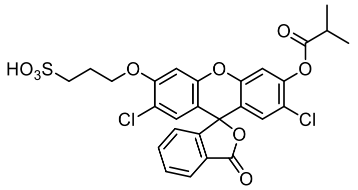

Prepared according to General Procedure A from **H$_2$-FS$_0$** (60 mg, 150 µmol, 1.0 eq) and sodium 3-bromo-propane-1-sulfonate (269 mg, 1.20 mmol, 8.0 eq). Reaction time for alkylation: 21 h. Ester cleavage in THF/water = 1:4. Purification of the probe by rp-column chromatography: 10→60% MeCN.

The product **iPS-FS$_1$** (21.0 mg, 42.1 µmol, 28% (3 steps)) was obtained as light orange solid.

**TLC** $R_f$ = 0.36 (*rp,* 40% MeCN); **$^1$H-NMR** (400 MHz, DMSO-d$_6$): *δ* (ppm) = 8.04 (d, J = 7.5 Hz, 1H), 7.83 (t, *J* = 7.2 Hz, 1H), 7.77 (t, *J* = 7.4 Hz, 1H), 7.53 (s, 1H), 7.42 (d, *J* = 7.5 Hz, 1H), 7.17 (s, 1H), 7.01 (s, 1H), 6.81 (s, 1H), 4.29 (m, 2H), 2.91 (hept, *J* = 7.0 Hz, 1H), 2.60 (m, 2H), 2.06 (m, 2H), 1.26 (d, *J* = 7.0 Hz, 6H); **$^{13}$C-NMR** (101 MHz, DMSO-d$_6$): *δ* (ppm) = 173.78, 168.21, 155.76, 151.46, 150.08, 149.68, 148.13, 136.13, 130.80, 128.63, 128.08, 125.58, 125.40, 124.08, 121.63, 117.99, 117.76, 113.20, 110.83, 101.95, 80.43, 68.54, 47.71, 33.35, 24.95, 18.66; **HRMS** (ESI+): m/z calc. for $C_{27}H_{23}Cl_2O_9S^+$ [M+H]$^+$: 593.0434, found: 593.0430.

**iPS-FS$_2$**

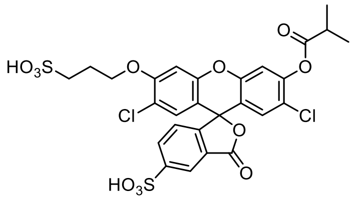

Prepared according to General Procedure A from **H$_2$-FS$_1$** (400 mg, 831 µmol, 1.0 eq) and 1,3-propanesultone (812 mg, 6.65 mmol, 8.0 eq) and heated to 100 °C for 7 h. Ester cleavage in water (no THF). Purification of the probe by rp-column chromatography: 5→40% MeCN.

The product **iPS-FS$_2$** (136 mg, 0.202 mmol, 24% (3 steps)) was obtained as light yellow solid.

**TLC** $R_f$ = 0.62 (*rp,* 40% MeCN); **$^1$H-NMR** (400 MHz, D$_2$O/DMSO-d$_6$ 2:1): *δ* (ppm) = 8.24 (s, 1H), 8.06 (d, J = 8.1 Hz, 1H), 7.33 – 7.27 (m, 2H), 7.04 (s, 1H), 6.97 (s, 1H), 6.82 (s, 1H), 4.14 (t, J = 6.0 Hz, 2H), 2.89 – 2.75 (m, 3H), 2.15 – 2.05 (m, 2H), 1.17 (d, J = 7.0 Hz, 6H).; **$^{13}$C-NMR** (101 MHz, D$_2$O/DMSO-d$_6$ 2:1): *δ* (ppm) = 177.56, 170.45, 157.37, 154.59, 151.69, 151.26, 149.79, 148.55, 134.98, 130.07, 129.64, 127.26, 126.08, 125.53, 124.00, 123.57, 119.76, 118.30, 111.30, 103.50, 83.48, 69.39, 49.12, 35.08, 25.61, 19.78; **HRMS** (ESI−): m/z calc. for $C_{27}H_{21}Cl_2O_{12}S_2^-$ [M−H]$^-$: 670.9857, found: 670.9864.

#### 6.2.5 2'-*O*-[mono]alkylated fluorophores

**H-Me-FS$_1$**

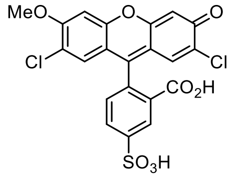

Prepared according to General Procedure B from **iMe-FS$_1$** (26 mg, 46.4 µmol, 1.0 eq). Purification by rp-column chromatography: 5→40% MeCN.

The product **H-Me-FS$_1$** (14 mg, 28.3 µmol, 61%) was obtained as orange solid.

**TLC** $R_f$ = 0.48 (*rp,* 40% MeCN); **$^1$H-NMR** (400 MHz, DMSO-d$_6$): *δ* (ppm) = 11.11 (s, br, 1H), 8.08 (d, J = 1.4 Hz, 1H), 8.00 (dd, J = 7.9, 1.5 Hz, 1H), 7.28 (d, J = 7.9 Hz, 1H), 7.20 (s, 1H), 6.93 (s, 1H), 6.83 (s, 1H), 6.75 (s, 1H), 3.94 (s, 3H); **$^{13}$C-NMR** (101 MHz, DMSO-d$_6$): *δ* (ppm) = 168.39, 156.73, 155.68, 152.09, 151.12, 150.81, 150.35, 133.71, 128.87, 128.59, 126.08, 124.18, 122.04, 117.49, 116.96, 111.51, 110.58, 104.02, 101.84, 81.63, 57.29; **HRMS** (ESI−): m/z calc. for $C_{21}H_{11}Cl_2O_8S^-$ [M−H]$^-$: 492.9557, found: 492.9559.

**H-EM-FS$_1$**

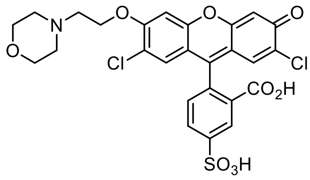

Prepared according to General Procedure B from **iEM-FS$_1$** (56 mg, 84.3 µmol, 1.0 eq). Purification by rp-column chromatography: 5→40% MeCN.

The product **H-EM-FS$_1$** (42 mg, 70.7 µmol, 84%) was obtained as red solid.

**TLC** $R_f$ = 0.48 (*rp,* 40% MeCN); **$^1$H-NMR** (400 MHz, DMSO-d$_6$): $\delta$ (ppm) = 11.11 (s, 1H), 9.89 (s, br, 1H), 8.09 (d, J = 1.4 Hz, 1H), 8.02 (dd, J = 8.0, 1.5 Hz, 1H), 7.33 (s, 1H), 7.24 (d, J = 8.0 Hz, 1H), 6.93 (s, 1H), 6.88 (s, 1H), 6.79 (s, 1H), 4.55 (s, 2H), 3.99 (s, 2H), 3.67 (s, 8H); **$^{13}$C-NMR** (101 MHz, DMSO-d$_6$): $\delta$ (ppm) = 167.90, 155.28, 154.71, 151.62, 150.71, 150.22, 149.82, 133.30, 128.52, 128.37, 125.60, 123.59, 121.69, 117.31, 116.64, 112.04, 110.05, 103.54, 102.55, 80.97, 64.35, 63.45, 54.79, 52.19; **HRMS** (ESI−): m/z calc. for $C_{26}H_{20}Cl_2NO_9S^-$ [M−H]$^-$: 592.0241, found: 592.0244.

**H-PS-FS$_1$**

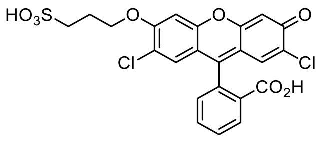

Prepared according to General Procedure B from **iPS-FS$_1$** (19 mg, 32.0 µmol, 1.0 eq). Purification by rp-column chromatography: 10→40% MeCN.

The product **H-PS-FS$_1$** (16 mg, 30.6 µmol, 95%) was obtained as orange solid.

**TLC** $R_f$ = 0.59 (*rp,* 40% MeCN); **$^1$H-NMR** (400 MHz, methanol-d$_4$): $\delta$ (ppm) = 11.13 (s, br, 1H), 8.02 (d, J = 7.5 Hz, 1H), 7.82 (t, J = 7.4 Hz, 1H), 7.75 (t, J = 7.4 Hz, 1H), 7.36 (d, J = 7.6 Hz, 1H), 7.18 (s, 1H), 6.94 (s, 1H), 6.75 (s, 1H), 6.67 (s, 1H), 4.27 (t, J = 6.3 Hz, 2H), 2.59 (t, J = 7.3 Hz, 2H), 2.10 – 1.98 (m, 2H); **$^{13}$C-NMR** (101 MHz, methanol-d$_4$): $\delta$ (ppm) = 168.32, 155.53, 155.19, 151.59, 150.39, 150.02, 135.98, 130.58, 128.24, 128.01, 125.81, 125.14, 124.03, 117.25, 116.38, 111.15, 110.36, 103.65, 101.98, 81.29, 68.41, 47.66, 24.93; **HRMS** (ESI−): m/z calc. for $C_{23}H_{15}Cl_2O_8S^-$ [M−H]$^-$: 520.9870, found: 520.9878.

**H-PS-FS$_2$**

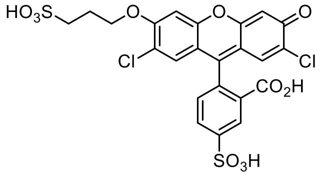

Prepared according to General Procedure B from **iPS-FS$_2$** (69 mg, 103 µmol, 1.0 eq). Purification by rp-column chromatography: 2→10% MeCN.

The product **H-PS-FS$_2$** (24 mg, 39.8 µmol, 39%) was obtained as orange solid.

**TLC** $R_f$ = 0.85 (*rp,* 40% MeCN); **$^1$H-NMR** (400 MHz, $D_2O$): $\delta$ (ppm) = 8.49 (s, 1H), 8.15 (d, J = 8.7 Hz, 1H), 7.45 (d, J = 8.1 Hz, 1H), 6.89 (s, 1H), 6.83 (s, 1H), 5.69 (s, 1H), 5.63 (s, 1H), 3.42 – 3.24 (m, 2H), 2.95 – 2.85 (m, 2H), 2.16 – 1.98 (m, 2H).; **$^{13}$C-NMR** (101 MHz, $D_2O$): $\delta$ (ppm) = 169.62, 155.56, 155.21, 152.46, 150.01, 149.98, 145.47, 133.29, 128.31, 128.18, 127.03, 125.09, 123.42, 118.19, 117.65, 109.78, 109.70, 102.71, 99.97, 91.14, 67.33, 47.53, 23.98; **HRMS** (ESI−): m/z calc. for $C_{23}H_{15}Cl_2O_{11}S_2^-$ [M−H]$^-$: 600.9438, found: 600.9443.

##### 6.2.6 Reducible disulfide probe

**MSS00-PS-FS₂**

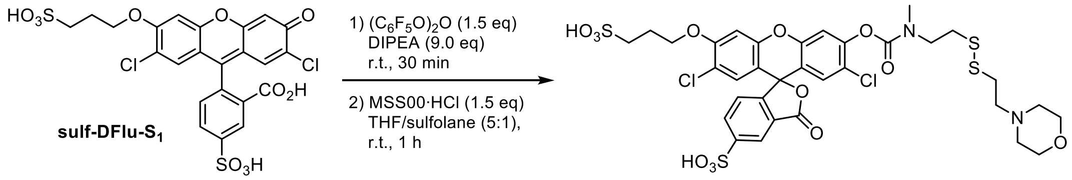

**H-PS-FS$_2$** (10 mg, 16.6 µmol, 1.0 eq) was dissolved in THF (3.0 mL) and sulfolane (0.6 mL). DIPEA (25 µL, 149 µmol, 9.0 eq) and bis(pentafluorophenyl) carbonate (10 mg, 24.9 µmol, 1.5 eq) were added and the reaction mixture was stirred at r.t. for 30 min. The redox trigger **MSS00·HCl** (13 mg, 41.5 µmol, 2.5 eq) was added and the reaction was stirred at r.t. for 1 h. The volatiles were removed *in vacuo* and the crude product was purified by reversed-phase-AQ flash column chromatography (5→30% MeCN, acetonitrile/water, 0.1% FA). The product (4.8 mg, 5.57 µmol, 34%) was obtained as colourless solid.

**TLC** $R_f$ = 0.72 (*rp,* 40% MeCN); **$^1$H-NMR** (800 MHz, DMSO-d$_6$) δ 8.20 (s, 1H), 8.07 (d, J = 7.9 Hz, 1H), 7.39/7.34 (2 × s, 1H), 7.32 (d, J = 7.6 Hz, 1H), 7.12/7.11 (2 × s, 1H), 6.94 (s, 1H), 6.82 (s, 1H), 4.21 – 4.17 (m, 2H), 3.97 – 3.85 (m, 2H), 3.77 – 3.67 (m, 1H), 3.66 – 3.55 (m, 3H), 3.41 – 3.31 (m, 4H), 3.03/2.91 (2 × s, 3H), 3.02 – 2.92 (m, 6H), 2.77 – 2.73 (m, 2H), 2.09 (p, J = 6.6 Hz, 2H).; **$^{13}$C-NMR** (201 MHz, DMSO-d$_6$) δ 168.62, 156.16, 153.51/153.11, 152.72/152.67, 150.44, 149.86, 149.16/148.97, 148.85, 133.94, 128.72/128.64, 128.39, 125.93/125.91, 124.59, 122.57, 122.46, 118.40, 117.05, 113.67/113.40, 110.39, 102.28, 81.57, 68.51, 63.61, 55.63/55.54/55.38/51.70, 48.06, 47.93, 47.04, 36.17, 35.38/35.22, 35.15/34.83/32.93/32.52, 30.42/30.30, 26.22, 24.68.; **HRMS** (ESI−): m/z calc. for $C_{33}H_{33}Cl_2N_2O_{13}S_4^-$ [M−H]$^-$: 863.0248, found: 863.0242.

##### 6.2.7 Esterase-labile fluorogenic probe for attachment onto macromolecules

**Dichlorofluorescein isobutyrate NHS-ester (NHS-i-Flu)**

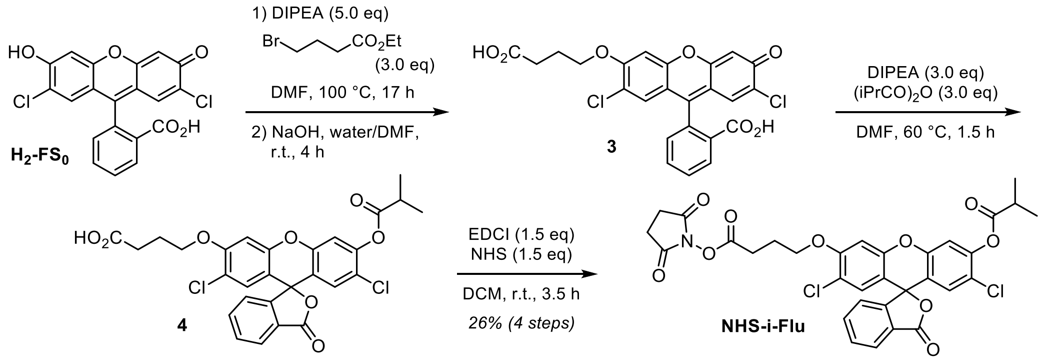

2′,7′-Dichlorofluorescein (**$H_2$-$FS_0$**, 600 mg, 1.50 mmol, 1.0 eq) was dissolved in dry DMF (20 mL). DIPEA (1.27 mL, 7.48 mmol, 5.0 eq) and ethyl 4-bromobutyrate (0.64 mL, 4.49 mmol, 3.0 eq) were added and the reaction mixture was heated to 100 °C for 17 h. After allowing the reaction mixture to cool to r.t., aqueous sodium hydroxide solution (2M, 12 mL, 16 eq) was added and the mixture was stirred for another 4 h. Then hydrochloric acid (2M, 14 mL, 19 eq) was added and the volatiles were removed *in vacuo*.

The crude **3** was dissolved in dry DMF (14 mL), DIPEA (0.51 mL, 2.99 mmol, 3.0 eq) and isobutyric anhydride (0.50 mL, 2.99 mmol, 3.0 eq) were added and the reaction mixture was heated to 60 °C for 1.5 h. Water (120 mL) and DCM (120 mL) were added, the layers were separated and the aqueous layer was extracted with DCM (1 × 120 mL). The organic layer was dried over sodium sulfate, the desiccant was filtered off and the solvent removed *in vacuo*. Reversed-phase column chromatography (acetonitrile/water, 0.1% FA, 10→50% MeCN) was used to semi-purify **4**, which was directly used for the NHS-ester formation.

**4** was dissolved in DCM (10 mL) and *N*-hydroxysuccinimide (259 mg, 2.25 mmol, 1.5 eq) and EDCI (431 mg, 2.25 mmol, 1.5 eq) were added. The reaction mixture was stirred at r.t. for 3.5 h, then the reaction mixture was diluted with DCM (20 mL) and water (20 mL), the layers were separated and the aqueous layer was extracted with DCM (2 × 20 mL). The crude product was purified by column chromatography (iso-hexanes/EtOAc, 40→100% EtOAc). **NHS-i-Flu** (255 mg, 0.393 mmol, 26%) was obtained as colourless solid.

**TLC** $R_f$ = 0.36 (*np, iso*-hexanes/EtOAc); **$^1$H-NMR** (400 MHz, $CDCl_3$): $\delta$ (ppm) = 8.03 (d, J = 7.3 Hz, 1H), 7.75 – 7.62 (m, 2H), 7.15 (d, J = 7.5 Hz, 1H), 7.09 (s, 1H), 6.82 (s, 1H), 6.79 (s, 1H), 6.73 (s, 1H), 4.15 (t, J = 5.9 Hz, 2H), 2.91 – 2.78 (m, 7H), 2.27 (p, J = 6.7 Hz, 2H), 1.36 – 1.29 (m, 6H); **$^{13}$C-NMR**(101 MHz, $CDCl_3$): $\delta$ (ppm) = 174.18, 169.18, 168.75, 168.21, 155.77, 151.96, 150.41, 150.01, 148.40, 135.69, 130.53, 129.00, 128.68, 126.04, 125.52, 123.96, 122.34, 118.96, 117.71, 112.67, 111.25, 101.38, 81.16, 67.28, 34.12, 27.51, 25.60, 24.19, 18.95; **HRMS** (EI+): m/z calc. for $C_{32}H_{25}Cl_2NO_{10}^+$ $[M]^{\cdot+}$: 653.0856, found: 653.0855.

### 7 Raw Biological Assay Data

**Table S4**: Cellular fluorescence values quantified from microscopy images for the cell penetration into healthy HEK cells (n = 3).

| Compound | Intracellular fluorescence | | | Extracellular fluorescence | | | Mean (intra) | Mean (extra) | Ratio intra/extra |
|---|---|---|---|---|---|---|---|---|---|
| $i_2$-$FS_0$ | 411 | 517 | 296 | 4.68 | 2.88 | 3.75 | 408 | 3.77 | 108 |
| $i_2$-$FS_1$ | 319 | 209 | 168 | 4.095 | 20.2 | 8.43 | 232 | 10.9 | 21.3 |
| $a_2$-$FS_1$ | 165 | 122 | 187 | 9.28 | 16.5 | 13.0 | 158 | 12.9 | 12.2 |
| iPS-$FS_1$ | 83.3 | 84.4 | 69.7 | 3.516 | 3.022 | 3.878 | 79.1 | 3.47 | 22.8 |
| iMe-$FS_1$ | 4.35 | 5.42 | 4.93 | 2.31 | 2.59 | 3.69 | 4.90 | 2.86 | 1.71 |
| iEM-$FS_1$ | 4.36 | 4.25 | 6.17 | 1.54 | 1.56 | 2.16 | 4.93 | 1.75 | 2.81 |
| iPS-$FS_2$ | 1.89 | 2.85 | 1.59 | 1.91 | 1.73 | 1.84 | 2.11 | 1.83 | 1.15 |
| FDP | 9.44 | 21.2 | 14.5 | 10.4 | 8.71 | 11.2 | 15.0 | 10.1 | 1.49 |
| fluorescein | 356 | 391 | 318 | 1287 | 1444 | 1379 | 355 | 1370 | 0.26 |
| $H_2$-$FS_0$ | 209 | 274 | 236 | 715 | 854 | 874 | 240 | 814 | 0.29 |
| $H_2$-$FS_1$ | 231 | 155 | 192 | 509 | 562 | 504 | 193 | 525 | 0.37 |
| H-PS-$FS_1$ | 22.7 | 23.1 | 21.1 | 89.3 | 55.8 | 72.9 | 22.3 | 72.7 | 0.31 |
| H-Me-$FS_1$ | 12.6 | 17.4 | 14.8 | 59.5 | 68.7 | 52.7 | 15.0 | 60.3 | 0.25 |
| H-EM-$FS_1$ | 6.91 | 8.13 | 9.85 | 39.0 | 38.1 | 37.3 | 8.30 | 38.1 | 0.22 |
| H-PS-$FS_2$ | 8.95 | 16.9 | 31.4 | 59.1 | 58.1 | 96.4 | 19.1 | 71.2 | 0.27 |
| DMSO | 1.34 | 0.98 | 0.64 | 0.13 | 0.15 | 0.05 | 0.98 | 0.11 | 8.86 |

**Table S5**: Intracellular fluorescence values quantified from microscopy images for the LLO membrane damage assay (n = 3).

| Compound | +LLO | | | −LLO | | | Mean (+LLO) | Mean (−LLO) | Ratio +/−LLO |
|---|---|---|---|---|---|---|---|---|---|
| $i_2$-$FS_0$ | 125 | 172 | 208 | 411 | 517 | 296 | 169 | 408 | 0.41 |
| $i_2$-$FS_1$ | 4977 | 2079 | 3305 | 319 | 209 | 168 | 3454 | 232 | 14.9 |
| $a_2$-$FS_1$ | 1406 | 1646 | 1523 | 165 | 122 | 187 | 1525 | 158 | 9.67 |
| iPS-$FS_1$ | 299 | 256 | 261 | 83.3 | 84.4 | 69.7 | 272 | 79.1 | 3.44 |
| iMe-$FS_1$ | 131 | 149 | 163 | 4.35 | 5.42 | 4.93 | 148 | 4.90 | 30.2 |
| iEM-$FS_1$ | 160 | 177 | 194 | 4.36 | 4.25 | 6.17 | 177 | 4.93 | 36.0 |
| iPS-$FS_2$ | 70.90 | 63.98 | 63.17 | 1.89 | 2.85 | 1.59 | 66.02 | 2.11 | 31.3 |
| FDP | 114 | 85.82 | 97.07 | 9.44 | 21.18 | 14.47 | 99.05 | 15.0 | 6.59 |
| fluorescein | 1447 | 1624 | 1225 | 356 | 391 | 318 | 1432 | 355 | 4.03 |
| $H_2$-$FS_0$ | 1178 | 1351 | 1033 | 209 | 274 | 236 | 1187 | 240 | 4.96 |
| $H_2$-$FS_1$ | 723 | 772 | 650 | 231 | 155 | 192 | 715 | 193 | 3.71 |
| H-PS-$FS_1$ | 114 | 94.45 | 107 | 22.65 | 23.11 | 21.14 | 105 | 22.3 | 4.72 |
| H-Me-$FS_1$ | 79.6 | 83.1 | 59.6 | 12.6 | 17.4 | 14.8 | 74.1 | 15.0 | 4.96 |
| H-EM-$FS_1$ | 44.2 | 40.4 | 39.6 | 6.91 | 8.13 | 9.85 | 41.4 | 8.30 | 4.99 |
| H-PS-$FS_2$ | 66.5 | 73.8 | 97.3 | 8.95 | 16.9 | 31.4 | 79.2 | 19.1 | 4.15 |
| DMSO | 1.83 | 1.29 | 1.29 | 1.34 | 0.98 | 0.64 | 1.47 | 0.98 | 1.49 |

**Table S6**: Intracellular fluorescence values quantified from microscopy images for the AAPH membrane damage assay (n = 3).

| Compound | +AAPH | | | −AAPH | | | Mean (+AAPH) | Mean (−AAPH) | Ratio +/−AAPH |
|---|---|---|---|---|---|---|---|---|---|
| i$_2$-FS$_0$ | 49.5 | 22.4 | 13.3 | 608 | 450 | 293 | 28.4 | 450 | 0.06 |
| i$_2$-FS$_1$ | 678 | 778 | 314 | 2490 | 2523 | 1764 | 590 | 2259 | 0.26 |
| a$_2$-FS$_1$ | 369 | 824 | 369 | 230 | 484 | 317 | 520 | 344 | 1.51 |
| iPS-FS$_1$ | 77.7 | 90.3 | 52.5 | 134 | 245 | 147 | 73.5 | 175 | 0.42 |
| iMe-FS$_1$ | 54.5 | 29.5 | 46.1 | 32.2 | 32.9 | 27.2 | 43.4 | 30.8 | 1.41 |
| iEM-FS$_1$ | 59.3 | 110 | 59.6 | 25.6 | 32.7 | 19.5 | 76.2 | 25.9 | 2.94 |
| iPS-FS$_2$ | 38.5 | 43.3 | 25.1 | 3.75 | 5.06 | 4.90 | 35.6 | 4.57 | 7.80 |
| FDP | 19.3 | 12.5 | 22.7 | 5.27 | 3.90 | 7.11 | 18.2 | 5.43 | 3.35 |
| fluorescein | 870 | 852 | 903 | 242 | 236 | 310 | 875 | 263 | 3.33 |
| H$_2$-FS$_0$ | 1088 | 571 | 538 | 232 | 100 | 116 | 733 | 149 | 4.90 |
| H$_2$-FS$_1$ | 389 | 544 | 373 | 77.7 | 91.9 | 101 | 435 | 90.23 | 4.83 |
| H-PS-FS$_1$ | 115 | 95.9 | 53.0 | 21.6 | 24.3 | 21.8 | 88.1 | 22.6 | 3.90 |
| H-Me-FS$_1$ | 63.2 | 54.1 | 56.6 | 14.5 | 12.4 | 13.9 | 57.9 | 13.6 | 4.26 |
| H-EM-FS$_1$ | 34.2 | 27.4 | 34.2 | 9.58 | 10.2 | 8.74 | 31.9 | 9.50 | 3.36 |
| H-PS-FS$_2$ | 42.5 | 49.6 | 37.8 | 9.89 | 14.0 | 12.5 | 43.3 | 12.1 | 3.57 |
| DMSO | 0.94 | 1.10 | 0.80 | 0.77 | 1.19 | 1.04 | 0.95 | 1.00 | 0.95 |

### 8 Flow cytometry (undamaged HeLa cells): Gating and controls

Gating used for graphs shown in **Fig 2** and **Fig S3**:

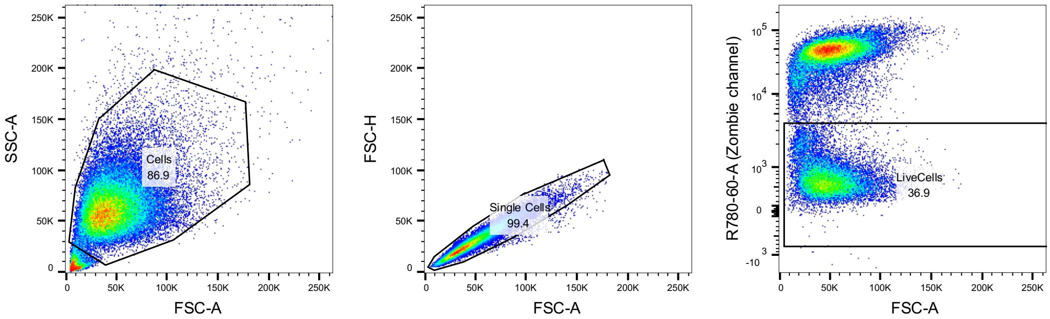

*gating from untreated Hela cells* *gating from untreated Hela cells from the "Cells" population* *gating from heat-shocked cells*

**Figure S13:** Gating and control samples for flow cytometry analysis of the probe and fluorophore permeability.

### 9 NMR Spectra

#### 5-Sulfo-2´,7´-dichlorofluorescein ($H_2$-$FS_1$)

##### $^1$H-NMR

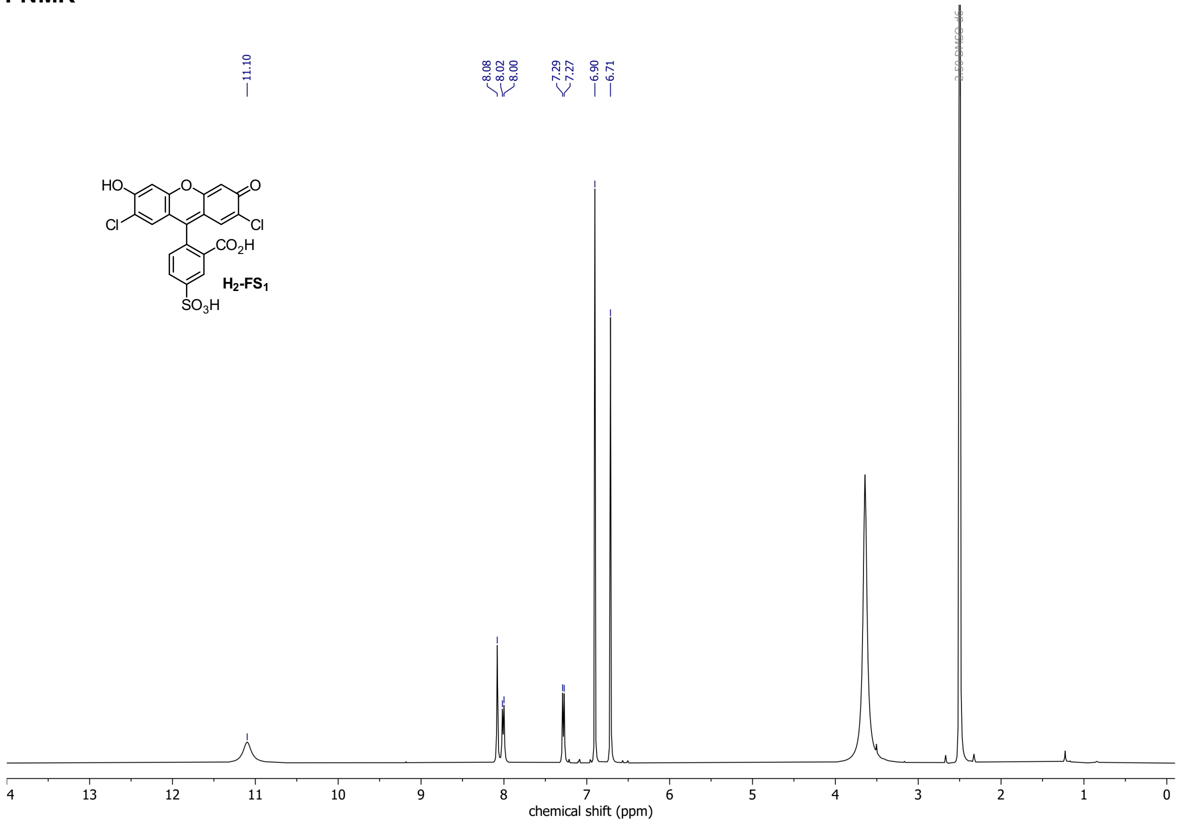

##### $^{13}$C-NMR

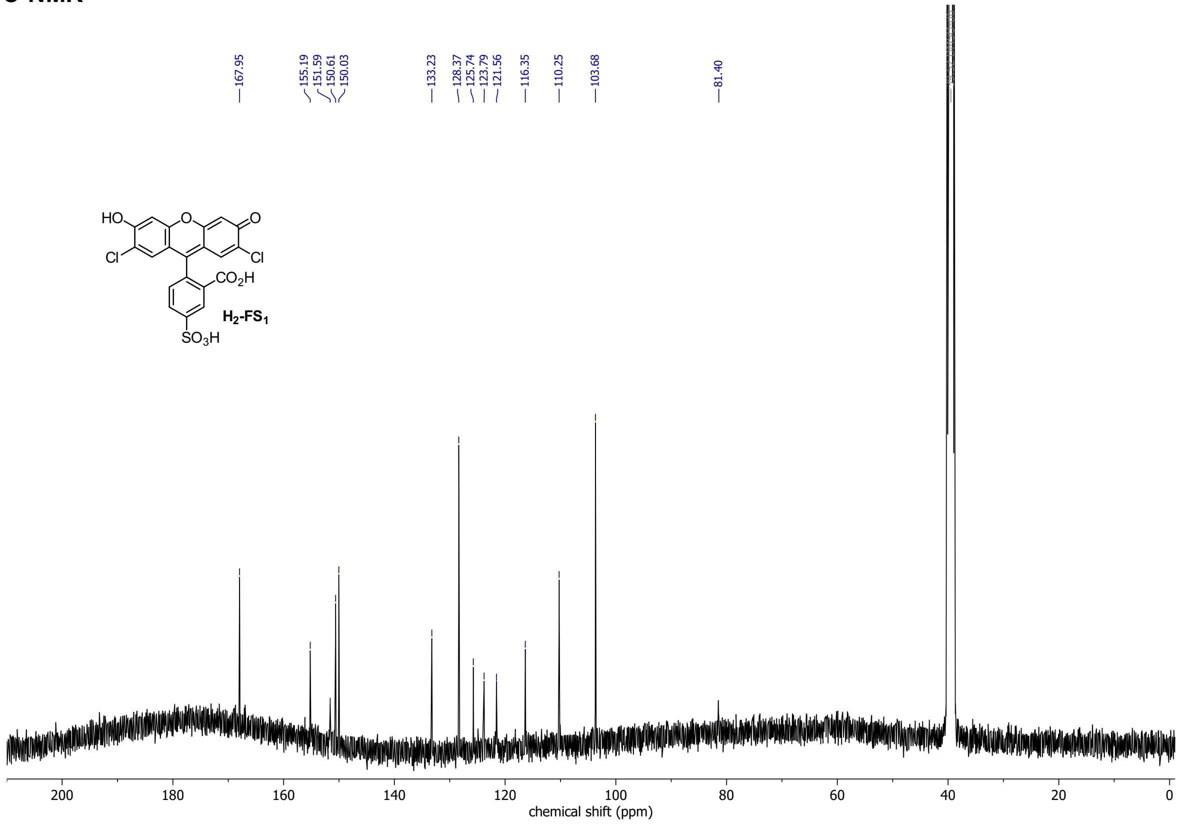

#### 5-Sulfo-2´,7´-dichlorofluorescein diacetate ($a_2$-$FS_1$)

##### $^1$H-NMR

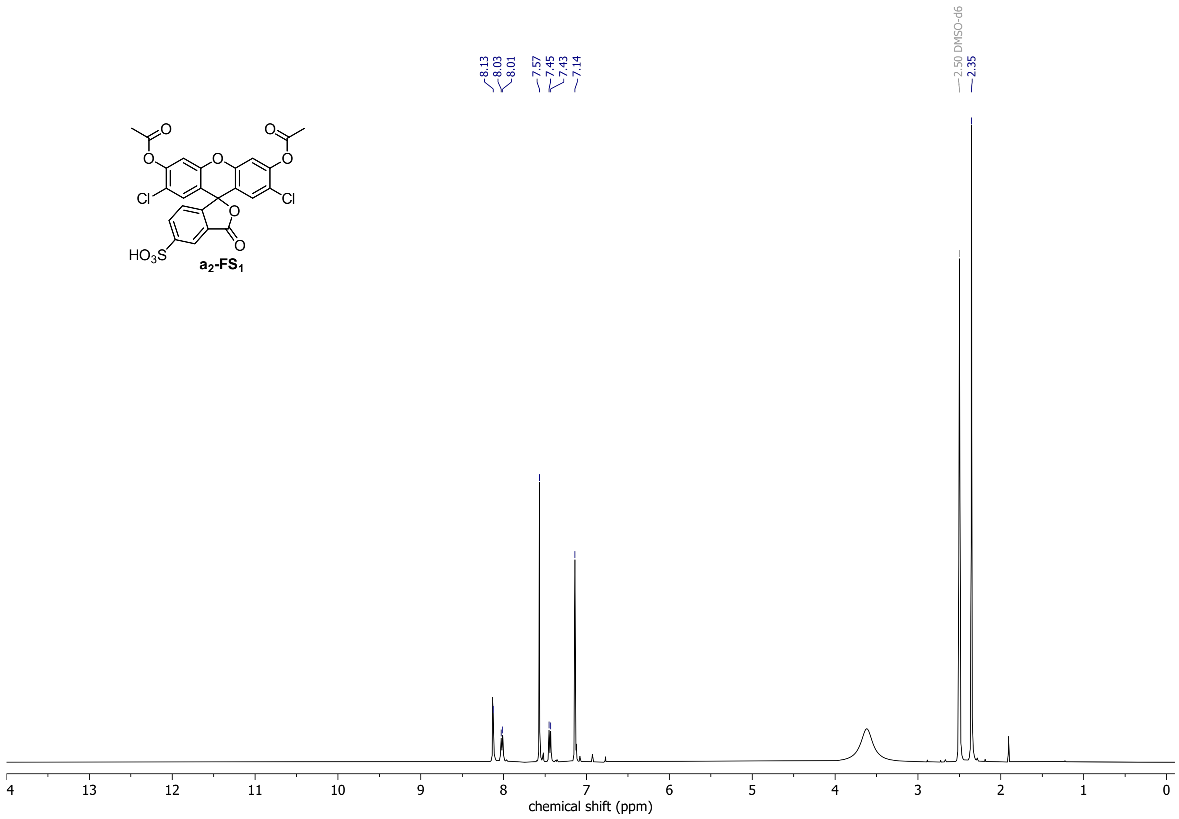

##### $^{13}$C-NMR

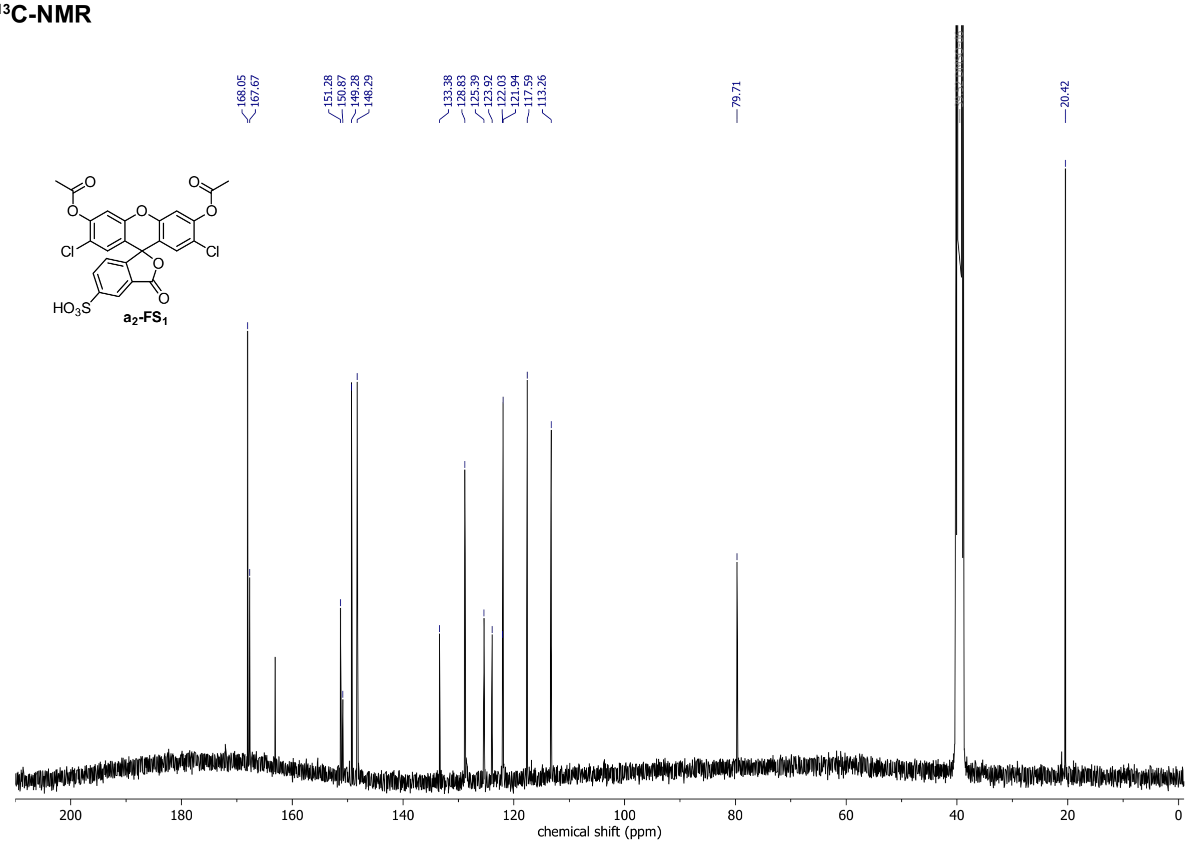

#### 5-Sulfo-2´,7´-dichlorofluorescein diisobutyrate ($i_2$-$FS_1$)

##### $^1$H-NMR

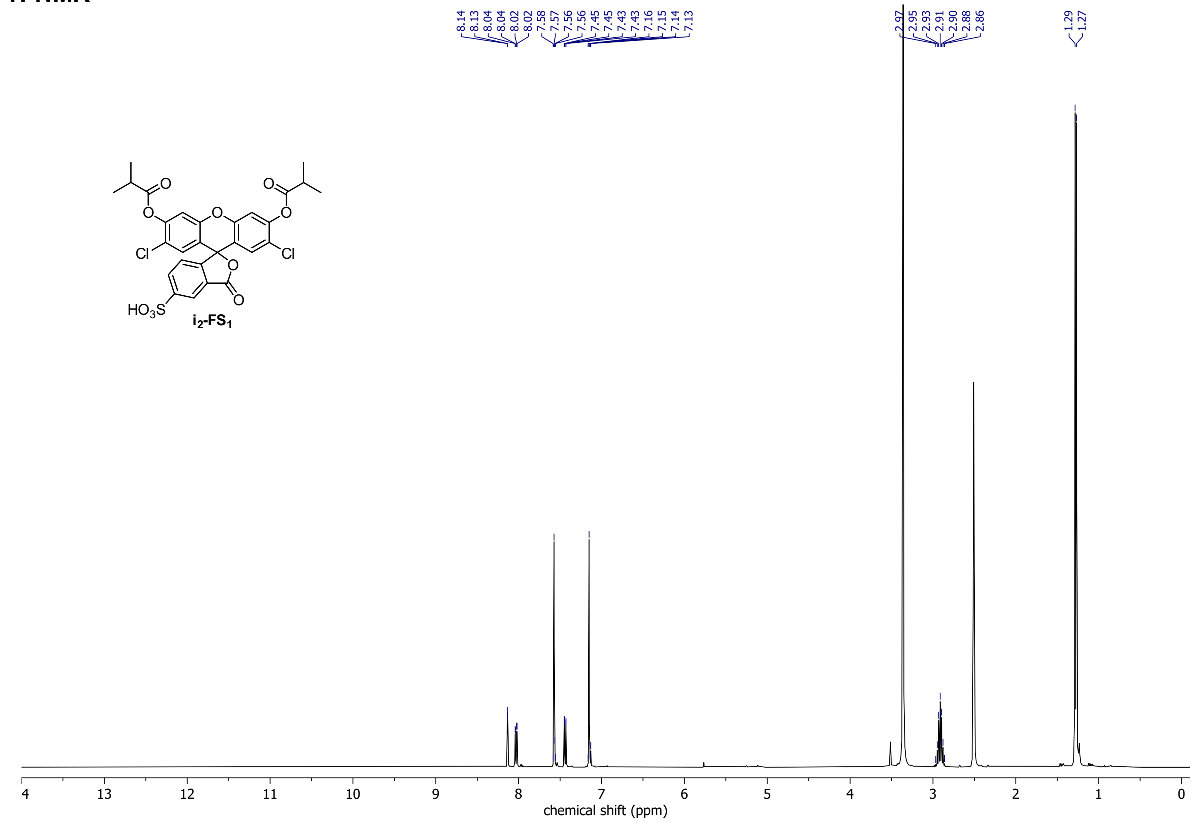

##### $^{13}$C-NMR

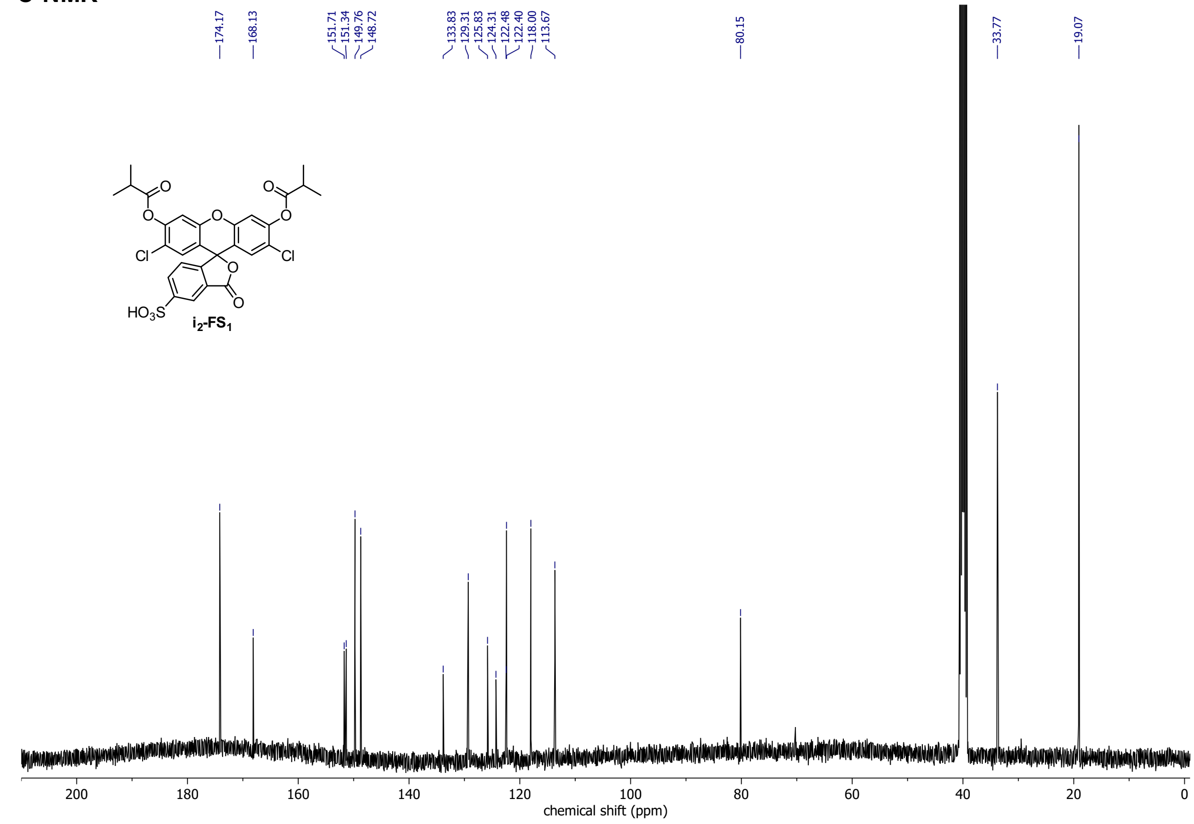

**iMe-FS$_1$**
**$^1$H-NMR**

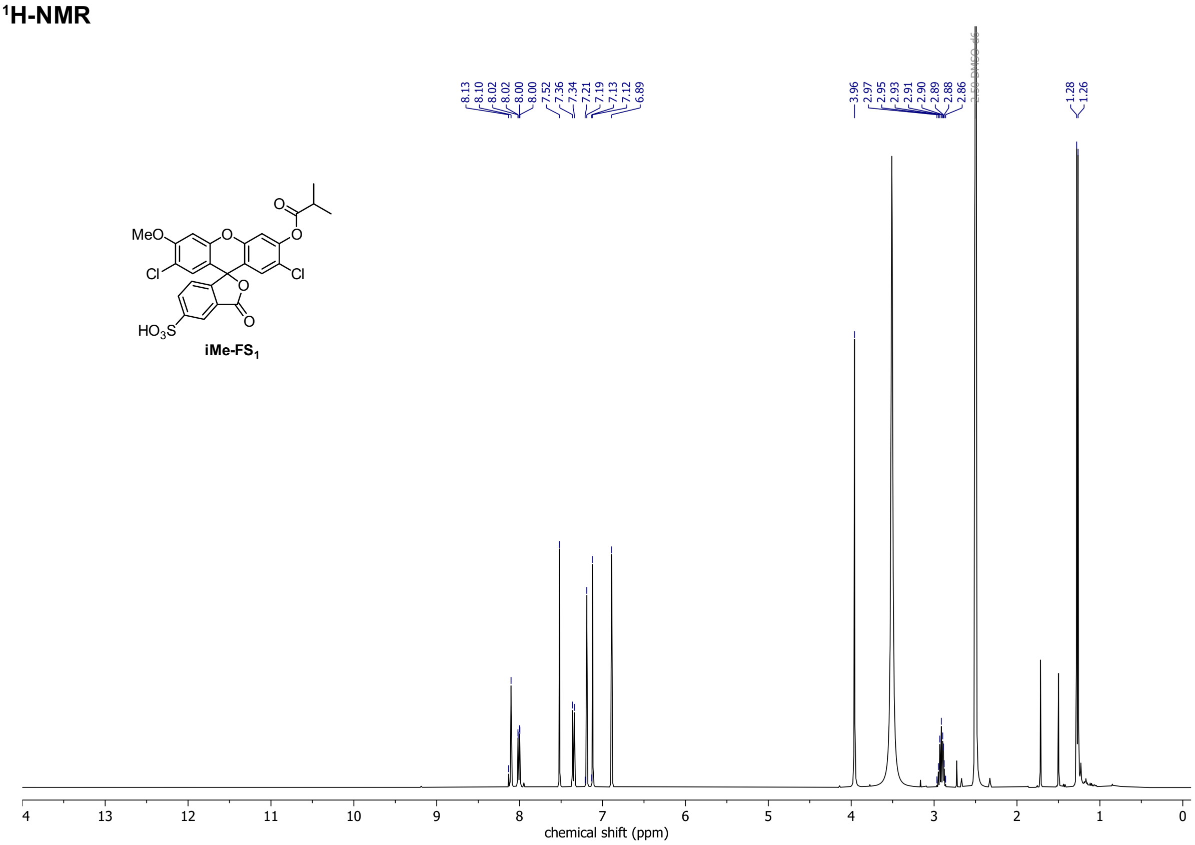

**$^{13}$C-NMR**

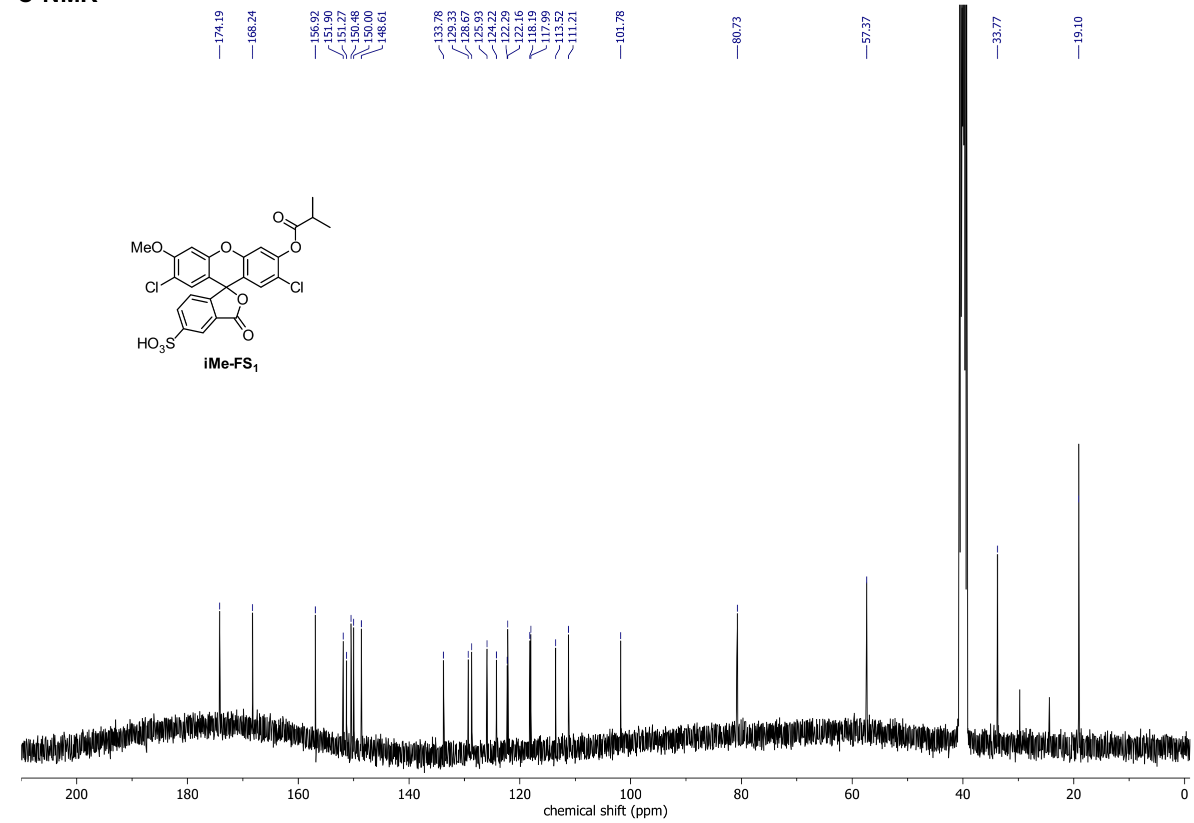

#### iEM-FS$_1$

##### $^1$H-NMR

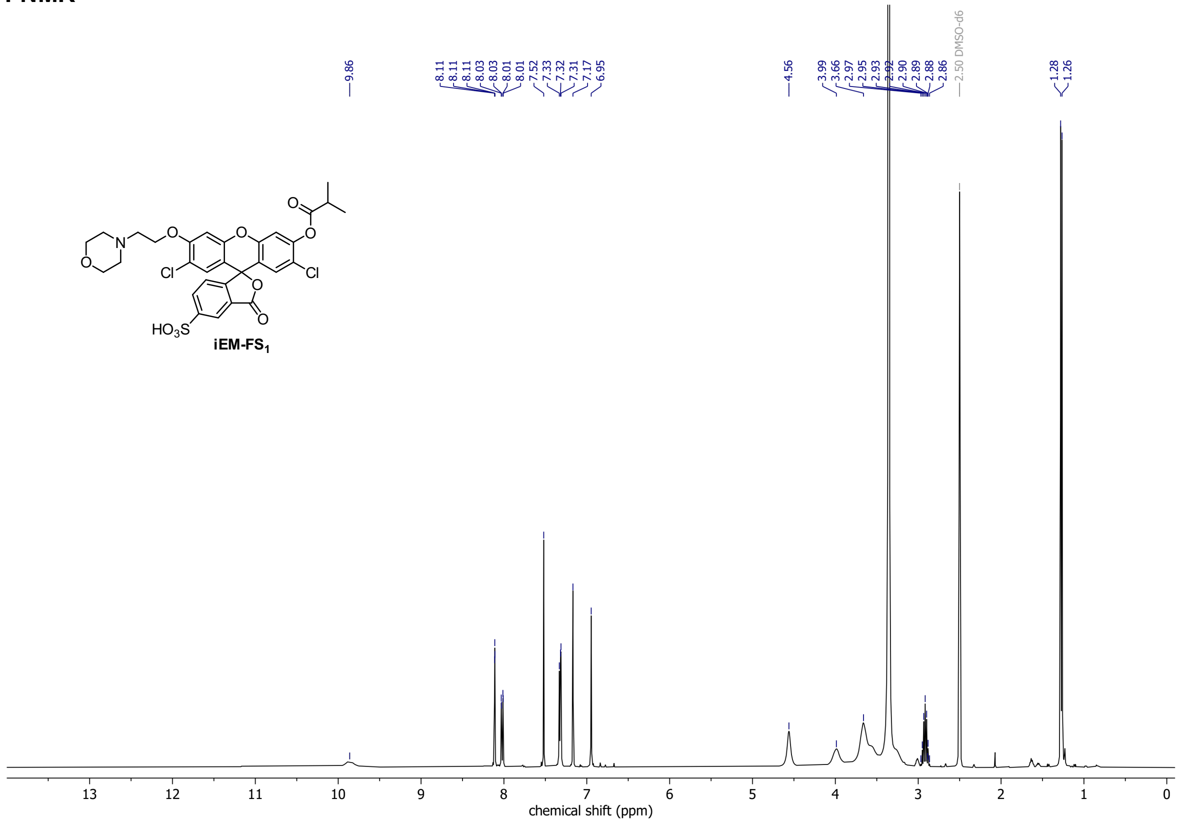

##### $^{13}$C-NMR

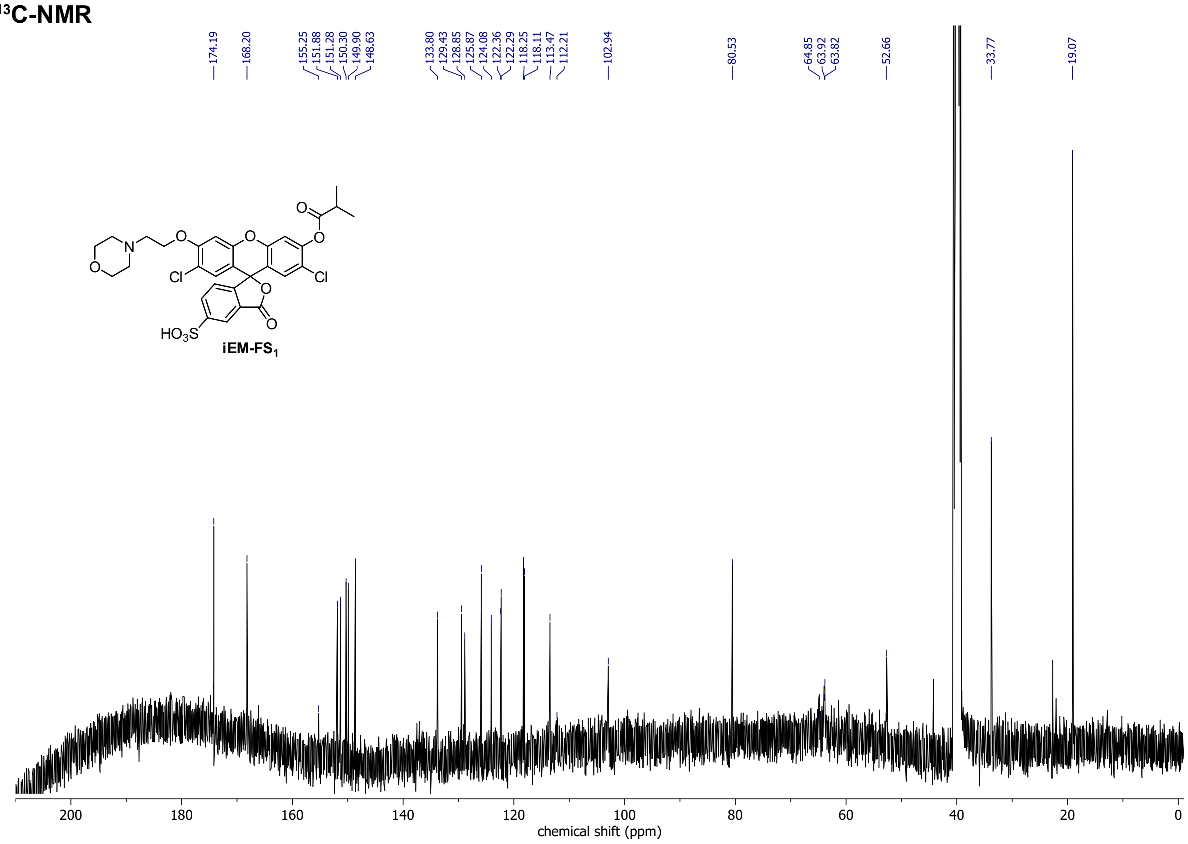

#### HSQC spectrum

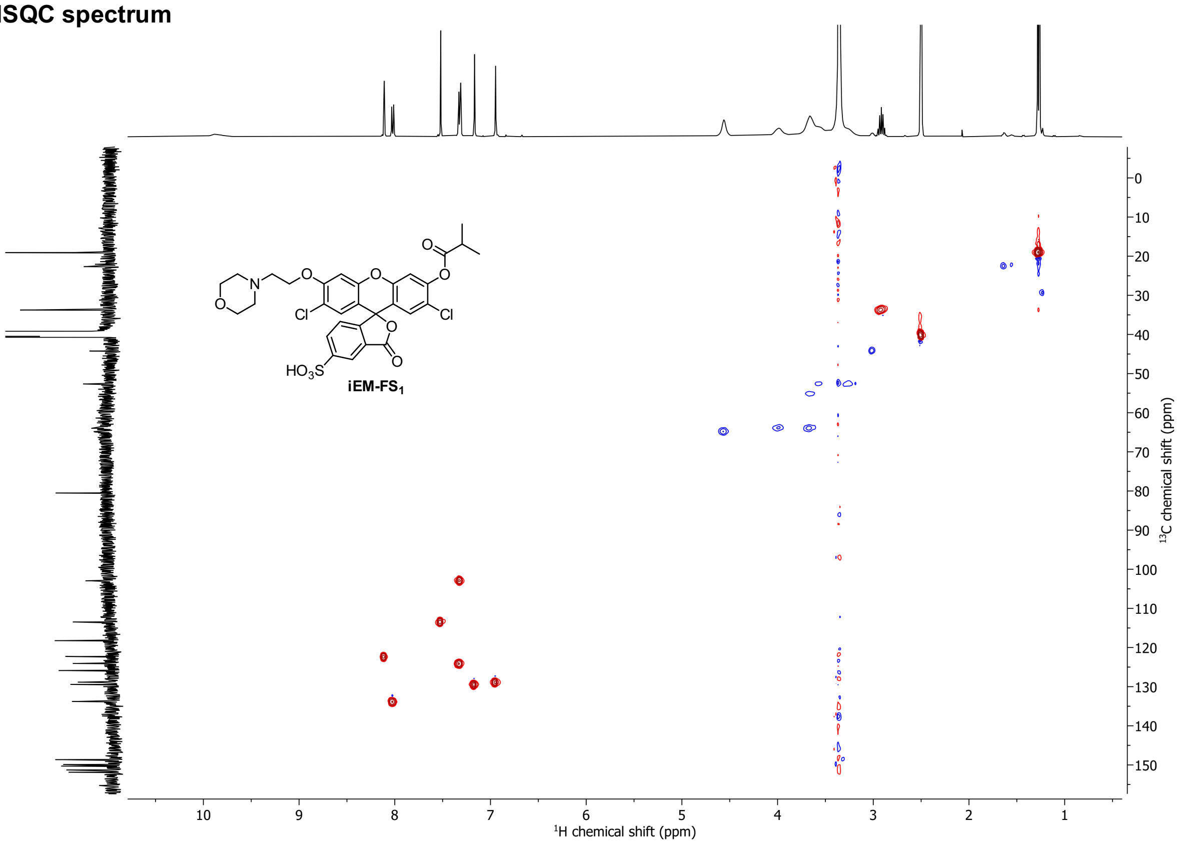

#### iPS-FS$_1$

##### $^1$H-NMR

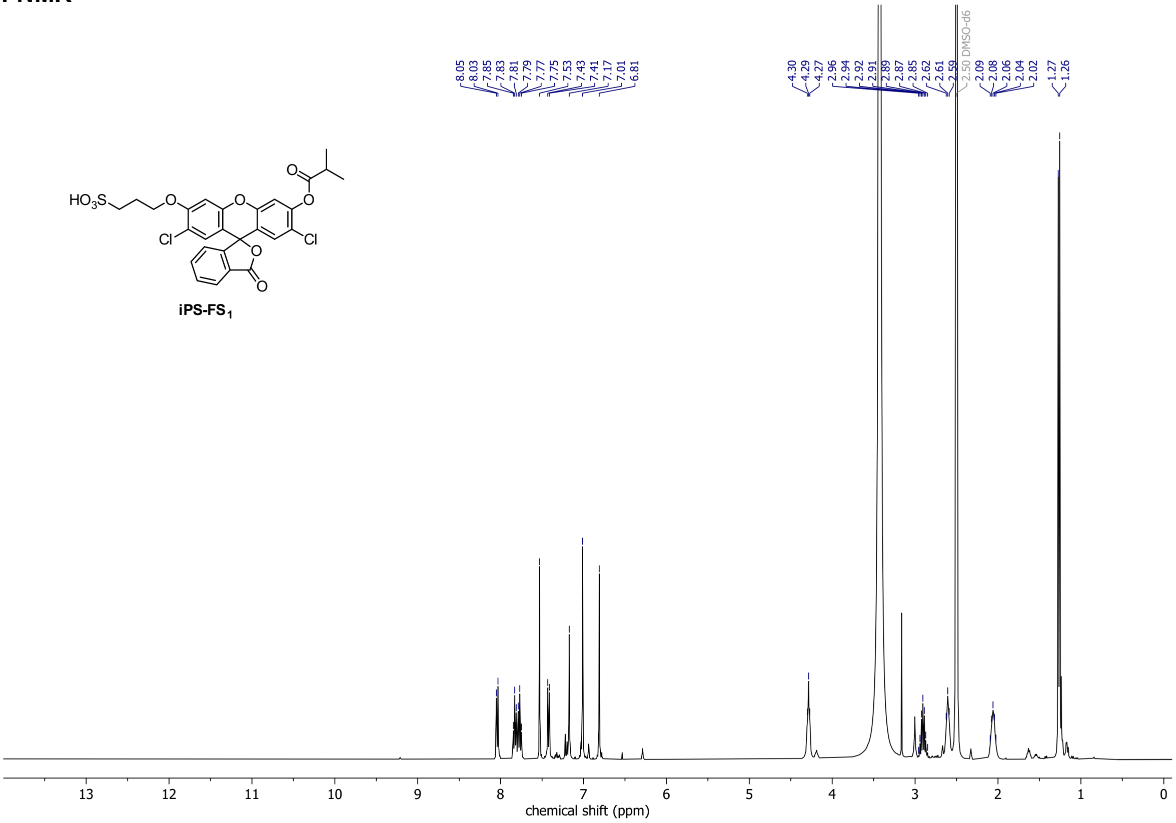

##### $^{13}$C-NMR

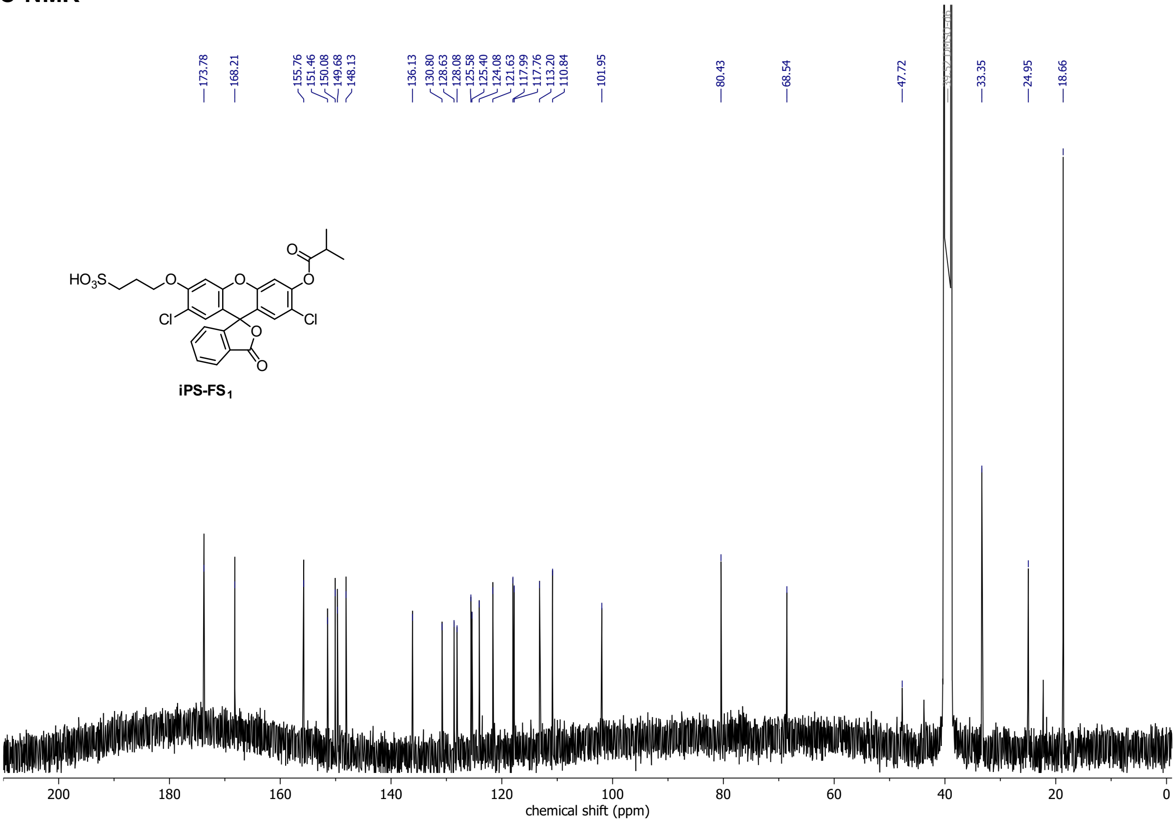

#### iPS-FS$_2$

##### $^1$H-NMR

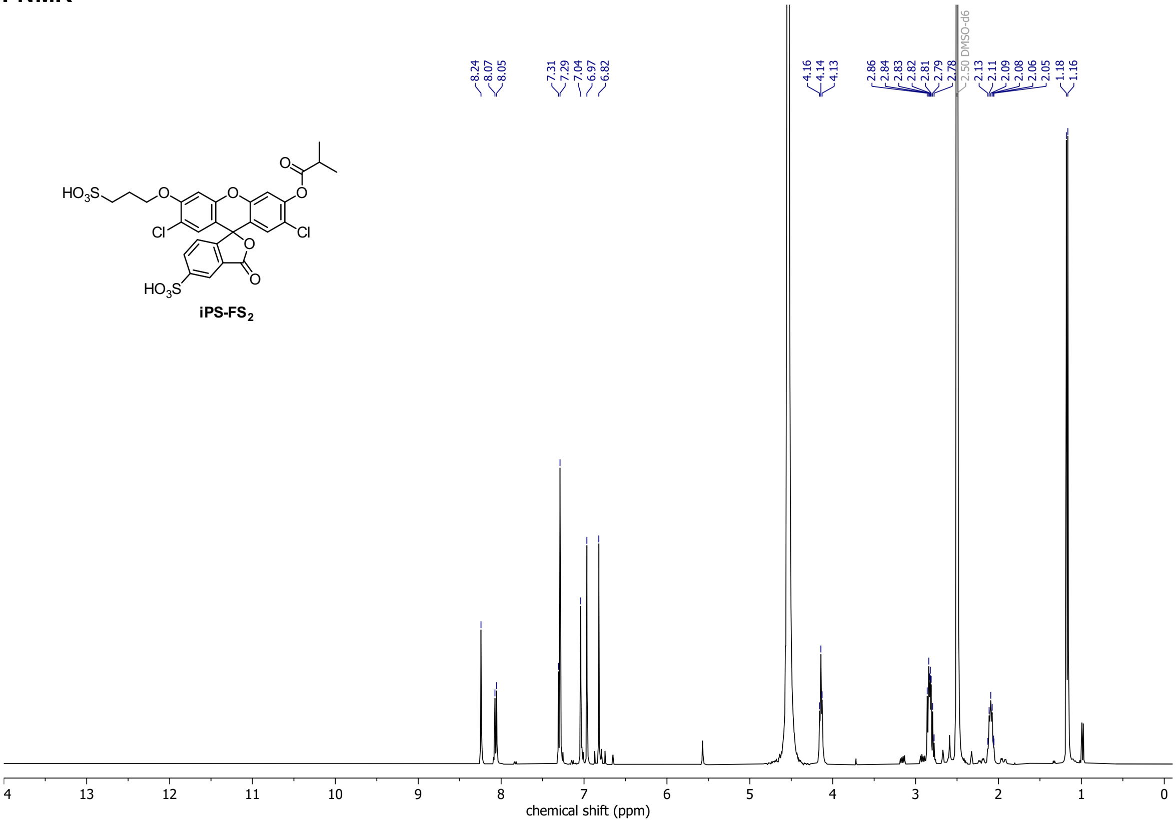

##### $^{13}$C-NMR

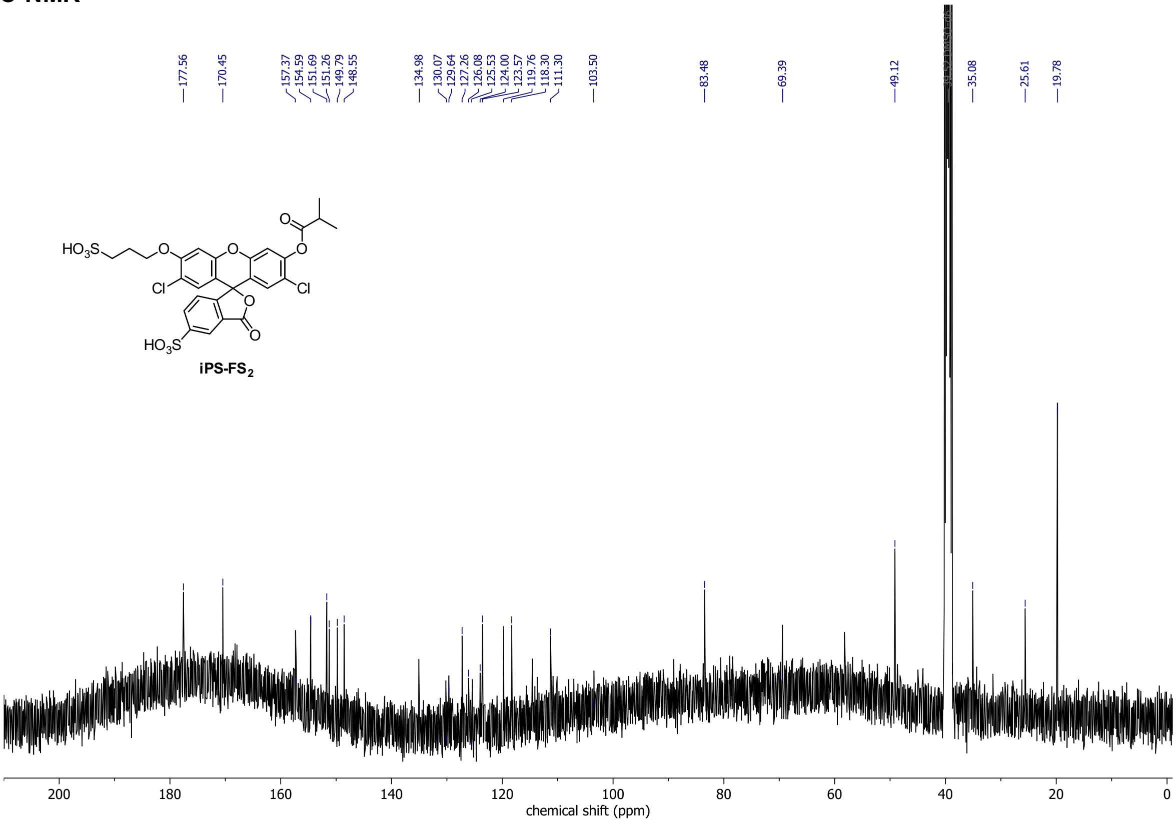

## H-Me-FS$_1$

##### $^1$H-NMR

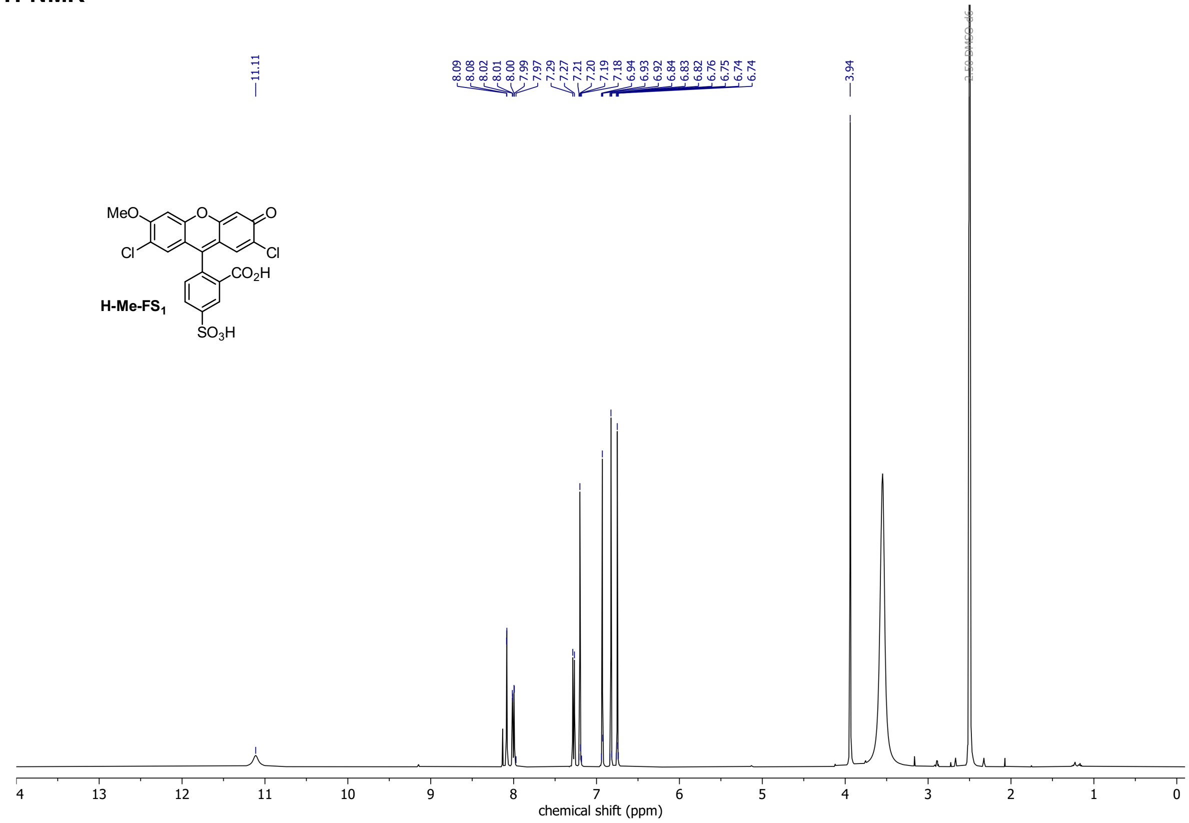

##### $^{13}$C-NMR

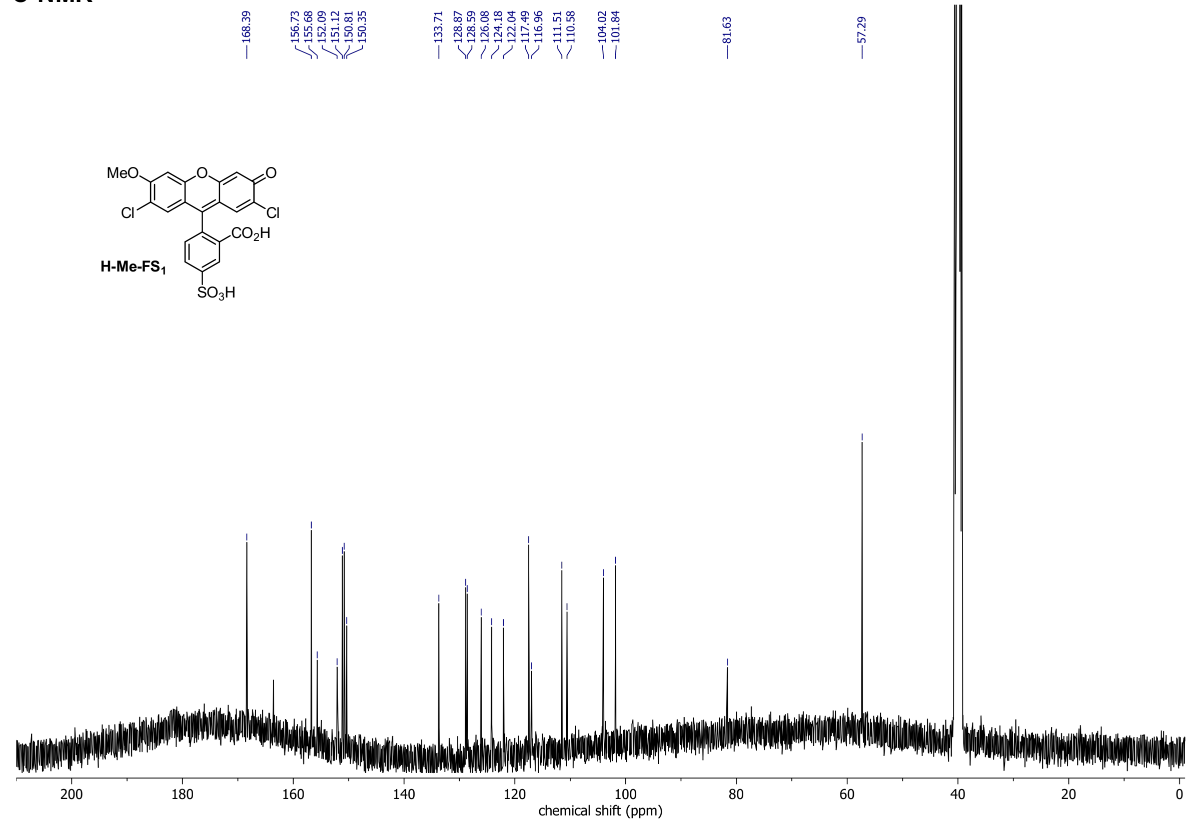

## H-EM-FS$_1$

##### $^1$H-NMR

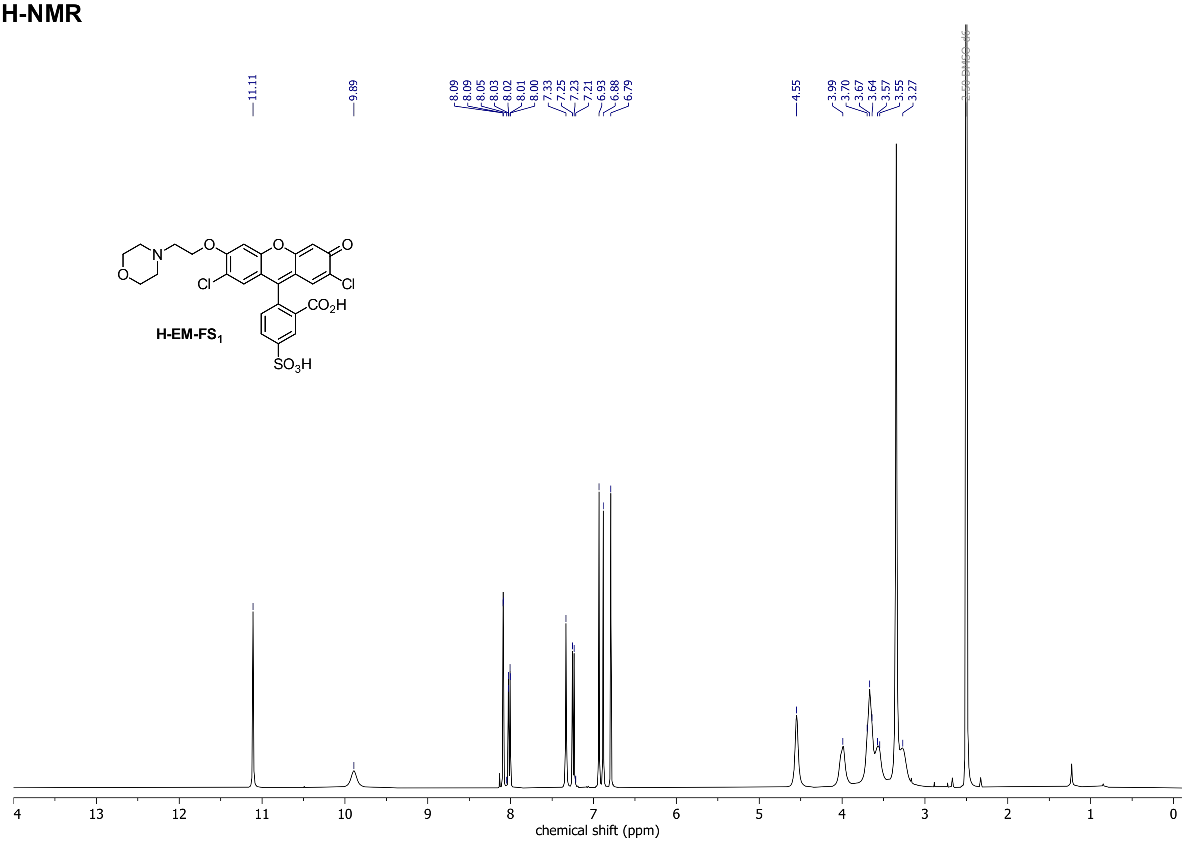

##### $^{13}$C-NMR

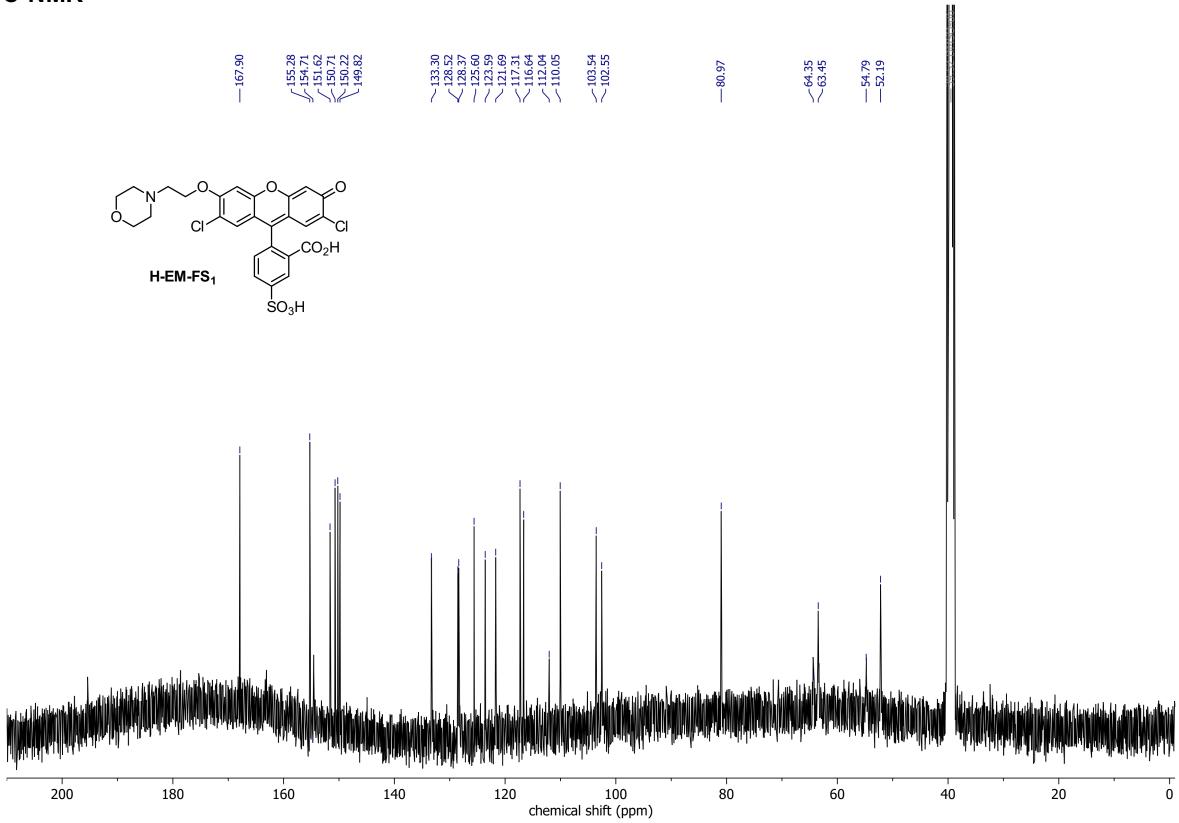

**HSQC spectrum**

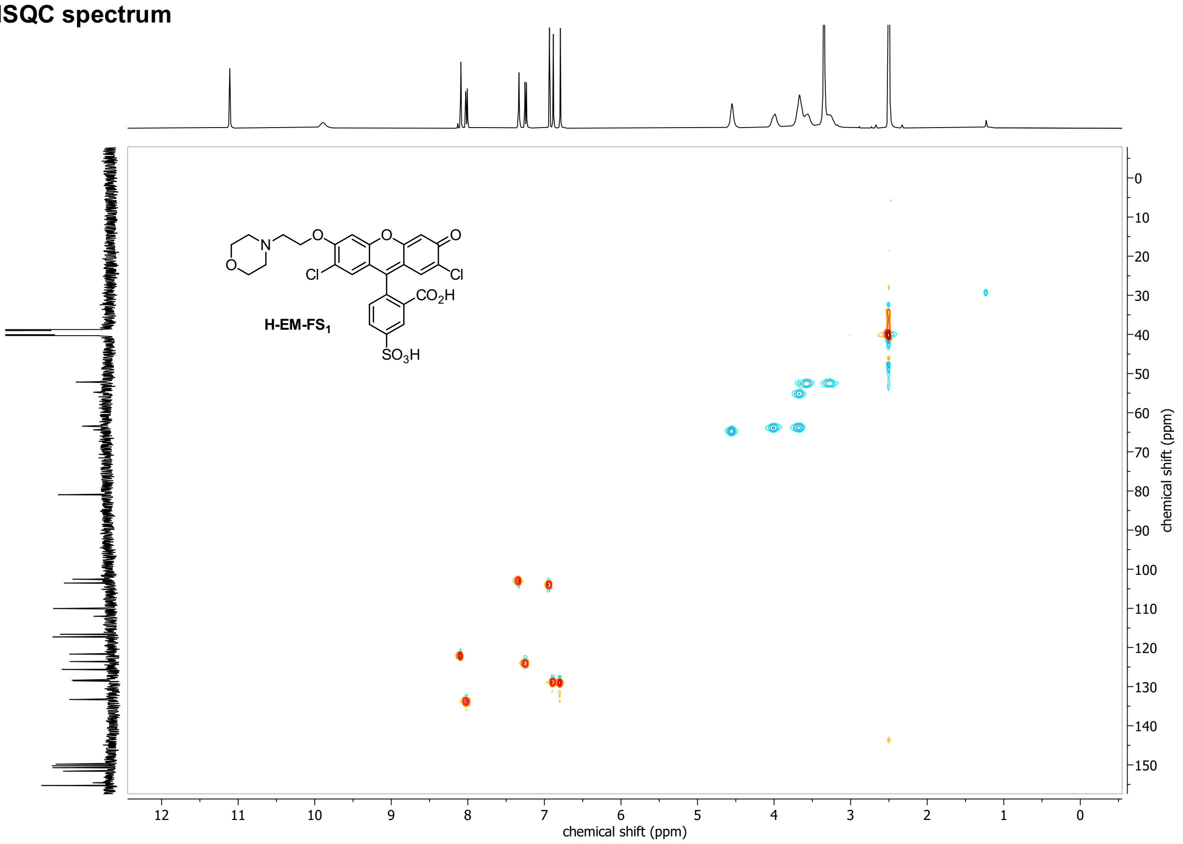

## H-PS-$FS_1$

##### $^1$H-NMR

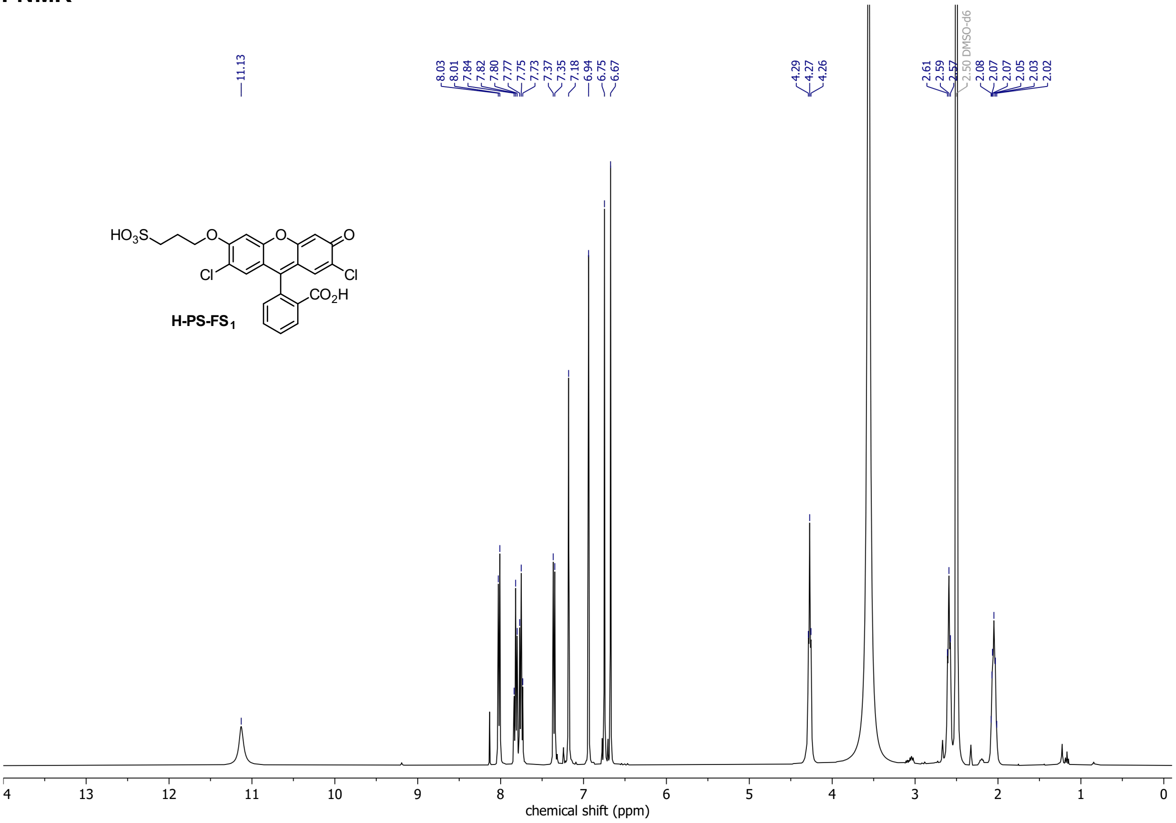

##### $^{13}$C-NMR

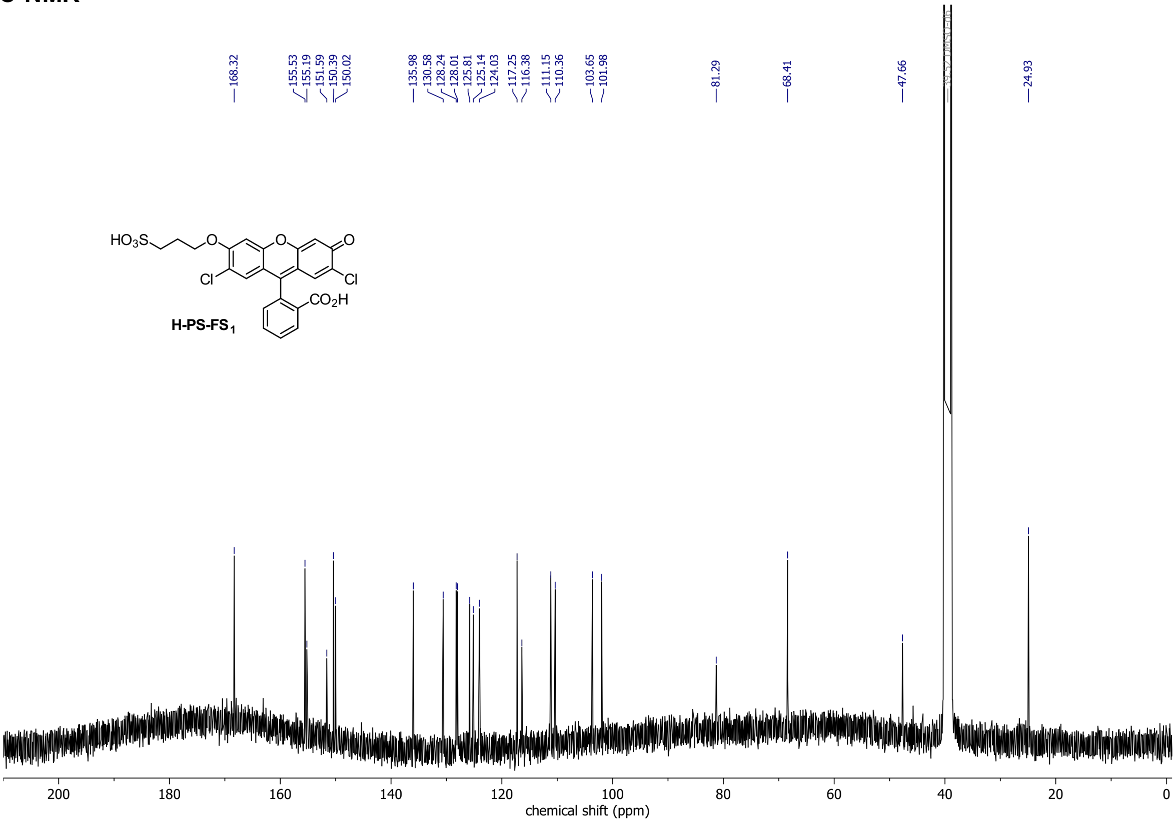

## H-PS-FS$_2$

##### $^1$H-NMR

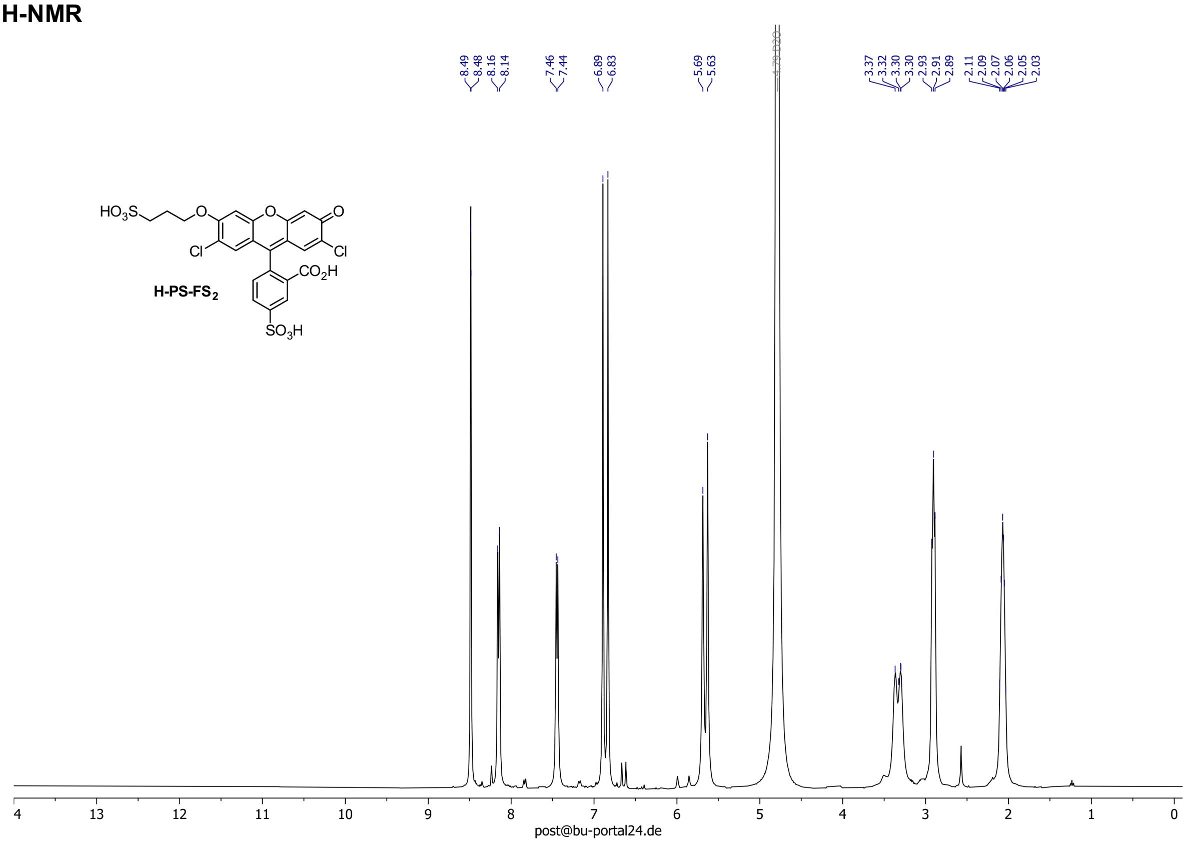

##### $^{13}$C-NMR

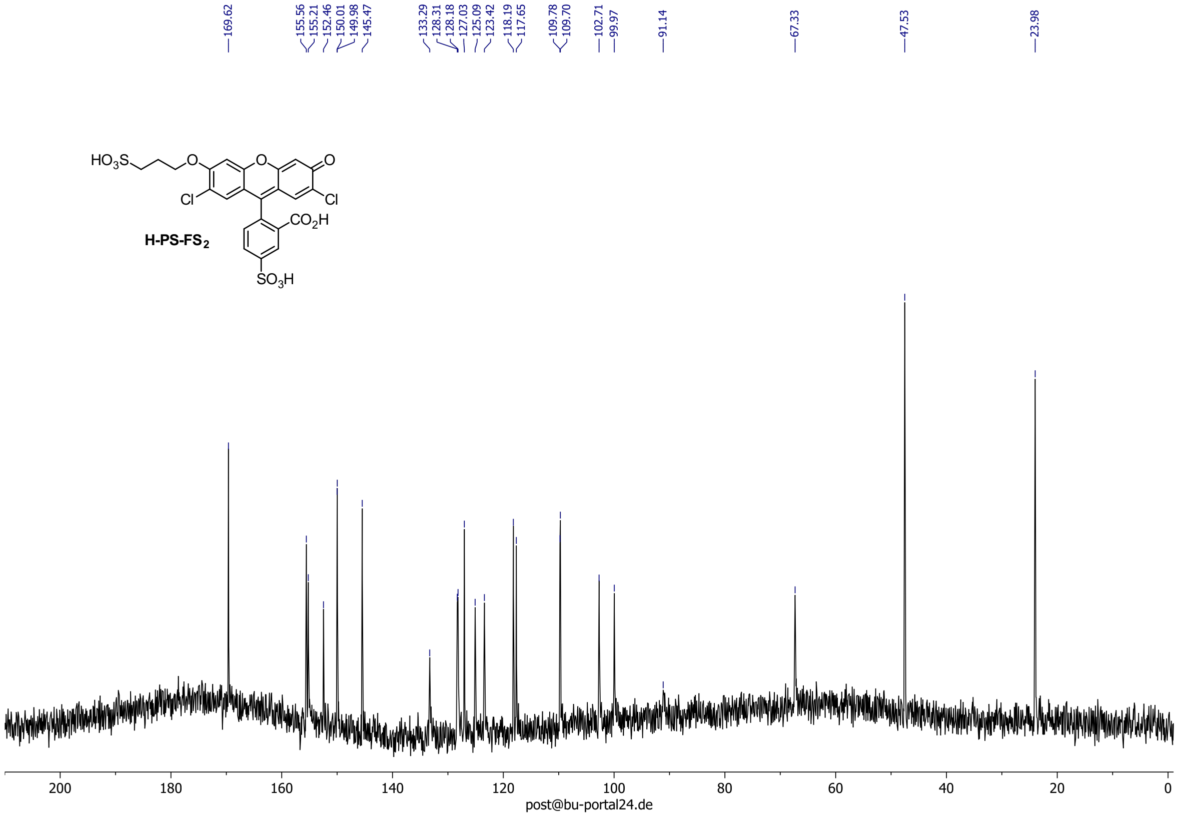

#### MSS00-PS-FS$_2$

##### $^{1}$H-NMR

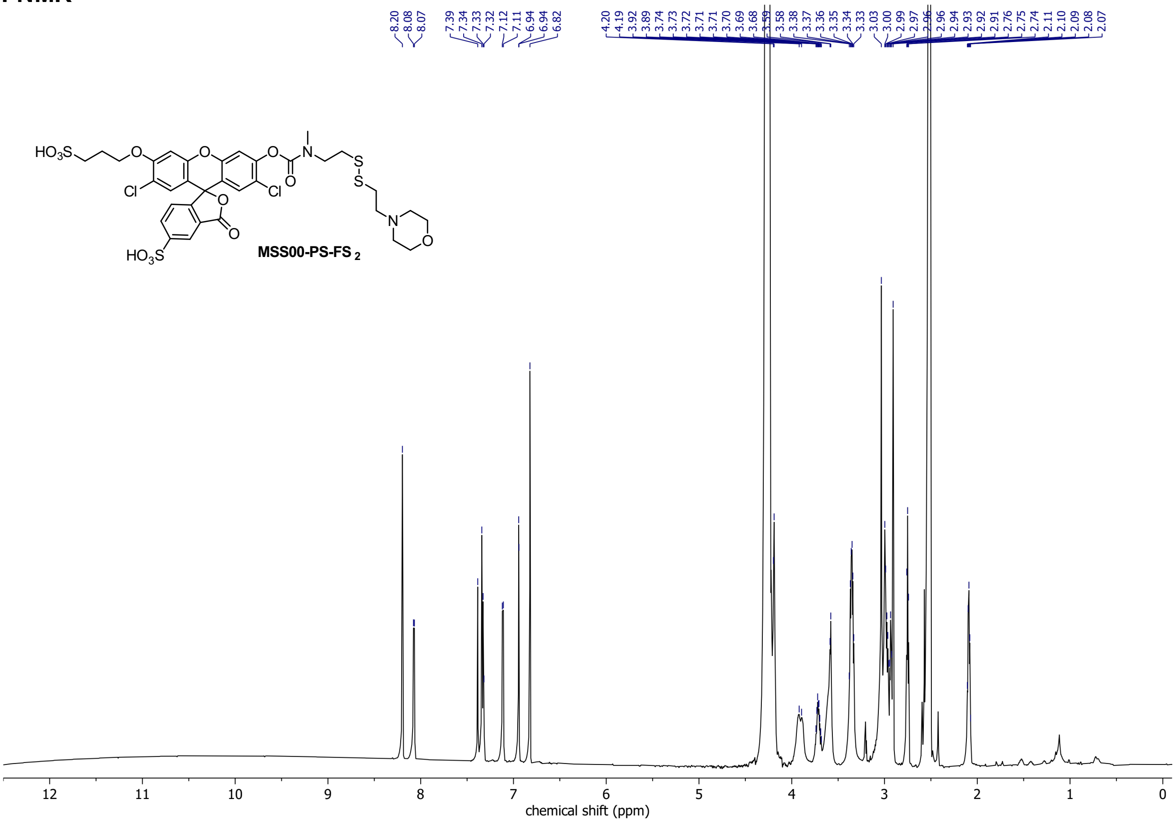

##### $^{13}$C-NMR

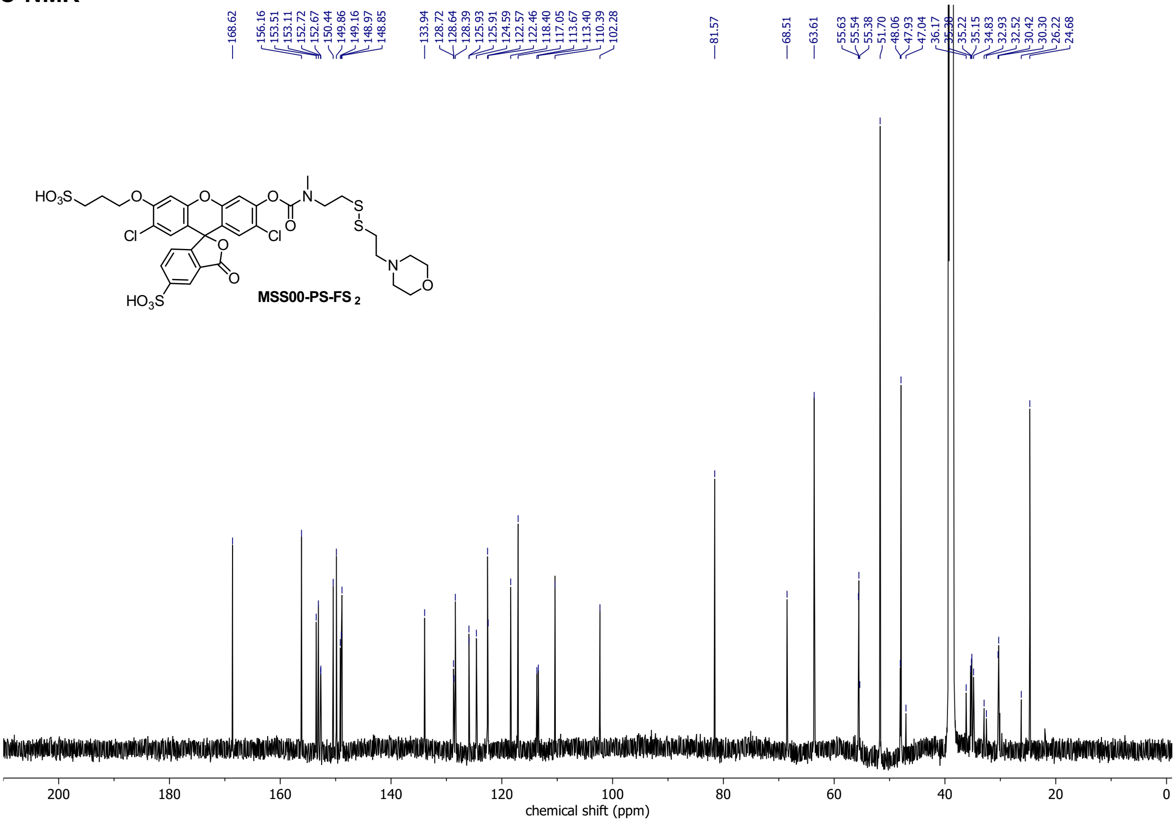

#### HSQC spectrum

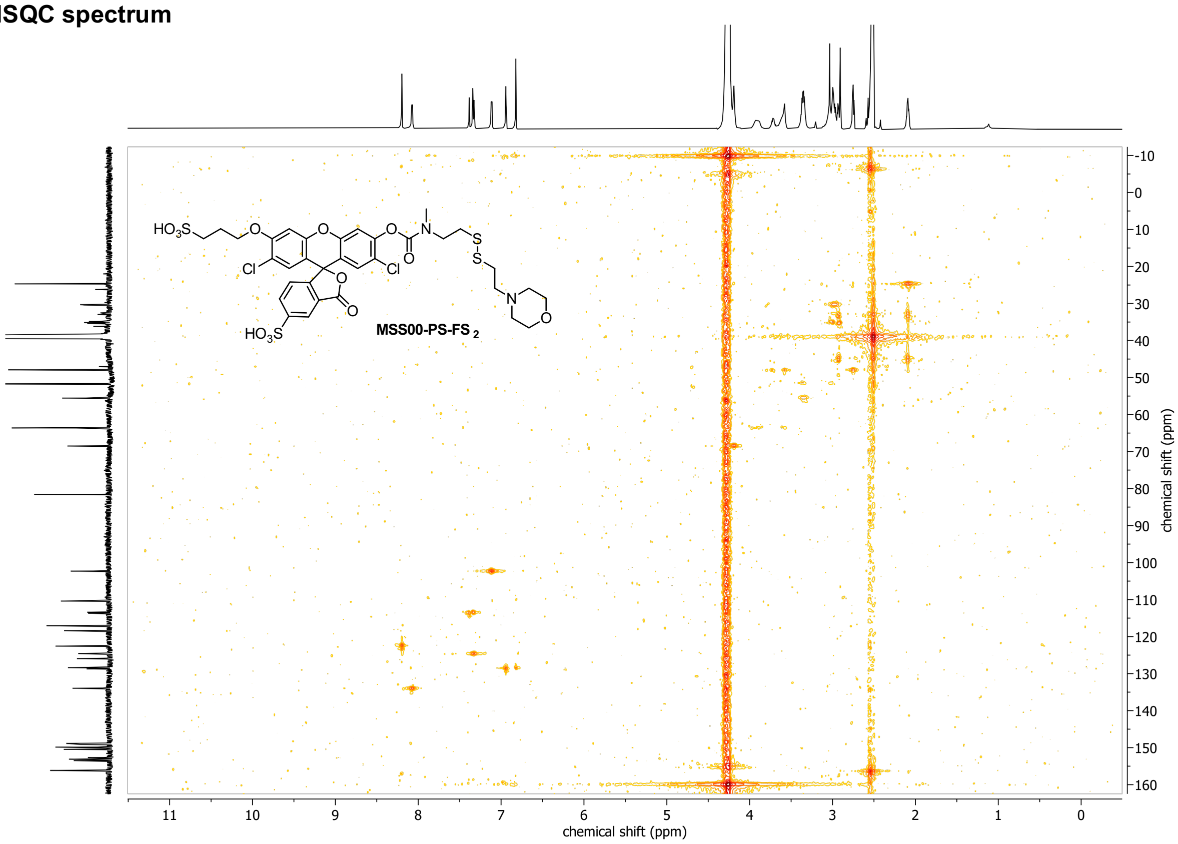

#### HSQC spectrum

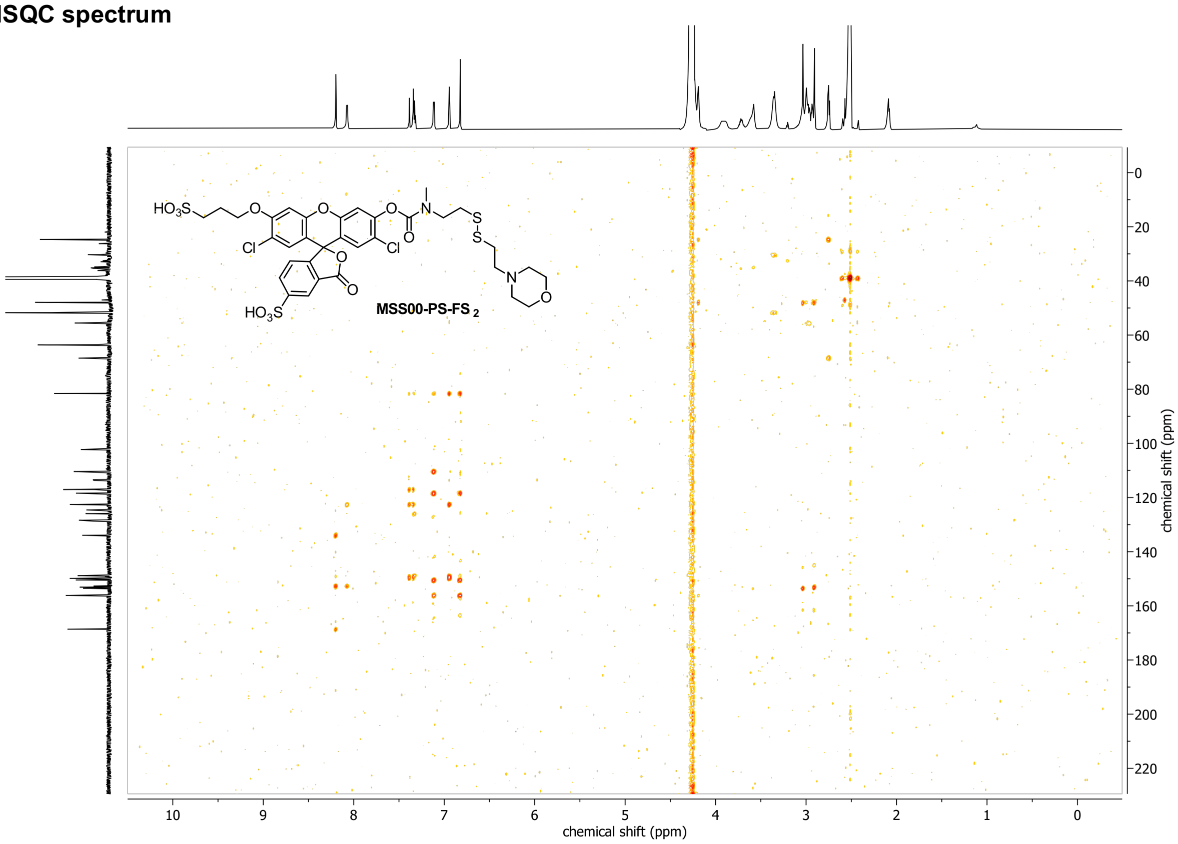

**Dichlorofluorescein isobutyrate NHS-ester (NHS-iFS$_0$)**

**$^1$H-NMR**

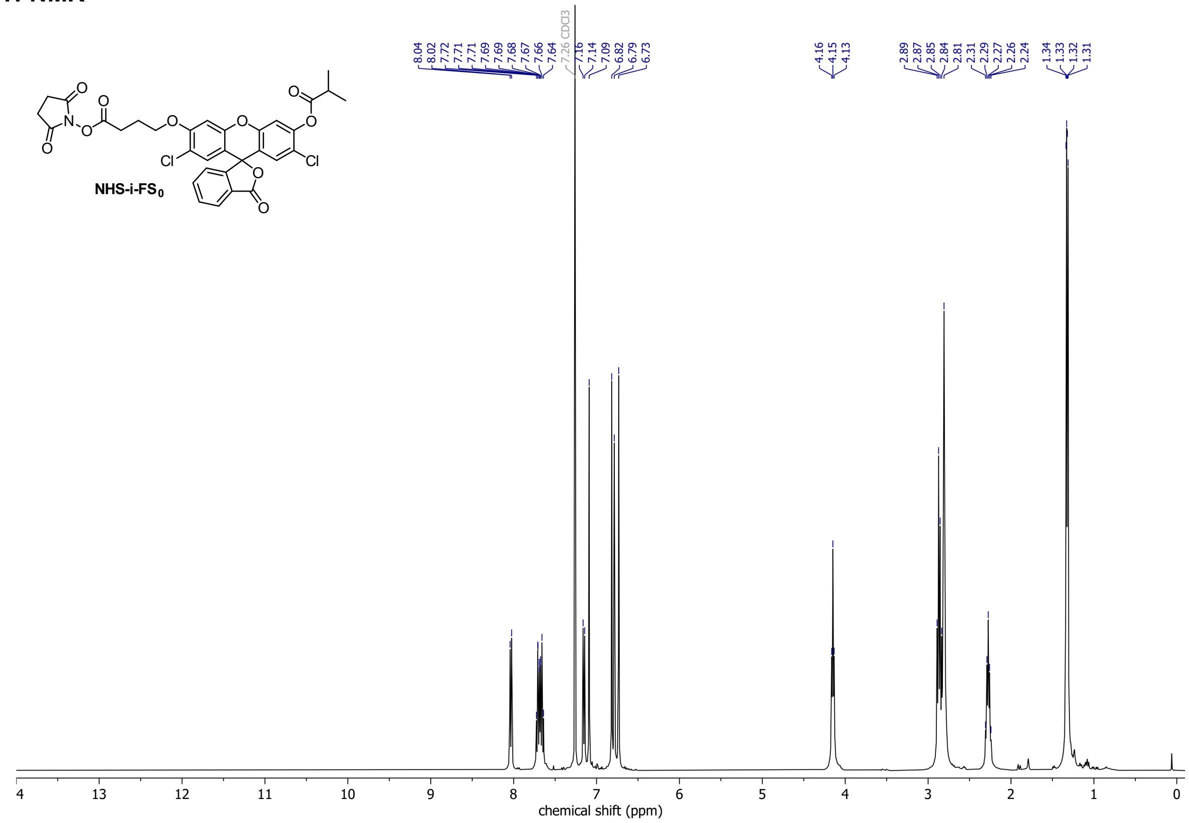

**$^{13}$C-NMR**

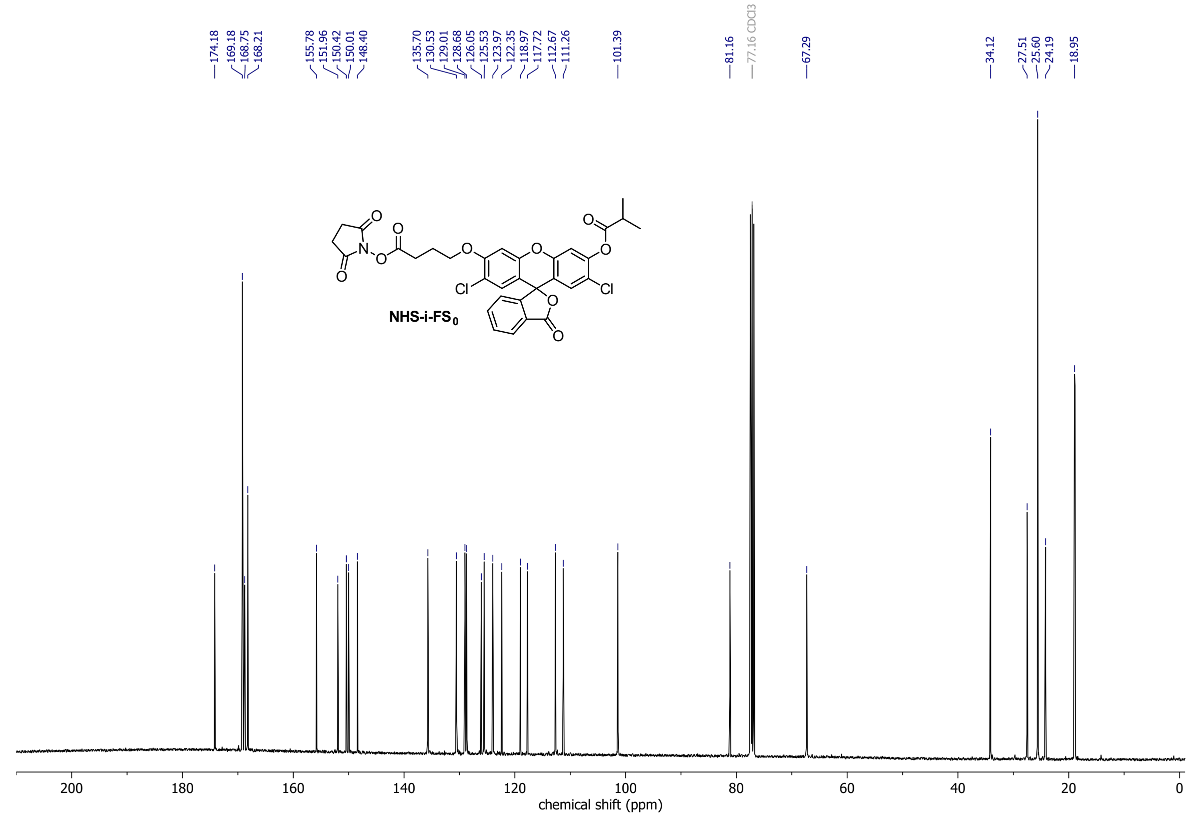
